# Human Saliva Modifies Growth, Biofilm Architecture and Competitive Behaviors of Oral Streptococci

**DOI:** 10.1101/2023.08.21.554151

**Authors:** Allen Choi, Kevin Dong, Emily Williams, Lindsey Pia, Jordan Batagower, Paige Bending, Iris Shin, Daniel I. Peters, Justin R. Kaspar

## Abstract

The bacteria within supragingival biofilms participate in complex exchanges with other microbes inhabiting the same niche. One example are the mutans group streptococci (*Streptococcus mutan*s), implicated in the development of tooth decay, and other health-associated commensal streptococci species. Previously, our group transcriptomically characterized intermicrobial interactions between *S. mutans* and several species of oral bacteria. However, these experiments were carried out in a medium that was absent of human saliva. To better mimic their natural environment, we first evaluated how inclusion of saliva affected growth and biofilm formation of eight streptococci species individually, and found saliva to positively benefit growth rates while negatively influencing biomass accumulation and altering spatial arrangement. These results carried over during evaluation of 29 saliva-derived isolates of various species. Surprisingly, we also found that addition of saliva increased the competitive behaviors of *S. mutans* in coculture competitions against commensal streptococci that led to increases in biofilm microcolony volumes. Through transcriptomically characterizing mono- and cocultures of *S. mutans* and *Streptococcus oralis* with and without saliva, we determined that each species developed a nutritional niche under mixed-species growth, with *S. mutans* upregulating carbohydrate uptake and utilization pathways while *S. oralis* upregulated genome features related to peptide uptake and glycan foraging. *S. mutans* also upregulated genes involved in oxidative stress tolerance, particularly manganese uptake, which we could artificially manipulate by supplementing in manganese to give it an advantage over its opponent. Our report highlights observable changes in microbial behaviors via leveraging environmental- and host-supplied resources over their competitors.

## INTRODUCTION

The bacteria that colonize the oral cavity live in complex communities within biofilms attached to the tooth’s surface. An array of complex intermicrobial interactions are present, with specific interactions developing to be synergistic, antagonistic or neutral between species (1, 2). One well studied antagonistic interaction occurs between mutans group streptococci (*Streptococcus mutans, Streptococcus sobrinus*) and oral commensal streptococci (*Streptococcus* sp. A12, *Streptococcus cristatus, Streptococcus gordonii, Streptococcus mitis, Streptococcus oralis, Streptococcus sanguinis*) (3, 4). Mutans group streptococci form microcolony structures that are encased and held structurally sound by glucan polysaccharides they abundantly produce (5–7). These species are also highly acidogenic, producing organic acids that are pooled within their microcolonies from fermentation of dietary carbohydrates, causing localized areas of acidic pH leading to demineralization of the tooth’s enamel (8, 9). Oral commensal streptococci are commonly acid sensitive and encode various defense mechanisms, such as hydrogen peroxide production (10, 11) and arginine metabolism via the arginine deiminase (ADS) pathway (12, 13) to combat the outgrowth of the mutans group streptococci as well as to buffer against glycolytic acids. One favorable strategy towards tooth decay (dental caries) prevention is thwarting the emergence of mutans group streptococci within the oral microbiome while keeping the normal, protective flora intact. Development of new therapeutic approaches relies on fully characterizing these exchanges between competing microbes and unraveling the strategies employed by each species that leads to a gain in competitive advantage over the other.

Our group recently transcriptomically characterized coculture growth between *S. mutans* and several species of commensal streptococci (14). However, these experiments were carried out in our lab-based experimental medium, tryptone and yeast extract (TY-). *In vivo*, supragingival biofilms are constantly bathed in saliva. Resting human saliva contains organic and inorganic ions, peptides, and over 400 different host-derived proteins, many of which are glycosylated (15, 16). Individual free amino acids are present at concentrations less than 12 mg L^-1^ and the preferred carbohydrate glucose at a concentration less than 100 µM (17, 18). During periods of host fasting, oral microbes must obtain carbon, nitrogen and energy from sources such as glycoproteins, requiring an assortment of enzymes that release oligosaccharides, peptides and amino acids. These include several different classes of glycosidases and peptidases that are abundant throughout the genomes of oral bacteria, but are often distributed between different species that leads to a consortium of different strains required to cooperatively degrade and liberate carbohydrate and peptide/amino acid moieties (19–21). For example, the glycan foraging activities of *S. oralis* are sufficient to digest the *N*-linked glycans of plasma human α_1_-acid glycoprotein (22), yet lack several glycoside hydrolases encoded by *S. mitis* and peptidases of *S. gordonii* to degrade proline-rich proteins (PRPs) (21, 23). Of note, the genome of the caries pathogen *S. mutans* does not contain many, if any glycoside hydrolases as well as lacks PRP degradation activity (21, 23).

To understand whether competitive behaviors are altered by culturing these species within a medium that more closely mimics their natural environment, we evaluated cocultures of *S. mutans* and *S. oralis* in a TY- / human saliva mix that was optimally chosen based on our initial characterization of individual oral streptococci growth in medium mixes containing saliva. Our results show that inclusion of saliva enhances the competition of *S. mutans* against commensal streptococci through upregulation of carbohydrate uptake and glycolytic pathways as well as oxidative stress tolerance mechanisms. In turn, commensals express polysaccharide utilization loci (PULs) for the foraging of salivary glycans, segregating into their own nutritional niche. Our work begins to describe how each of these competing species leverages environmental- and host-supplied resources over the other, and further documents the critical role of oxidative stress tolerance and metals acquisition in the physiology of the caries pathogen *S. mutans*.

## RESULTS

### Growth of oral streptococci are enhanced with human saliva

To examine if/how the growth of oral streptococci are altered in human saliva, we first tested whether eight different strains could grow in only human saliva and/or human saliva supplemented with glucose as a carbohydrate source, consisting of six commensal species and two mutans group species **(Figure S1)**. None of the eight strains had measurable growth in pure human saliva alone, while each strain, except for *S*. sp. A12, displayed low to moderate growth in the saliva containing glucose.

Due to the marginal growth recorded for each strain after a 24 h monitoring period, an alternative strategy was chosen to mix our lab-based experimental medium, tryptone and yeast extract (TY-), half and half with either water (TY-Water, control) or human saliva (TY-Saliva) and observe the growth of the same eight species panel **(Figure 1A-H)**. The same amount of carbohydrate (20 mM glucose) was added at the beginning of each experiment into each medium. With this methodology, six of the eight strains had final yields (optical density at 600 nm) at or even higher than TY-, and all eight had yields greater than 0.4 in medium containing saliva. Additionally, five of the eight strains showed improved growth in the TY-Saliva condition through either faster doubling times and/or shorter lag times (measured as time taken for the culture to breach the OD_600 nm_ = 0.1 threshold). We also tested three other non-streptococci species including *Actinomyces oris, Corynebacterium matruchotii* and *Lactobacillus casei* **(Figure S2)**. None of these three species showed improved growth in TY-Saliva compared to TY-alone, and *A. oris* was not able to grow in 100% human saliva even with the addition of glucose.

**Figure 1.**
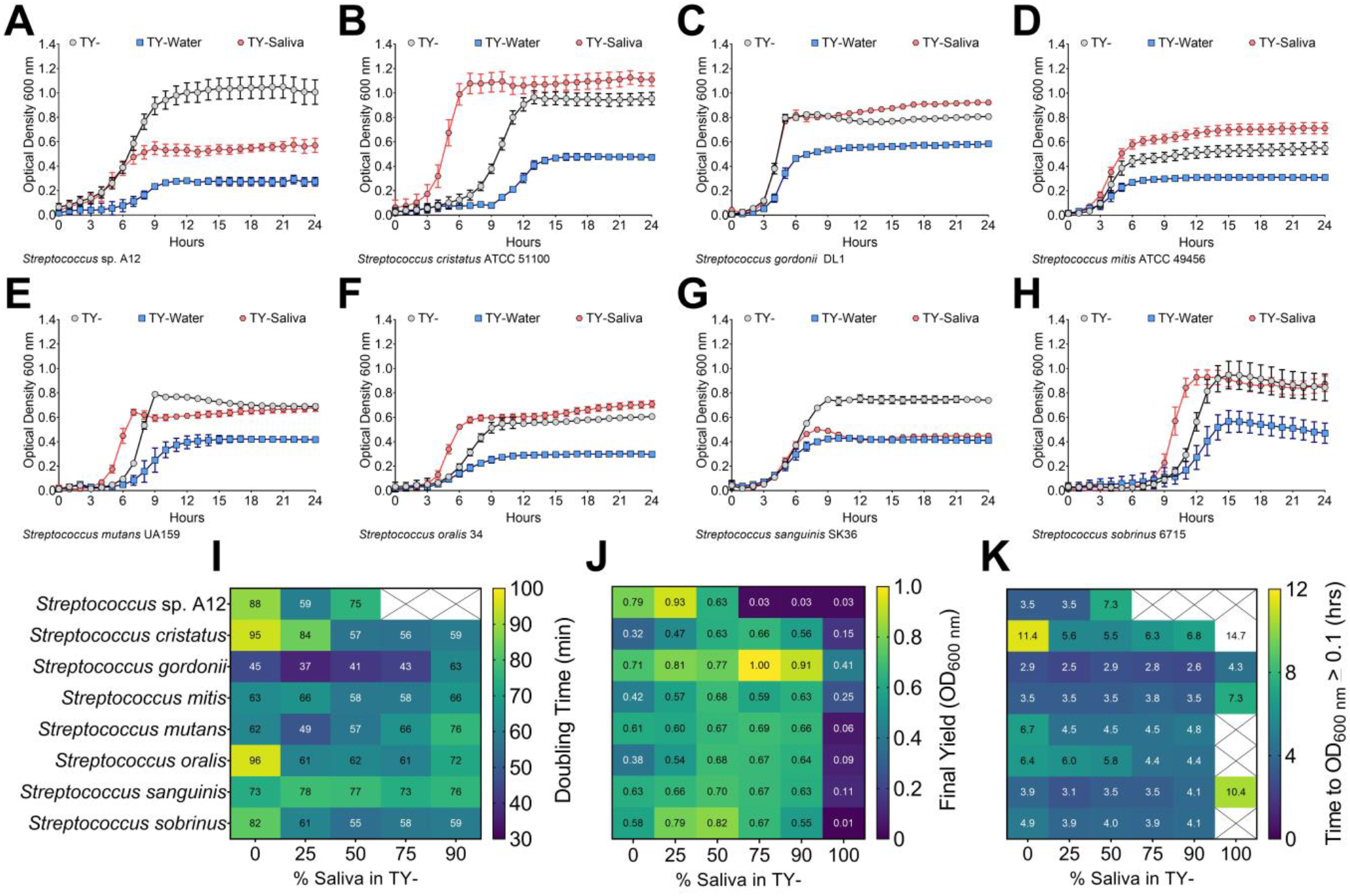
Growth of oral streptococci in human saliva. Growth curves of oral streptococci species **(A)** *S*. sp. A12, **(B)** *S. cristatus* ATCC 51100, **(C)** *S. gordonii* DL1, **(D)** *S. mitis* ATCC 49456, **(E)** *S. mutans* UA159, **(F)** *S. oralis* 34, **(G)** *S. sanguinis* SK36 and **(H)** *S. sobrinus* 6715 in either 100% TY (TY-; light grey circles), 50% TY / 50% H_2_O (TY-Water; blue squares), or 50% TY / 50% human saliva (TY-Saliva; red hexagons). Data points for optical density at 600 nm recorded every hour over a 24 h period are shown. Growth characteristic heat maps of the eight different oral streptococci with varying percentages (0-100%) of human saliva mixed with TY. Characteristics shown include **(I)** doubling times (minutes), **(J)** final yield (OD_600 nm_) and **(K)** time to OD_600 nm_ ≥ 0.1 (hours). Dark blue denotes lower values with green-yellow denoting the highest values within each heat map. White boxes crossed out without values indicate no growth in that condition.

We next wanted to determine the optimal amount of human saliva to mix with TY-for the remainer of our experiments by recording doubling times, final yields and lag times **(Figure 1I-K)**. Each strain saw improvements in at least one of these categories with 25-75% saliva mixed in TY-. However, growth was impaired for some strains starting at 75% saliva and all strains with 90% saliva. Therefore, it was determined that a 50% TY-, 50% saliva mix would be used for all experiments going forward.

To ensure these measured growth enhancements with human saliva were not just TY-specific, we performed a similar set of growth assays in the chemically defined medium CDM (24–26) **(Figure S3)**. Here, all eight streptococci species displayed growth improvements over CDM alone with both *S. mitis* and *S. oralis* only growing in the CDM-Saliva condition. These data demonstrate that the growth of several oral streptococci species are improved in mediums containing human saliva.

### Biofilm architecture is modified with human saliva

Previous studies have shown that salivary mucins can reduce oral streptococci biofilm formation (27, 28). To determine if similar reductions occur with human saliva, we first compared the accumulated biomass after a 24 h period using a crystal violet biofilm assay **(Figure 2AB)**. All seven species, except for *S. sobrinus*, had similar or lower quantifiable biomass with the addition of saliva. Interestingly, *S. sobrinus* nearly doubled its accumulated biomass in TY-containing saliva.

**Figure 2.**
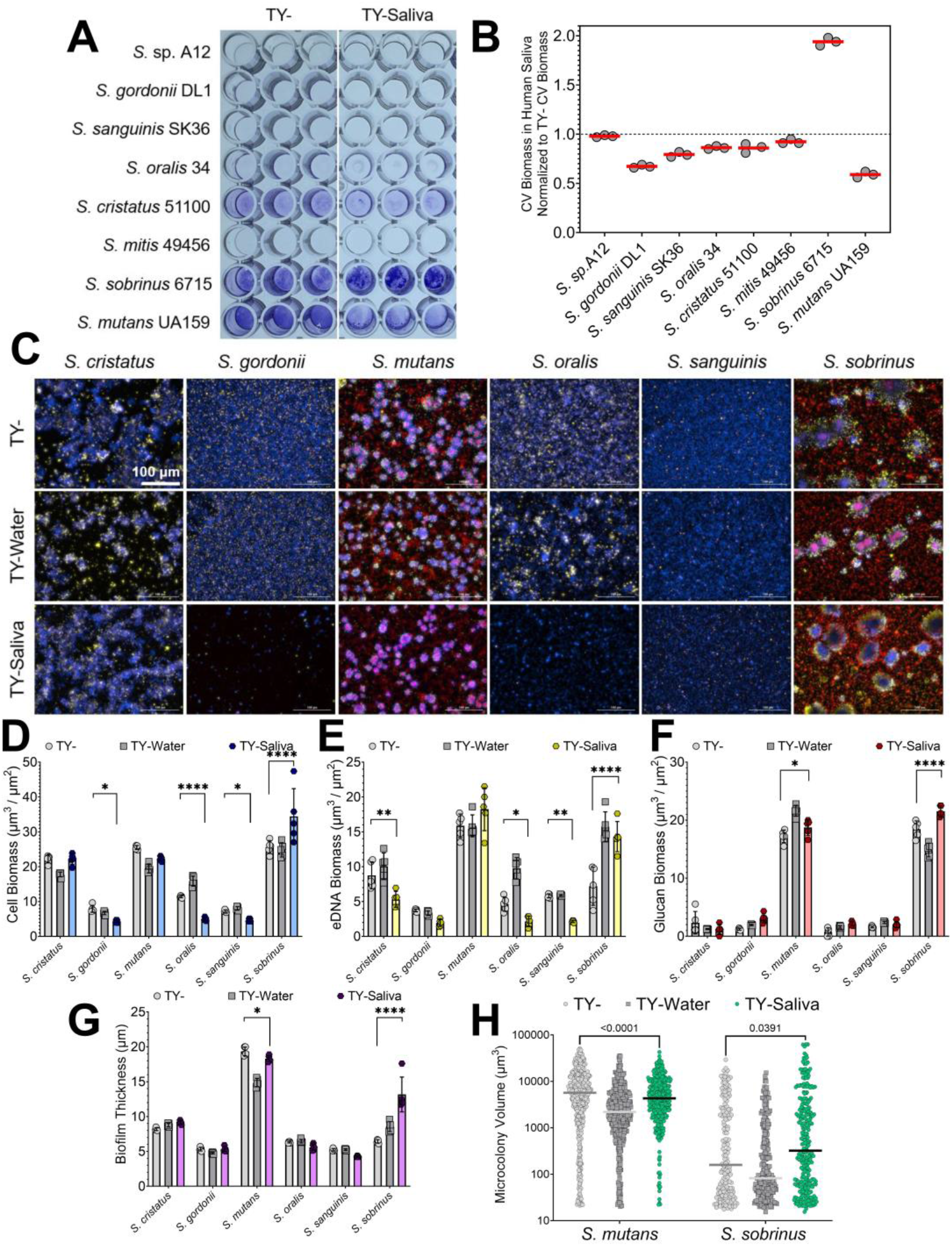
Biofilm formation of oral streptococci in human saliva. **(A)** Representative image of a crystal violet (CV) biofilm biomass assay for eight different oral streptococci species (listed on y-axis) in TY- or TY-Saliva. **(B)** CV biomass in TY-Saliva normalized to values obtained from TY-only. Data was quantified from the image shown on the left by extracting the crystal violet with 30% acetic acid and measuring the absorbance at 575 nm. The red bar denotes the mean of three biological replicates (grey circles). **(C)** Merged representative maximum intensity 20x Z-projections of 24 h oral streptococci biofilms grown in either TY-(top), TY-Water (middle), or TY-Saliva (bottom). Bacterial cells were stained with Hoechst 33342 (blue), eDNA probed with labeled antibodies (yellow), and glucans visualized with labeled dextran (red). Scale bar (100 µm) is shown in top left image. **(D)** Quantified cell, **(E)** eDNA, and **(F)** glucan biomass (µm^3^/µm^2^) along with **(G)** biofilm thickness (µm) and **(H)** individual microcolony volume from the image set shown in panel A between TY- (light grey circles), TY-Water (dark grey squares) or TY-Saliva (colored hexagons). Bars represent the mean of biomass or thickness from five independent images acquired (n = 5) with standard deviation. *S*. sp. A12 and *S. mitis* were not included within this panel, as they do not form adherent biofilms in monoculture. Quantification was completed using Gen5 Image+ software. Data graphing and two-way analysis of variance (ANOVA) with multiple comparisons was completed in GraphPad Prism software. * *p* < 0.05; ** *p* < 0.01; *** *p* < 0.001; **** *p* < 0.0001.

To better understand biofilm compositional and structural changes that may be occurring, we imaged and quantified single species biofilms in our three different growth mediums (TY-, TY-Water and TY-Saliva) **(Figure 2C-H)**. We observed a significant reduction in *S. gordonii, S. oralis* and *S. sanguinis* cell biomass along with a decline in *S. cristatus, S. oralis* and *S. sanguinis* eDNA biomass in TY-Saliva compared to TY-. However for *S. sobrinus*, a significant increase in cell, eDNA and glucan biomass was recorded. This additionally led to an overall increase in biofilm thickness and individual microcolony volumes. We also observed a change in biofilm matrix localization patterns within *S. sobrinus* microcolonies – eDNA appeared much more abundantly around the periphery of the microcolony. This was not seen with fellow mutans group species *S. mutans* that showed a reduction in individual microcolony volume in saliva. Overall, human saliva alters biofilm accumulation and architecture of oral streptococci species while enhancing biofilm formation of an odontopathogen (*S. sobrinus*).

### Similar changes in growth and biofilm formation with saliva-derived bacterial isolates

To verify that documented changes in oral streptococci growth and biofilm formation patterns with saliva were not limited to lab-adapted strains, we first isolated 29 different bacterial colonies that grew on BHI agar plates from our commercially-sourced human saliva. Viable colonies were streaked for isolation of a single colony morphology and then species identified by 16S sequencing. Identities returned included a range of different *Streptococcus, Actinomyces, Rothia* and *Granulicatella* species. We then monitored the growth of each isolate, given a SOSUI identifier number, in TY-, TY-Water and TY-Saliva. Doubling times for 26 out of 29 isolates were quickest in the TY-Saliva condition, and 18 out of 29 recorded their highest final yields in this condition as well **(Figure 3AB)**. Of note, two *A. oris* isolates (SOSUI_020 and SOSUI_026) and a *Streptococcus mitis* isolate (SOUSI_012) struggled to grow in either of the TY- and TY-Water conditions, but grew well only in mediums that contained saliva. To measure potential changes in biofilm formation, we first screened 17 of the isolates (SOSUI_001 – SOSUI_018, SOSUI_003 failed to grow after isolation and was excluded from all experiments) for measurable biomass accumulation after 24 h using the crystal violet method **(Figure S4)**. Four isolates, SOSUI_001, SOSUI_011, SOSUI_016 and SOSUI_018, were selected to carry forward for further analysis (each isolate also represented a different genus). Three of the four isolates saw significant reductions in biomass accumulation in TY-Saliva compared to TY-while the fourth, SOSUI_16, saw a non-significant 75±1% reduction **(Figure 3C)**. Similar patterns of reduction in both cell biomass as well as biofilm matrix components could be visualized by microscopy **(Figure 3DE)**. These data suggest that observations made with lab-adapted oral streptococci species are similar to phenotypes displayed by saliva-derived isolates including non-*Streptococcus* species.

**Figure 3.**
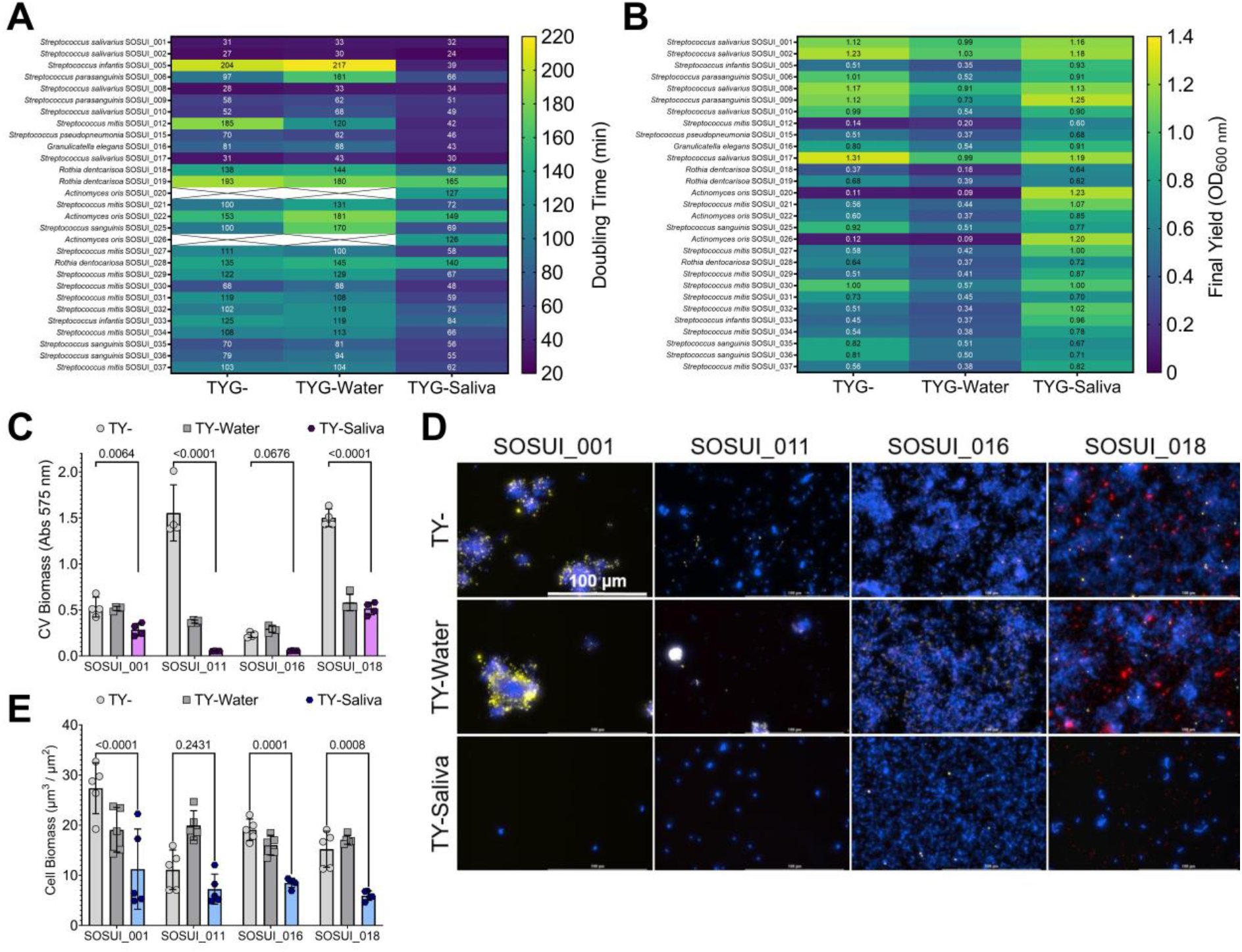
Growth and biofilm formation of human saliva-derived bacterial isolates. Growth characteristic heat maps of **(A)** doubling times (minutes) and **(B)** final yield (OD_600 nm_) shown for 29 different human saliva-derived bacterial isolates. Their SOSUI identifier #, along with species identity, is shown on the left y-axis. Dark blue denotes lower values with green-yellow denoting the highest values within each heat map. White boxes crossed out without values indicate no growth in that condition. **(C)** CV biomass (absorbance at 575 nm) of SOSUI_001, _011, _016, and _018 in TY- (light grey circles), TY-Water (dark grey squares) or TY-Saliva (purple hexagons). n = 4. **(D)** Merged representative maximum intensity 40x Z-projections of 24 h saliva isolate biofilms grown in either TY- (top), TY-Water (middle), or TY-Saliva (bottom). Bacterial cells were stained with Hoechst 33342 (blue), eDNA probed with labeled antibodies (yellow), and glucans visualized with labeled dextran (red). Scale bar (100 µm) is shown in top left image. **(E)** Quantified cell biomass (µm^3^/µm^2^) from the image set shown in panel D between TY- (light grey circles), TY-Water (dark grey squares) or TY-Saliva (colored hexagons). n = 5. Quantification was completed using Gen5 Image+ software. Data graphing and two-way analysis of variance (ANOVA) with multiple comparisons was completed in GraphPad Prism software with resulting *p*-values displayed.

### Saliva enhances the competitive behaviors of S. mutans

Previous studies reported that salivary mucins can alter the competitive behaviors between *Streptococcus mutans* and commensal streptococci (29). To determine if similar changes occurred with human saliva, we performed a fluorescent-intensity based coculture competition assay between a chromosomally integrated, constitutively-expressing *S. mutans gfp* strain (30) and unmarked competitor species **(Figure 4)**. Specific growth of *S. mutans* could be tracked via fluorescent intensity measurements of green fluorescent protein (GFP) every half hour over a 24 h period, while optical density of the entire coculture was also recorded. An area under the curve (AUC) was calculated using the intensity measurements and was compared statistically between the three different growth mediums. A significant increase in *S. mutans* intensity was recorded for five out of the seven cocultures within the TY-Saliva condition. *S. mutans* additionally displayed increased intensity within cocultures using CDM-, ensuring this change in competitive behavior was not specific for TY-**(Figure S4)**.

**Figure 4.**
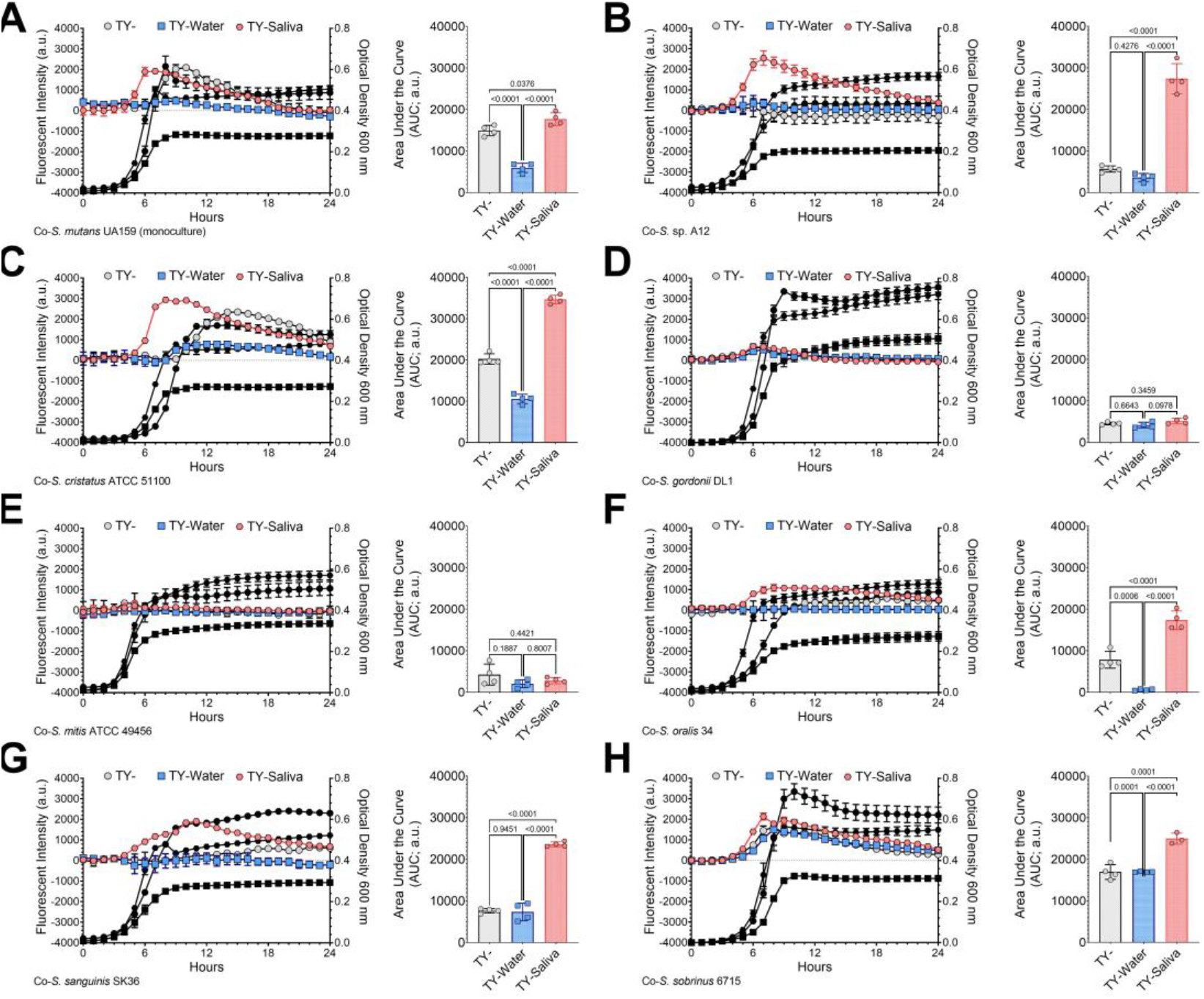
GFP-based competition assays between S. mutans and commensal streptococci competitors. Fluorescence-based (intensity, arbitrary units, a.u., left y-axis) growth profile of *S. mutans* in coculture competition against **(A)** *S. mutans* UA159 (monoculture growth control), **(B)** *S*. sp. A12, **(C)** *S. cristatus* ATCC 51100, **(D)** *S. gordonii* DL1, **(E)** *S. mitis* ATCC 49456, **(F)** *S. oralis* 34, **(G)** *S. sanguinis* SK36 and **(H)** *S. sobrinus* 6715 in either TY- (light grey circles), TY-Water (blue squares), or TY-Saliva (red hexagons). Growth of the entire coculture, measured by optical density at 600 nm, is shown in black using the same symbol (right y-axis). The area under the curve (AUC) of the fluorescent intensity of each condition was quantified and is shown in the graft on the right (n = 4). Data graphing and one-way analysis of variance (ANOVA) with multiple comparisons was completed in GraphPad Prism software with resulting *p*-values displayed.

To verify that the fluorescent-intensity based coculture competition led to quantifiable changes in cell number, we set up a competitive index assay between *S. mutans* and *S. oralis* that enumerated colony forming units (CFUs) during inoculation (t_i_ = 0 h) and at the time of cell harvest (t_f_ = 24 h) **(Figure 5)**. We recorded a negative competitive index for *S. mutans* in TY- (i.e., favoring *S. oralis*), but the competitive index flipped positive in TY-Saliva (i.e., favoring *S. mutans*). Using the constitutively-expressing *gfp* strain, we also found significantly higher quantified *S. mutans* biomass within cocultured biofilms with *S. oralis* and *S. sanguinis* that increased overall biofilm thickness, and for *S. oralis*-inoculated biofilms, also led to more eDNA and glucans within the biofilm matrix. Interestingly, lower *S. mutans* biomass was observed in coinoculated biofilms with *S. cristatus* and *S. sobrinus*. Finally, *S. mutans* cocultures with five out of the seven competitors had increased biomass via crystal violet with increasing amounts of saliva mixed in TY-**(Figure S5)**. Together, these data indicate that human saliva can enhance the competitive behaviors of *S. mutans* during growth with some commensal species.

**Figure 5.**
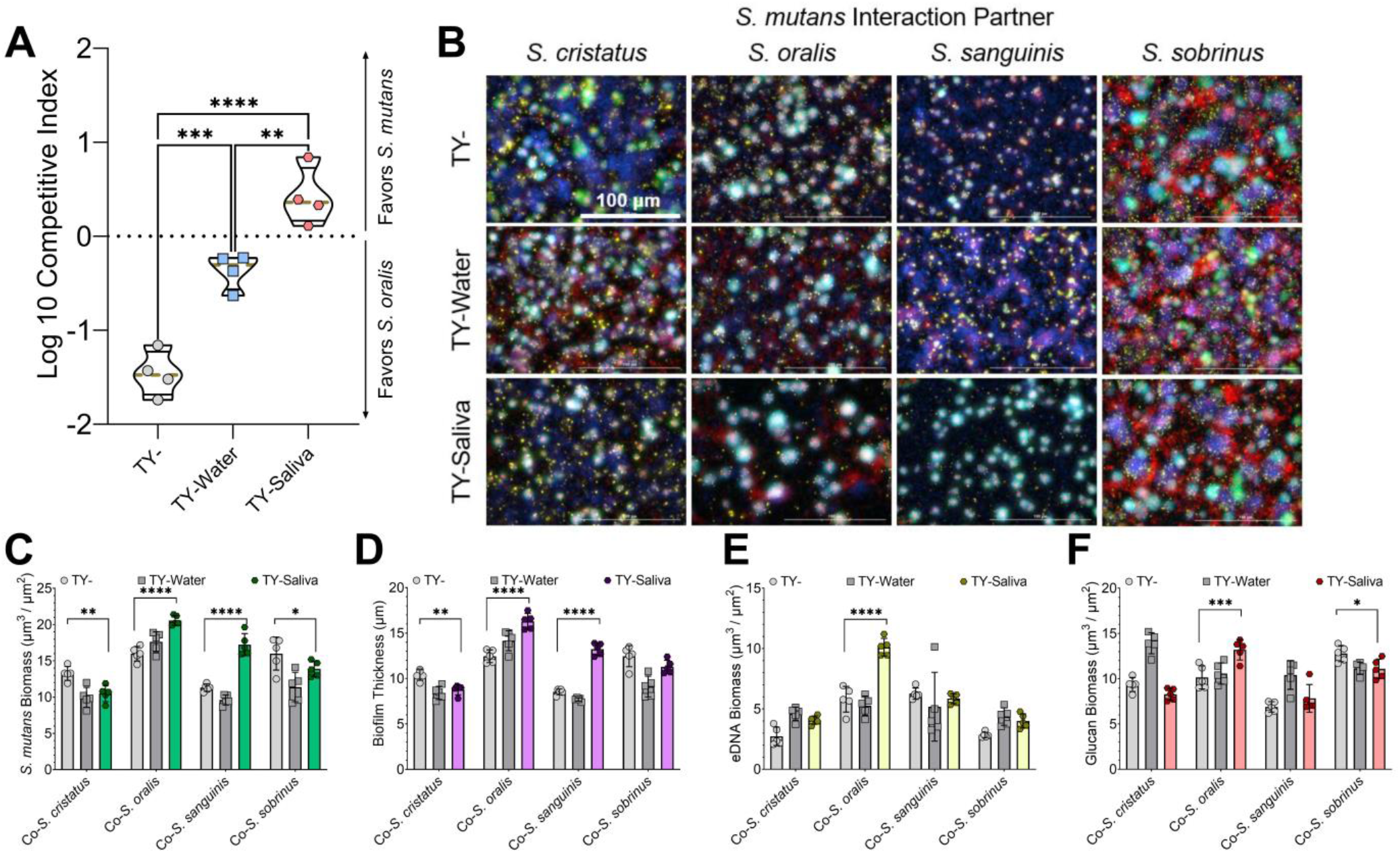
Increased competitiveness of S. mutans within human saliva. **(A)** Violin plot of Log10 competitive index of *S. mutans* in coculture competition against *S. oralis* in either TY- (light grey circles), TY-Water (blue squares), or TY-Saliva (red hexagons). Positive values represent a competitive advantage for *S. mutans* while negative values represent a competitive advantage for *S. oralis*. Black solid lines represent the quartiles while a gold dashed line represents the median. n = 4. **(B)** Merged representative maximum intensity 40x Z-projections of 24 h cocultured biofilms between a *S. mutans* GFP+ strain (green) and oral commensal competitors (stained with Hoechst 33342, blue). Biofilm matrix components eDNA was probed with labeled antibodies (yellow) and glucans visualized with labeled dextran (red). **(C)** Quantified *S. mutans* biomass (µm^3^/µm^2^), **(D)** biofilm thickness (µm) **(E)** eDNA and **(F)** glucan biomass from the image set shown in panel B between TY- (light grey circles), TY-Water (dark grey squares), or TY-Saliva (colored hexagons). Bars represent the mean of biomass or thickness from five independent images acquired (n = 5) with standard deviation. Quantification was completed using Gen5 Image+ software. Data graphing and one-way or two-way analysis of variance (ANOVA) with multiple comparisons was completed in GraphPad Prism software. * *p* < 0.05; ** *p* < 0.01; *** *p* < 0.001; **** *p* < 0.0001.

### Human saliva reprograms the transcriptome of oral streptococci

To understand how saliva affects changes in growth, biofilm formation and competition between the oral streptococci, we performed RNA-Seq on monocultures of *S. mutans* and *S. oralis* in TY-Saliva and compared to TY- **(Figure 6)**. We found 133 differentially expressed genes (DEGs) in *S. mutans* grown in saliva **(Table S1)**, with the most upregulated genes (59 total DEGs) belonging to phosphotransferase systems (PTS) specific for fructose, mannose and/or N-acetyl glucosamine (GlcNAc) (*levDEFGX*, SMU.1956c – SMU.1961c) (31–33), cellobiose (SMU.1596 – SMU.1600) (34), and α-1,3-linked carbohydrates (SMU.100 – SMU.105) (35). Other upregulated genes of note included the glucosyltransferases *gtfBC* (SMU.1004, SMU.1005), the bacitracin resistance ABC transporters *mbrAB* (SMU.1006-SMU.1007) (36), and the TnSmu2 gene cluster which contains genes for mutanobactin synthesis (37). In contrast, many of the downregulated genes (74 DEGs) belonged to the integrative and conjugative element TnSmu1 (38, 39), the genetic competence regulon (40), a CRISPR2-Cas system (41, 42) and predicted amino acid ABC transporters (SMU.932-SMU.936). Interestingly, the most downregulated genes within the dataset belonged to the trehalose-specific PTS (43, 44). The commensal *S. oralis* had a higher number of DEGs overall (206 total, 11% of coding genome features) **(Table S2)**, consisting of upregulation (148 DEGs) in 18 different ABC transporters, 13 transcriptional regulators, 7 glycosyl hydrolases and genes involved in *de novo* purine and tryptophan biosynthesis. Downregulated genes (58 DEGs) included queuosine biosynthesis (45), transporters for iron (46), vitamin B12 (47) and glutamine (48), and transcriptional regulators *ciaR* (49) and *mntR* (50). These described DEGs highlight the alternations in metabolic networks for oral streptococci during growth in saliva.

**Figure 6.**
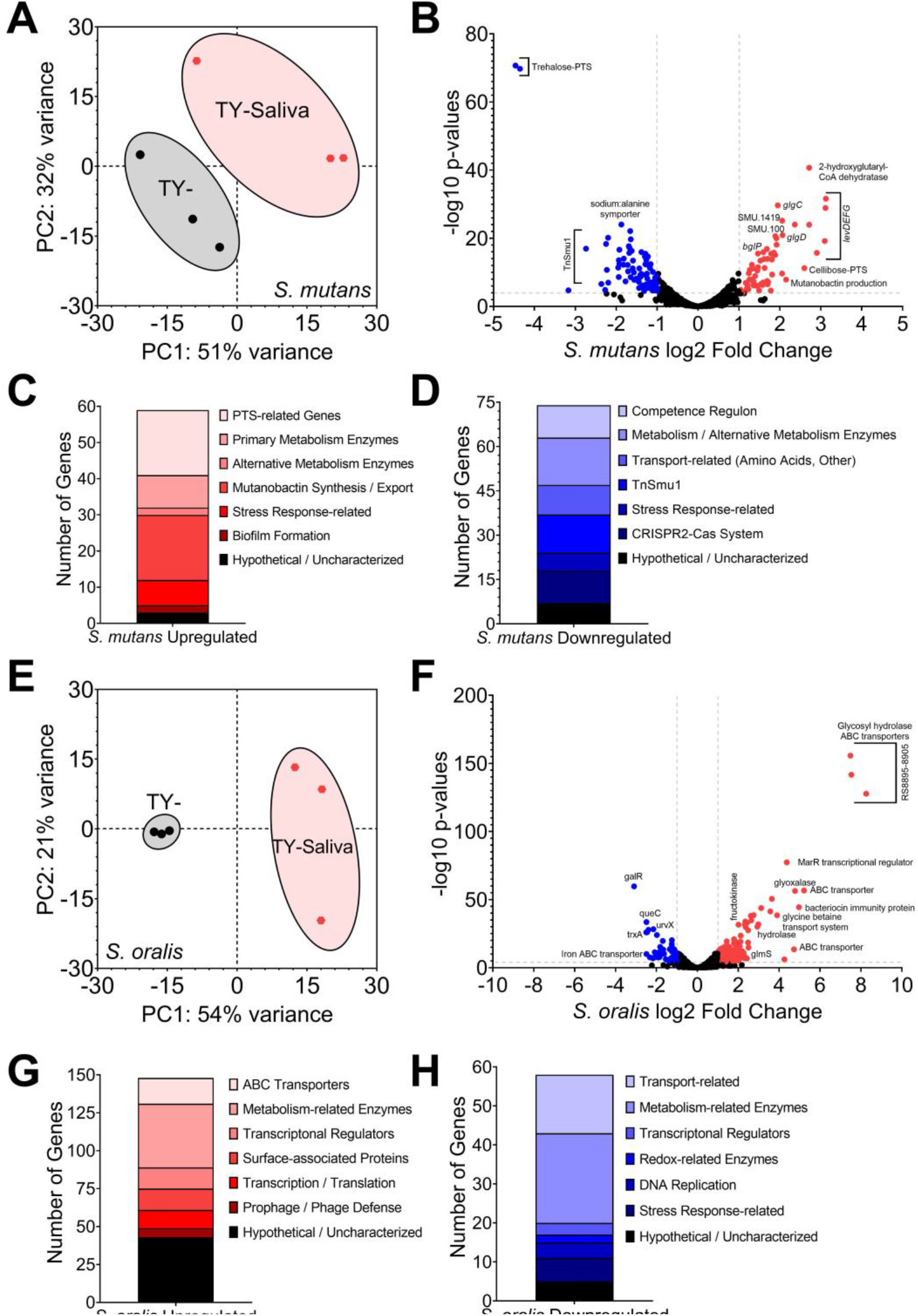
Transcriptome profiling of S. mutans and S. oralis monocultures in human saliva. **(A)** Principal component analysis (PCA) from RNA-Seq expression data (n = 3) of *S. mutans* monocultures grown in TY- (black circles) or TY-Saliva (red hexagons). The proportion of variance for either PC1 (x-axis) or PC2 (y-axis) are listed. **(B)** Volcano plot of changes within individual *S. mutans* genes (circles) between TY- and TY-Saliva. Differentially expressed genes (DEGs; = genes with ≥4 Log10 *p* value and Log2 fold change ≥ (-)1) are shown in either red (upregulated, right) or blue (downregulated, left). Individual gene identifier, name and/or characterizable function are displayed, if able. **(C)** Stacked bar chart of upregulated *S. mutans* DEGs grouped by pathway/operon/function. **(D)** Stacked bar chart of downregulated *S. mutans* DEGs. **(E)** PCA from RNA-Seq expression data of *S. oralis* monocultures (n = 3). **(F)** Volcano plot of changes within individual *S. oralis* genes. **(G)** Stacked bar chart of upregulated *S. oralis* DEGs. **(H)** Stacked bar chart of downregulated *S. oralis* DEGs. DEGs were determined from Degust using edgeR analysis. Data graphing and PCA calculations were completed in GraphPad Prism software.

Recently our group highlighted a repetitive DEG pattern for *S. mutans* in coculture with commensal streptococci that included *S. oralis* (14). We wanted to determine how coculture growth in medium containing saliva modified this pattern and the overall interaction between the species. To accomplish this, we first compared DEGs of both *S. mutans* and *S. oralis* grown in coculture TY-Saliva to coculture TY-**(Figure 7)**. By already characterizing monoculture responses to saliva, we were able to categorize genome features that overlapped with species-specific growth in saliva as well as isolate new features that were specific to the coculture interaction. For *S. mutans*, this included upregulation of transporters related to manganese (*slo* operon and *mntH*) (51), maltose (*malXFGK*) (52), glutamine, glutathione and L-cysteine uptake (SMU.1939c-SMU.1942c) (53), the rhamnose-glucose cell wall polysaccharide (*rgp*) operon (54, 55) and genes within the citrate pathway (SMU.670-SMU.672) for glutamate biosynthesis (56) **(Table S3)**. Additionally the trehalose-specific PTS, the most downregulated genes in monoculture, were now upregulated two-fold in coculture. The lactose utilization operon (*lacABCDEFGX*) (57, 58) was downregulated in coculture saliva growth, but not in monoculture. We additionally compared the DEGs during transition from monoculture to coculture growth with *S. oralis* in either TY- and TYG-Saliva (i.e., DEGs from coculture in TY-vs DEGs from coculture in TY-Saliva). Of the 219 total *S. mutans* DEGs captured during coculture growth with *S. oralis* in both medium conditions, 78 were specific to TY-, 108 specific to TY-Saliva and 33 were present under both conditions **(Table S4)**. These 33 genome features consisted of the CRISPR2-Cas system, *levDEFGX*, the transglycosylase SMU.2146c, the LexA-like transcriptional regulator *hdiR* (SMU.2027), and amino acid ABC transporters SMU.932-SMU.936.

**Figure 7.**
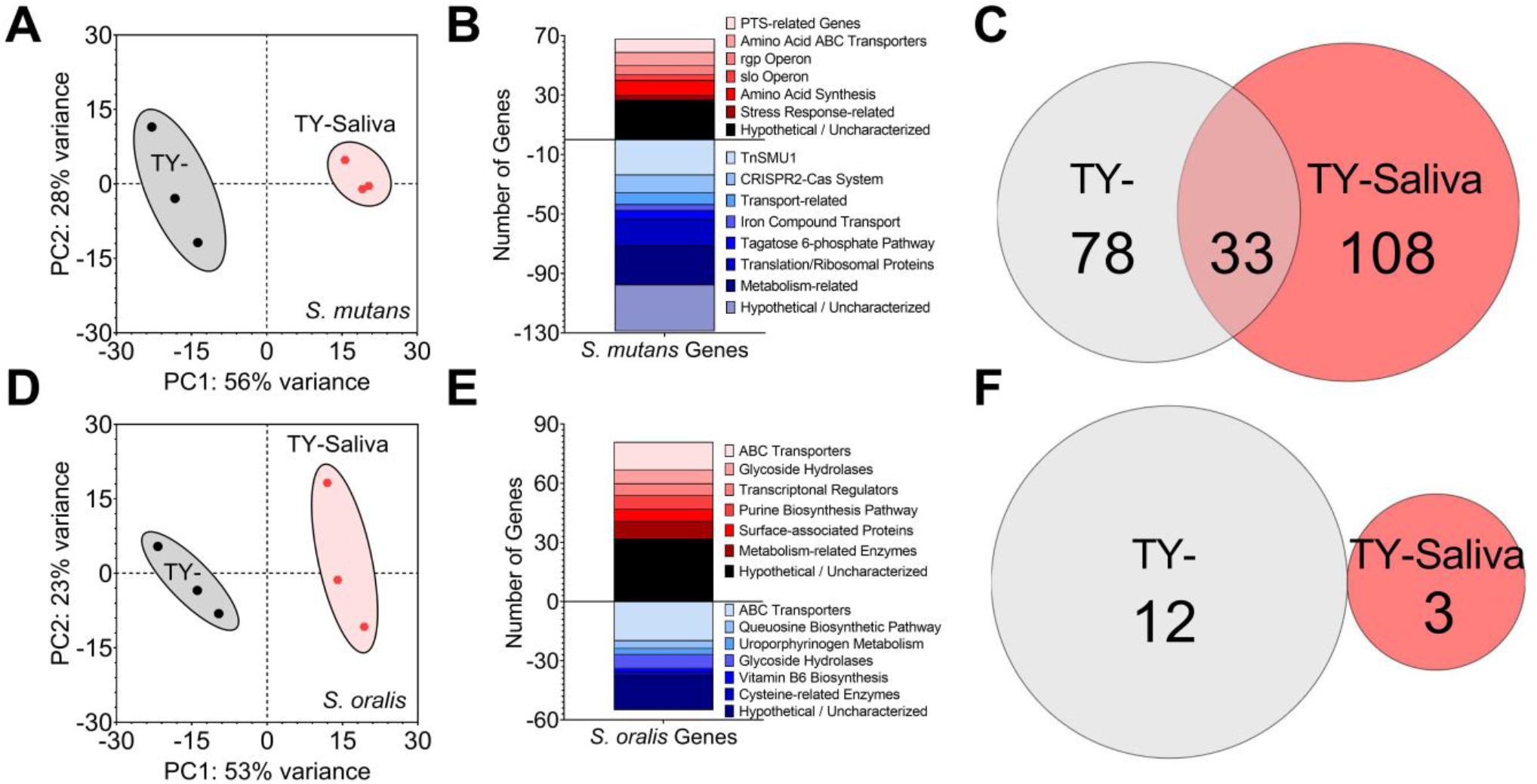
Transcriptome profiling of S. mutans and S. oralis cocultures in human saliva. **(A)** Principal component analysis (PCA) from RNA-Seq expression data (n = 3) of *S. mutans* cocultures with *S. oralis* grown in TY- (black circles) or TY-Saliva (red hexagons). The proportion of variance for either PC1 (x-axis) or PC2 (y-axis) are listed. **(B)** Stacked bar chart of *S. mutans* upregulated (red, positive) and downregulated (blue, negative) differentially expressed genes (DEGs) grouped by pathway/operon/function. **(C)** Venn diagram of the number of *S. mutans* DEGs in coculture with *S. oralis* specific to TY- (light grey), TY-Saliva (red) or common to both conditions (overlap). **(D)** PCA from RNA-Seq expression data of *S. oralis* cocultures with *S. mutans* (n = 3). **(E)** Stacked bar chart of *S. oralis* upregulated and downregulated DEGs. **(F)** Venn diagram of the number of *S. oralis* DEGs in coculture with *S. mutans* specific to each condition.

For *S. oralis*, only 24 DEGs were isolated genome features specific to the coculture interaction **(Table S5)**. Eleven were upregulated and included genes within the tagatose 6-phosphate pathway, an endo-α-*N*-acetylgalactosaminidase (59), an *N*-acetylmannosamine kinase (60) and α-mannosidase, and a thiamin transporter while downregulated genes included *oppA*, other amino acid transporters and the uroporphyrinogen metabolism pathway. For comparison of DEGs during transition from monoculture to coculture, we only found 15 total DEGs with no overlap between conditions **(Table S6)**. In summary, the intermicrobial interactions between *S. mutans* and *S. oralis* are amended in the presence of human saliva and suggest changes in preferred nutritional niches for each species.

### S. mutans growth enhanced with manganese but not alternative carbohydrates

To determine if some of the results from our transcriptomics analysis can be utilized in better understanding the interaction between *S. mutans* and *S. oralis*, we modified the culturing condition by supplementing media with either metals or alternative carbohydrates. Addition of 1 µM iron (Fe), manganese (Mn) or zinc (Zn) to monocultures of *S. mutans* within our fluorescent-intensity based coculture competition assay showed significant inhibitory growth defects only with Mn **(Figure 8A)**. However, in coculture competition with *S. oralis*, addition of Mn significantly benefited *S. mutans* while Zn was now inhibitory **(Figure 8B)**. This result validates some of the coculture characterization via RNA-Seq where manganese transporters were upregulated, but also reveals an unexpected phenotype with zinc. In a similar way, we wanted to determine if documented changes in carbohydrate uptake and utilization pathways could shift competitive behaviors. We first compared fructose, galactose and GlcNAc as sole available carbohydrate sources in comparison to glucose in TY- and TY-Saliva. While *S. mutans* growth in all TY-Saliva cultures significantly improved over TY-, there were no significant differences between mediums that contained any of the other carbohydrates and glucose **(Figure 8C)**. Finally, we tested whether having an additional, alternative carbohydrate source (5 mM or 2.5 mM each) available alongside glucose (15 mM) affected the competitiveness of *S. mutans*. Again, while all TY-Saliva cultures supported better overall *S. mutans* growth compared to TY-, there were no significant changes compared to glucose alone **(Figure 8D)**. Together these data indicate that supplementation of metals has potential to alter the competition between *S. mutans* and commensals, while a strategy centered on alternative carbohydrates was not apparent within these sets of assays.

**Figure 8.**
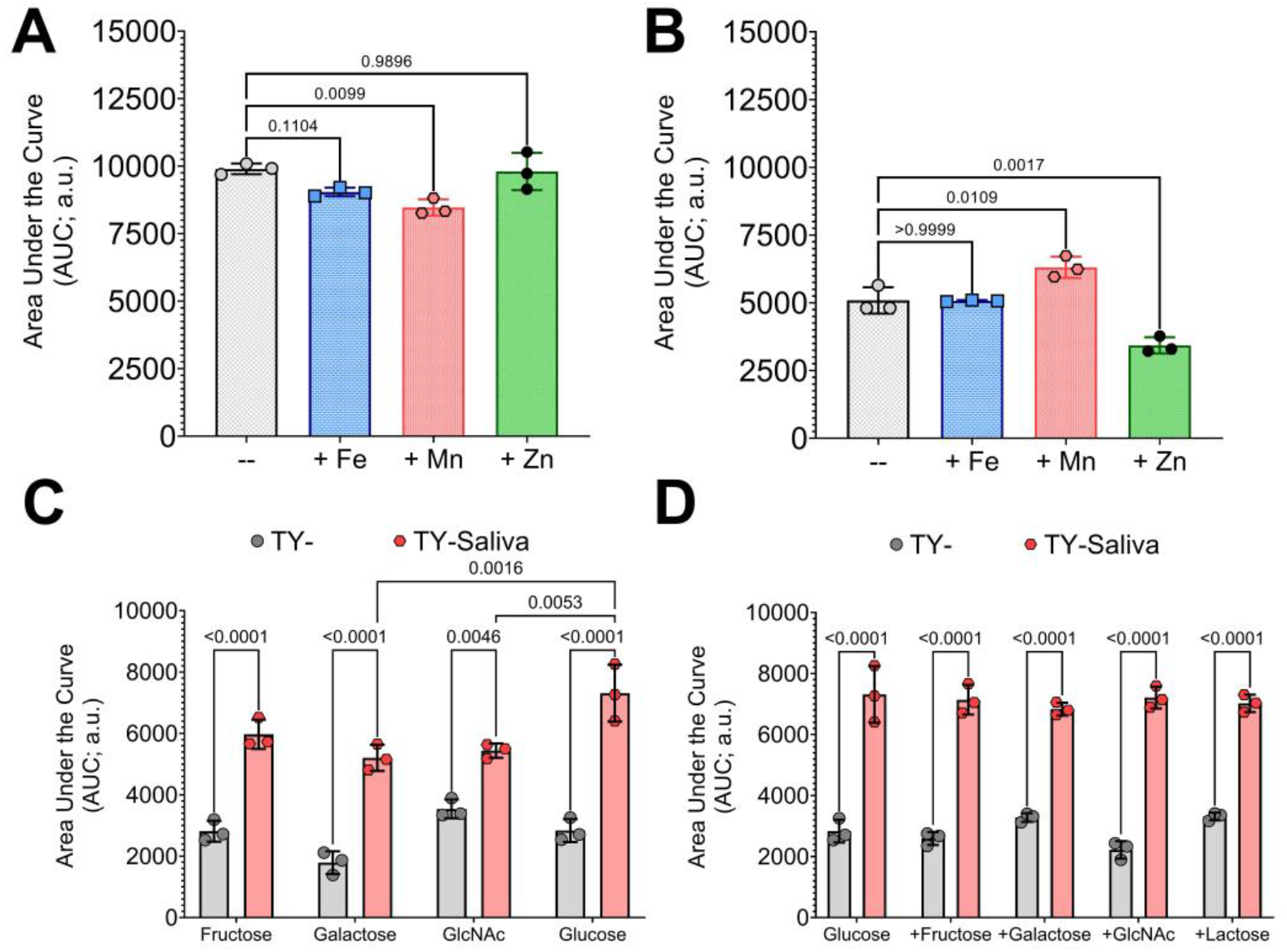
Altering the competitive behaviors of S. mutans with metals or alternative carbohydrate sources. **(A)** Area under the curve (AUC) of the fluorescent intensity (arbitrary units, a.u.) from GFP-based competition assays between *S. mutans* and *S. oralis*. **(A)** *S. mutans* monoculture growth in either TY- (grey circles-bar) or TY-supplemented with 1 µM FeSO_4_ (blue squares-bar), MnSO_4_ (red hexagons-bar) or ZnSO_4_ (green diamonds-bar). **(B)** *S. mutans* coculture growth with *S. oralis* with TY- or 1 µM FeSO_4_, MnSO_4_ or ZnSO_4_. **(C)** *S. mutans* coculture growth in either TY- (grey circles-bar) or TY-Saliva (red hexagons-bar) with 20 mM fructose, galactose, N-acetyl glucosamine (GlcNAc) or glucose. **(D)** *S. mutans* coculture growth in 15 mM glucose or 15 mM glucose with addition of 5 mM fructose, galactose, GlcNAc or 2.5 mM lactose. n = 3. Data graphing and one-way or two-way analysis of variance (ANOVA) with multiple comparisons was completed in GraphPad Prism software with resulting *p*-values displayed.

## DISCUSSION

Our foray into examining the growth and competition of oral streptococci in human saliva began with the goal of documenting how the interaction(s) between an odontopathogen (*S. mutans*) and commensal streptococci are modified when cultured in conditions that include the presence of saliva. To accomplish this, we first needed to determine how saliva altered the growth patterns and biofilm formation of these species, as well as the optimal growth condition to conduct the observations. We were able to verify prior reports that most streptococci strains are not able to grow in 100% saliva unless glucose is added as a carbon source (61). Even then, *S. gordonii* was the only species that achieved growth levels that would have been suitable for competition assays and harvesting of RNA, which also follows previous observations (62). Therefore, we used the strategy of mixing our lab-based experimental medium, TY-, with different proportions of saliva. Conveniently, mixing an equal volume of TY with saliva led to growth benefits for almost all eight species of streptococci tested, that included faster doubling times and higher final yields. Including even just small amounts of saliva led to growth benefits, with *S. oralis* recording faster doubling times in TY-with as little as 1% saliva added **(Figure S7)**. Mixing amounts higher than 50% began a reversal in doubling times and yields for some species, with detrimental effects seen for all species at a 90% saliva mix. While the mixing of saliva and lab-based mediums does not entirely replicate an *in vivo* condition, it does allow for similar testing and sampling procedures as lab-based mediums alone, which encompasses both user handling convenience as well as inclusion of host-derived factors into *in vitro* experiments. This offers an alternative to several artificial saliva mediums – inclusion of mucin, heme and sheep’s blood within these formulations often make optical density measurements and fluorescent intensity-based assays difficult if not impossible which are fundamental assays within this report (63, 64). However, the tradeoff of our approach is a large volume requirement of human saliva that may be hard to obtain from volunteer donors and/or a higher cost from commercial sources, which can be a prohibitive barrier for researchers. One surprise to us was finding several species that clearly grew better with the addition of saliva such as *S. cristatus* and *S. oralis*, with saliva even allowing for growth of *S. oralis* and *S. mitis* in CDM. While we have not yet determined the factor(s) present in saliva that are promoting growth under these conditions, the displayed phenotypes should provide a clear test to determine their identity. These discoveries could also be translatable to other species, as similar growth-promoting phenotypes were recorded with saliva-derived isolates of *Actinomyces, Rothia* and *Granulicatella* species.

We also found saliva to limit the amount of biofilm biomass accumulation for most species, aside from *S. sobrinus*. The amount of *S. sobrinus* accumulated biomass, measured by crystal violet staining, doubled in medium containing saliva that could be attributed to increases in both cell and biofilm matrix (eDNA and glucans) biomass upon microscopy visualization and quantification. This ultimately led to larger volumes within *S. sobrinus* microcolonies, which may be a virulence determinant for the initiation of dental caries. Interestingly, the amount of eDNA also increased in the TY-Water condition. Higher amounts of eDNA in TY-Water could also be seen with *S. cristatus, S. oralis*, and most notably with saliva-derived isolates SOSUI_001 and SOSUI_011. This may represent increased cell lysis and release of DNA, and/or an undescribed lysis-independent mechanism with higher activity under this condition. Still, *S. sobrinus* was the only species to have higher amounts of eDNA in TY-Saliva compared to TY-. The role of this increased eDNA accumulation and whether this phenotype is in fact specific to *S. sobrinus* or found with other mutans group streptococci strains (was not observed in *S. mutans* UA159) remains to be explored.

Another surprise occurred during both GFP- and CFU-based coculture competitions between *S. mutans* and several different commensal species. The accumulation of our data suggests that growth in human saliva enhances the competitive behaviors of *S. mutans*, leading ultimately to higher recoverable cell numbers and accumulation of biomass within biofilms. This was true for four out of the six different commensal species we tested, as well as with *S. sobrinus* that is commonly found together with *S. mutans* in carious lesions (65–67). Our current working hypothesis is that saliva “jump starts” carbohydrate uptake and glycolytic pathways in *S. mutans* that are seemingly not as active in medium lacking saliva during growth in coculture against commensal streptococci **(Figure 9)**. This allows the commensals to effectively outcompete *S. mutans* in lab-based medium alone, but quickly lose this advantage in mediums that contain saliva, even when their own growth is improved as well. We were able to capture via transcriptional profiling upregulation of at least three *S. mutans* PTS operons in saliva, as well as the trehalose PTS and maltose ABC transporters during coculture with *S. oralis*. We also saw upregulation of *fruAB*, pyruvate formate-lyase (*pfl*) and the glycogen synthesis operon (*glg*), suggesting heterofermentative growth on non-glucose sugars with potential concurrent intracellular polysaccharide (IPS) accumulation. Addition of a variety of carbohydrates likely present in human saliva that are not found in the TY-only condition is plausibly responsible for these transcriptional changes, even if present in low µM amounts. While we did not specifically test for other carbohydrates, we did quantify less than 125 µM glucose present in our saliva preparations, similar to other previously reported values (15, 18). Additionally, we found upregulation of genes responsible for mutanobactin synthesis, a compound described to be important for commensal-produced hydrogen peroxide tolerance (37). Enhanced *S. mutans* competitiveness may also be due, in part, to increased oxidative stress tolerance supplied by this compound. Genes related to genetic competence and bacteriocin production were downregulated, a similar transcriptional profile for *S. mutans* cultured with mucin *O*-linked glycans (68).

**Figure 9.**
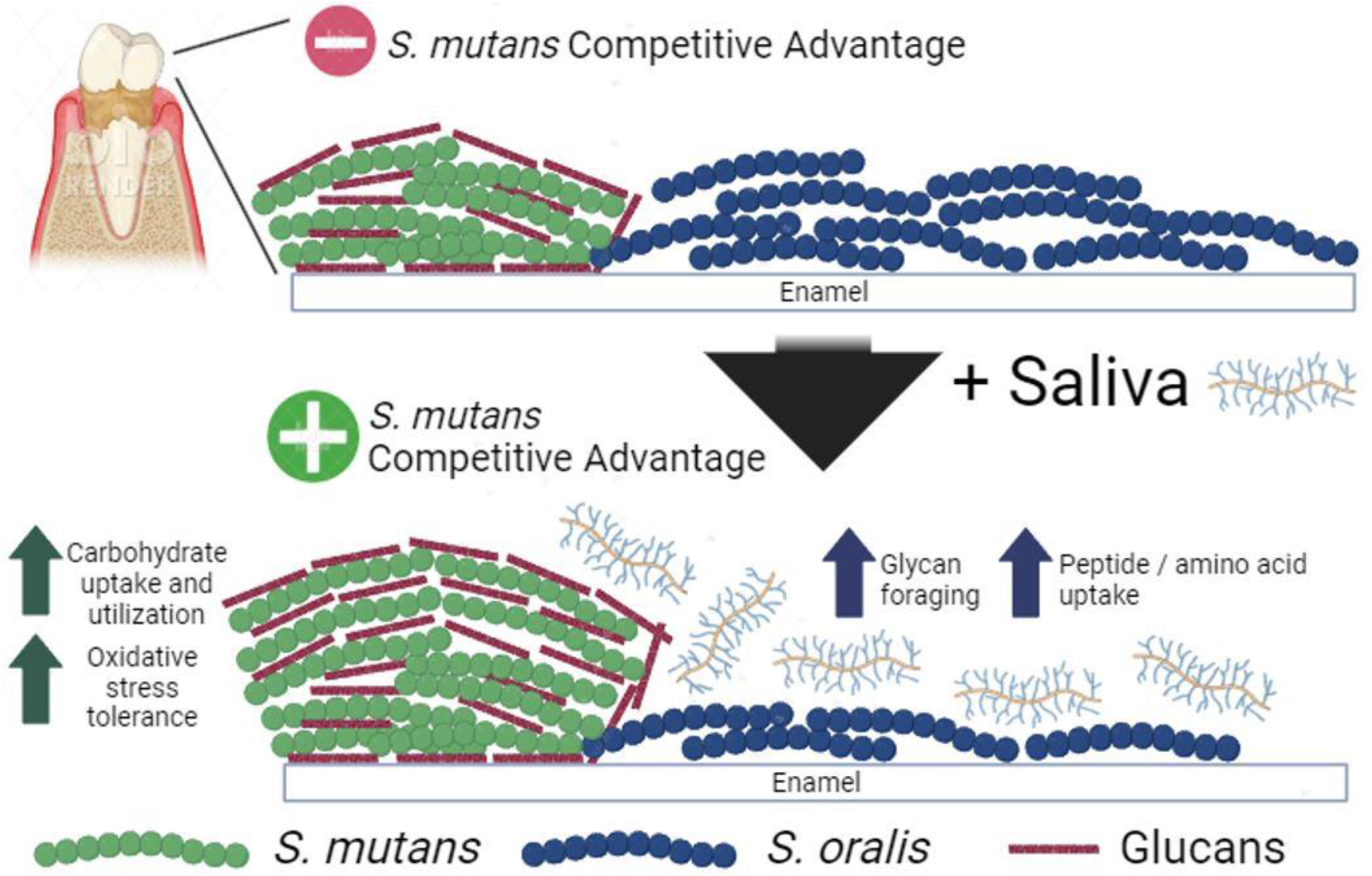
Saliva modifies the competitive behaviors of oral streptococci. Two oral streptococci, *S. mutans* (green) and *S. oralis* (blue), colonize supragingival biofilms attached to the tooth’s surface. In cocultures competed in lab-based medium, *S. oralis* maintains a competitive advantage over *S. mutans*. However when saliva is added to the medium, the competitive behaviors of *S. mutans* is enhanced through upregulation of operons related to carbohydrate uptake and utilization as well as pathways for oxidative stress tolerance. In *S. oralis*, the presence of saliva leads to enhanced expression of genes related to glycan foraging and amino acid transporters. Figure was created with BioRender.

Our transcriptional data of *S. mutans* and *S. oralis* in coculture also hints of each species developing nutritional niches under mixed-species growth. The best example of this involves the *lac* operon and tagatose 6-phosphate pathway for the metabolism of galactose. During coculture, *S. mutans* downregulates the genes within this pathway while they are concurrently upregulated in *S. oralis. S. mutans* also upregulates pathways for trehalose and maltose uptake, two carbohydrates that *S. oralis* 34 does not breach a final yield greater than 0.15 with after a 24 h period **(Figure S8)**. In turn, we see upregulation of genes in *S. oralis* related to glycan foraging such as *N*-acetylmannosamine kinase, the second enzyme within the sialic acid catabolic pathway, and endo-α-*N*-acetylgalactosaminidase, which catalyzes the hydrolysis of an *O*-glycosidic α-linkage between galactosyl β1,3 *N*-acetyl-D-galactosamine (Galβ1,3GalNAc) and the serine or threonine residue in mucins and mucin-type glycoproteins. Rather than competing for the same non-glucose carbohydrates, it is tempting to speculate that these organisms may be utilizing carbon sources the other species does not. To begin to test this hypothesis, we completed several versions of our fluorescence-based competition assay, but found no change in *S. mutans* growth when other alternative carbohydrates were provided alone or in combination with glucose. Other experimental protocols will be required to test this hypothesis. Of note, *S. mutans* is capable of metabolizing a wider variety of carbohydrates than any other Gram-positive organism (69) – targeting of transport systems that do not overlap with those found in commensal species may be an attractive strategy towards limiting its emergence. Likewise, the taxonomic diversity and resistance to dysbiosis of saliva-derived oral communities have recently been shown to be increased with mucin glycans (28), highlighting that these described nutritional niche partitionings may be key in maintaining microbiome homeostasis by preventing outgrowth of pathogens.

A final note was the observation of multiple oxidative stress tolerance pathways upregulated in *S. mutans* with saliva, especially under the coculture condition. This included upregulation of manganese transporters *sloABC* and *mntH*, recently characterized to show cooperative activity in manganese-restricted conditions (51, 70), as well as concurrent downregulation in iron transporters. Manganese can assist in oxidative stress tolerance through acting as a cofactor of superoxide dismutase to neutralize hydroxyl radicals (71), while free iron can produce hydroxyl radicals via Fenton chemistry (72). Additionally, manganese may also stimulate carbohydrate metabolism (73, 74). We noted increased *S. mutans* growth in coculture with addition of 1 µM manganese, a physiologically-relevant concentration that is present in resting saliva (75). This result not only validates our approach of characterizing intermicrobial interactions leading to strategies of manipulation that can shift the balance between organisms, but also highlights the critical role of trace metals in *S. mutans* physiology. In addition, we also noted upregulation in glutathione import and/or synthesis, another protective mechanism against oxidative stress tolerance (53).

Our study into how human saliva impacts the growth, biofilm formation and competitive behaviors between commensal and mutans group streptococci has solidified the importance of studying mixed-species interactions in conditions that more closely resemble their *in vivo* condition. While many lingering questions remain, including identification of specific streptococcal pathways that increase fitness during growth in saliva and whether niche partitioning plays a role in competition between species, our results provide new insights and findings that offer promising directions for the further evaluation of supporting the persistence of health-associated microbes while developing interventions to combat oral pathogens.

## MATERIALS AND METHODS

### Human Saliva and Growth Mediums

Commercially available pooled human saliva was purchased from Innovative Research (IRHUSL250ML). The saliva is collected fresh from two or more donors, pooled and stored frozen. Upon receipt, the saliva was thawed, centrifuged at 4500 RPM for 10 minutes to remove debris, and then passed through a Millipore Express PES Membrane 0.22 µm Filter Unit (SLGPR33RB) prior to local frozen storage as 10 mL aliquots. For experimental use, the saliva was thawed at room temperature and used same day. Bacterial strains used in this study **(Table S7)** were cultured in Bacto Brain Heart Infusion (BHI; 237500), Tryptone Yeast extract supplemented with Glucose (TYG, 20 mM glucose final concentration; 10 g tryptone [Fisher Bioreagents, BP1421], 5 g yeast extract [Fisher Bioreagents, BP1422], 3 g K_2_HPO_4_ [Sigma-Aldrich, P3786], and 3.6 g glucose per 1 L H_2_O [Sigma-Aldrich, G8270]) or the chemically-defined medium CDM (24–26). 1.7 g L^-1^ sucrose (Sigma-Aldrich, S7903) was added for all biofilm-related experiments (5 mM sucrose final concentration). To prepare mediums using –Water or –Saliva, a percentage (%) of the final volume of either liquid was first added to either TY or CDM that had been prepared without addition of carbohydrate. After mixing, carbohydrate(s) of choice were then added to all mediums for a consistent concentration within all groups being tested. Catalog numbers for addition carbohydrates and metals used within growth measurements and/or competitive assays are listed within **Table S8**.

### Overnight Cultures, Strain Inoculation and Growth Measurements

Overnight cultures of all strains were inoculated from single, isolated colonies on BHI agar plates into BHI broth and incubated at 37°C and 5% CO_2_ with the appropriate antibiotics. Antibiotics were added to overnight growth medium BHI at 1 mg/mL for both kanamycin and spectinomycin and 0.01 mg/mL for erythromycin. The next day, cultures were harvested by centrifugation, washed to remove all traces of overnight growth medium, and normalized to OD_600 nm_ = 0.1 with 1x phosphate-buffered saline (PBS) prior to back dilution (1:100) into experimental medium of choice. For coculture competitions with *S. mutans*, strains were inoculated according to **Table S9**. All strains were maintained for long-term storage at -80ºC in BHI containing 25% glycerol. Growth measurements were completed using a Bioscreen C MBR automated turbidometric analyzer (Growth Curves Ab Ltd., Helsinki, Finland) with the optical density at 600 nm (OD_600 nm_) recorded every 30 minutes for 24 h. Wells containing liquid culture medium were overlaid with 0.05 mL sterile mineral oil (Fisher Chemical, O121) to reduce the growth inhibitory effects of oxygen during the experiment. A media-only control (blank) was used to subtract background absorbance from all wells containing bacterial cells. Resulting data was used to calculate doubling time (D = [time in minutes; 120]/[n, number of generations]), final yield (average of OD_600 nm_ for last two time points recorded), and time to OD_600 nm_ = 0.1 (half hour interval when OD_600 nm_ was first > 0.1). All experiments were completed with at least three biological replicates measured in technical triplicates.

### Biofilm Microscopy

Bacterial strains were inoculated into medium that contained 1 µM Alexa Fluor 647-labeled dextran (10,000 molecular weight; anionic, fixable; Invitrogen, D22914), added to Cellvis 12 well, glass bottom, black plates (P12-1.5H-N) and incubated at 37°C and 5% CO_2_ for 24 h. Resulting biofilms were first washed with 1x PBS to remove loosely-bound cells and incubated with BSA blocking buffer at room temperature for 0.5 h (Thermo Scientific, 3% BSA is PBS; J61655.AK). Biofilms were then probed with two murine monoclonal antibodies against B-DNA (Anti-dsDNA, 3519 DNA, Abcam, ab27156) and Z-DNA (Absolute Antibodies Ab00783-23.0) [2 µg mL^-1^] within BSA blocking buffer for 1 h at room temperature. The biofilms were then washed once and incubated for 1 h at room temperature with an Alexa Fluor 594-labeled goat anti-mouse IgG highly cross-absorbed secondary antibody (Invitrogen, A32742)) [2 µg mL^-1^] within BSA blocking buffer. Finally, the biofilms were washed and stained with Hoechst 33342 solution (5 µM final concentration, Thermo Scientific, 62249) for 15 minutes. All biofilms were imaged within Invitrogen Live Cell Imaging Solution (1X, A14291DJ) using a 20x (plan fluorite, 6.7 mm working distance, 0.45 numerical aperture) or 40x (plan fluorite, 2.7 mm working distance, 0.6 numerical aperture) phase objective on a Agilent Biotek Lionheart FX automated microscope (Agilent Biotek, Winooski, Vermont, United States) equipped with 365 nm, 465 nm, 523 nm and 623 nm light-emitting diodes (LED; 1225007, 1225001, 1225003, 1225005) for acquiring fluorescent signals with DAPI (377/447; 1225100), GFP (469/525; 1225101), RFP (531/593; 1225103) and Cy5 (628/685; 1225105) filter cubes, respectively. Images were captured using Gen5 Image+ software, and quantification of biomass and biofilm thickness were completed either with the Gen5 Image+ software, or by importing .TIF files into BiofilmQ (76). For analysis, single channel images were analyzed by setting object threshold intensity to greater than or equal to 5000 a.u. (arbitrary units) and minimum object size to greater than 5 microns. Options selected included ‘split touching objects’ and ‘fill holes in masks’. Primary edge objects were excluded from analysis. At least five images of each sample, taken at 2500 micron increments to avoid observer bias, were acquired during each experiment.

### Isolation of Bacterial Strains from Human Saliva

During preparation of commercially available pooled human saliva, purchased from Innovative Research, 0.05 mL of saliva or diluted saliva was spread-plated on BHI agar plates before and after processing to confirm sterility prior to experimental use (incubated 37°C and 5% CO_2_ for 48 h). Agar plates from thawed-only, non-processed sample contained several different bacterial colony morphologies that were picked randomly (n = 50) using a sterile toothpick and grown overnight in BHI medium (37°C and 5% CO_2_). Cultures that grew where then streaked for single colony isolation again on a BHI agar plate. Plates that were confirmed to contain a single colony morphology were then selected, stored long-term, given a SOSUI (<u>s</u>aliva <u>O</u>hio <u>S</u>tate <u>U</u>niversity isolate) identifier number, and species identified by Sanger sequencing from Eurofins Genomics using 16S rRNA primers (F - AGA GTT TGA TCC TGG CTC AG; R – TAC GGG TAC CTT GTT ACG ACT) purchased from Integrated DNA Technologies. SOSUI isolate and species identity are listed in **Table S10**.

### Crystal Violet Biofilm Biomass Quantification

Strains were inoculated into 96 well plates and incubated for 24 h at 37ºC and 5% CO_2_. Following, medium from the biofilms was aspirated and plates were dunked into a bucket of water to remove all non-attached cells. After drying, 0.05 mL of 0.1% crystal violet (Fisher Chemical, C581) was added to each well and incubated at room temperature for 15 minutes. The crystal violet solution was then aspirated and plates were dunked into a bucket of water again to remove excess crystal violet. Plates were dried and imaged. Next, 0.2 mL of 30% acetic acid solution (RICCA Chemical, 1383032) was added to extract the bound crystal violet. Extracted crystal violet solution was diluted 1:4 with water into a new 96 well plate before the absorbance at 575 nm was recorded within an Agilent Biotek Synergy H1 multimode plate reader using Gen5 microplate reader software [v 3.11 software]. All biofilm experiments were completed with at least three biological replicates measured in technical quadruplicates.

### Fluorescent Intensity-based Competitive Growth Assays

*S. mutans* UA159 GFP-(pMZ) or GFP+ strains (pMZ-P*veg*::*gfp*) were inoculated into their respective growth mediums along with competitor species at ratios listed in **Table S9**. Cultures were plated into Costar 96 well black assay plates with a clear bottom (Corning; 3603) along with a 0.05 mL sterile mineral oil overlay. The plate was then incubated for 24 h at 37ºC in a Agilent Biotek Synergy H1 multimode plate reader with the optical density at 600 nm (OD_600 nm_) and the fluorescent intensity of GFP (excitation 485 nm, emission 528 nm, optics bottom, gain 100) recorded every 30 minutes. For data analysis, the OD_600 nm_ of a medium-only blank was subtracted from respective optical density readings and fluorescent intensity (arbitrary units; a.u.) of cultures containing *S. mutans* GFP-were subtracted from cultures containing *S. mutans* GFP+ (medium/cell background fluorescence). After plotting the resulting data points in GraphPad Prism, an Area Under the Curve (AUC) of the fluorescent intensity was calculated using built-in analysis tools. All experiments were completed with at least three biological replicates measured in technical triplicates.

### Colony Forming Unit Competitive Index Assays

1 x 10^7^ cells mL^-1^ each of *S. mutans* UA159 pMZ (kanamycin resistant) and *S. oralis* 34 ΔKX728_03760 (spectinomycin resistant; gene is transcriptionally silent in multiple conditions, verified by RNA-Seq and has no fitness defect compared to the parental) were inoculated into TYG. At this time, part of the inoculum was serially diluted and plated onto both BHI kanamycin (selection of *S. mutans*) and BHI spectinomycin (selection of *S. oralis*) agar plates. Colony forming units (CFUs) were later enumerated from these agar plates after incubation to determine the initial cell count (t_i_ = 0 h). The remaining inoculum was incubated at 37°C and 5% CO_2_ for 24 h. Resulting cultures were harvested, washed and resuspended with 1x PBS while transferring to a 5 mL polystyrene round-bottom tube. Cultures were then sonicated within a water bath sonicator for four intervals of 30 s each while resting in between for two minutes on ice to isolate single cells. Cultures were serially diluted and plated on both BHI kanamycin and spectinomycin agar plates and incubated for 48 h at 37°C and 5% CO_2_. Following CFU counting of the final cell count (t_f_ = 24 h), a competitive index was calculated using the following formula:

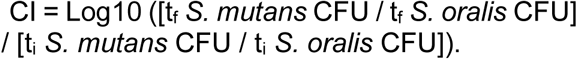

### Harvesting Cultures and RNA Isolation for RNA-Seq

Selected bacterial strains, in either monoculture or cocultures, were grown in therespective mediums until a measured optical density (OD_600 nm_) of 0.4 was reached before harvesting by centrifugation. Cell pellets were resuspended in 2 mL of RNAprotect bacterial reagent (Qiagen; 76506) and incubated at room temperature for 5 minutes prior to storage at - 80°C until further processing. For RNA isolation, the cell suspensions were thawed and RNAprotect removed by centrifugation and aspiration. Cell pellets were resuspended in a cell lysis buffer containing 0.5 mL TE buffer (Invitrogen; 12090015) along with 25 mg lysozyme (Sigma-Aldrich, L4919) and 20 units mutanolysin (Sigma-Aldrich, M9901) and incubated at 37°C for 1 h. 0.025 mL of Proteinase K solution (Qiagen; 19133) was then added to each tube and incubated at room temperature for 10 minutes. Cells suspended in lysis buffer were transferred to a 2 mL screw cap tube that contained 0.5 mL of 0.1 mm disruption beads for bacteria and 0.7 mL QIAzol™ Lysis Reagent (Qiagen; 79306). Cells were lysed via mechanical disruption using a bead beater for four rounds of 30 s each, with cells resting on ice in between each round. 0.2 mL of chloroform (Fisher Chemical, AC423550250) was added to each tube and vortexed vigorously before centrifugation at max speed for 10 minutes at 4°C. The top aqueous phase from each tube was moved into a new microcentrifuge tube (∼0.7 mL), and 0.6 mL of ice cold isopropanol was added along with 1/10^th^ volume sodium acetate solution (Invitrogen, 3 M, pH 5.2; R1181) and 1 μL GlycoBlue coprecipitant (Qiagen; AM9515). RNA was precipitated after holding overnight at -80°C and centrifugation. The RNA pellets were washed in 70% ethanol and air-dried. The resulting RNA pellets were then resuspended in RLT buffer from the RNeasy Mini Kit (Qiagen; 74524) containing 2-mercaptoethanol (Sigma-Aldrich, M3148). RNA was column purified with DNase digestion (Qiagen; 79254) according to the RNeasy Mini Kit protocol. RNA concentration was measured using a Qubit Flex Fluorometer and the Qubit RNA BR Assay Kit (Thermo Scientific; Q10210).

### RNA Sequencing and Transcriptome Analysis

RNA was sequenced through SeqCenter with their 8 M Single Reads package applied to RNA from monocultures and their 16 M Single Reads package applied to RNA from cocultures. Delivered .FASTQ files were uploaded and analyzed through Galaxy (77) with a custom pipeline (14, 78, 79) that included FASTQ Groomer [v 1.1.5], FASTQ Quality Trimmer [v 1.1.5], mapping of reads to respective genome files with Bowtie2 [v 2.3.4.3] and htseq-count [v 0.9.1] on genome features from species-specific .GFF3 files that resulted in a .CSV file containing non-normalized reads counts. All read counts were combined into a single .CSV file and uploaded to Degust (80) and edgeR analysis performed to determine log2 fold change and false discovery rates (FDR) for all genome features. The *p*-value was obtained by taking the –log10 of the FDR. All genome files for this analysis were accessed through NCBI and are listed in **Table S11**.

### Graphing and Statistics

Graphing of data was completed with GraphPad Prism [version 9.0] software. All statistical analysis was completed within GraphPad Prism using the built-in analysis tools, including Principal Component Analysis (PCA) of RNA-Seq data, one-way or two-way ANOVA with post hoc tests (Dunnett’s or Tukey’s test) for a multiple comparison, and AUC calculations.

### Data Availability

The resulting RNA-Seq raw sequencing and data files from this study are available from NCBI-GEO (Gene Expression Omnibus) under accession number GSE239483

## ACKNOLWEDGEMENTS

Several of the strains included in this manuscript were generously provided by Matthew M. Ramsey’s group at the University of Rhode Island and by Robert A. Burne at the University of Florida.

## AUTHOR CONTRIBUTIONS

AC, KD and EW each contributed equally to the original conceptual design of the project, data acquisition of growth and competitive behaviors in saliva, data analysis, data interpretation, drafted and critically revised the manuscript; LP contributed to the isolation and testing of saliva-derived isolates; JB and PB contributed through imaging and quantifying both single- and mixed-species biofilms; IS contributed with data from fluorescent-based competitive assays during supplementation with metals; DP completed various experiments, data acquisition and interpretation as well as critically revised the manuscript, JK led and contributed to the original conceptual design of the project, data interpretation, wrote and critically revised the manuscript. Listing of first authors is based on alphabetical order of the author’s last names.

