## Supplementary material for "Human Saliva Modifies Growth, Biofilm Architecture and Competitive Behaviors of Oral Streptococci": (Figure S1)

#### Table of Contents:

Supplemental Figures and Figure Legends (1-8): Pages 2 - 9

Supplemental Tables (1-11): Pages 10 – 35

* Co-first authors
### Corresponding author

**Mailing address:**
Division of Biosciences, The Ohio State University, College of Dentistry,
305 W. 12th Avenue, Postle Hall Rm 4185, Columbus, OH 43210.


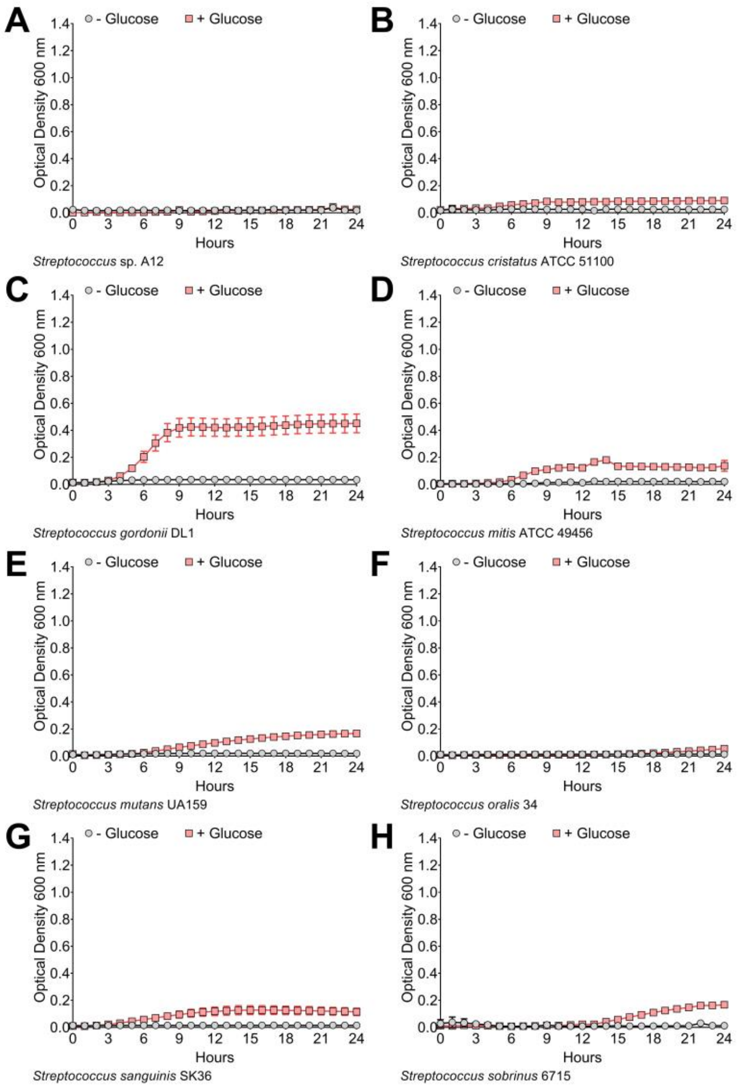


**Supplemental Figure 1. Growth of oral streptococci in pure human saliva.** Growth curves of oral streptococci species **(A)** *S*. sp. A12, **(B)** *S. cristatus* ATCC 51100, **(C)** *S. gordonii* DL1, **(D)** *S. mitis* ATCC 49456, **(E)** *S. mutans* UA159, **(F)** *S. oralis* 34, **(G)** *S. sanguinis* SK36 and **(H)** *S. sobrinus* 6715 in either 100% human saliva with no addition of carbohydrate (- Glucose; grey circles) or supplemented with 20 mM glucose (+ Glucose; red squares). Data points for optical density at 600 nm recorded every hour over a 24 h period are shown. All growth curves were completed with three biological replicates measured in technical triplicates.

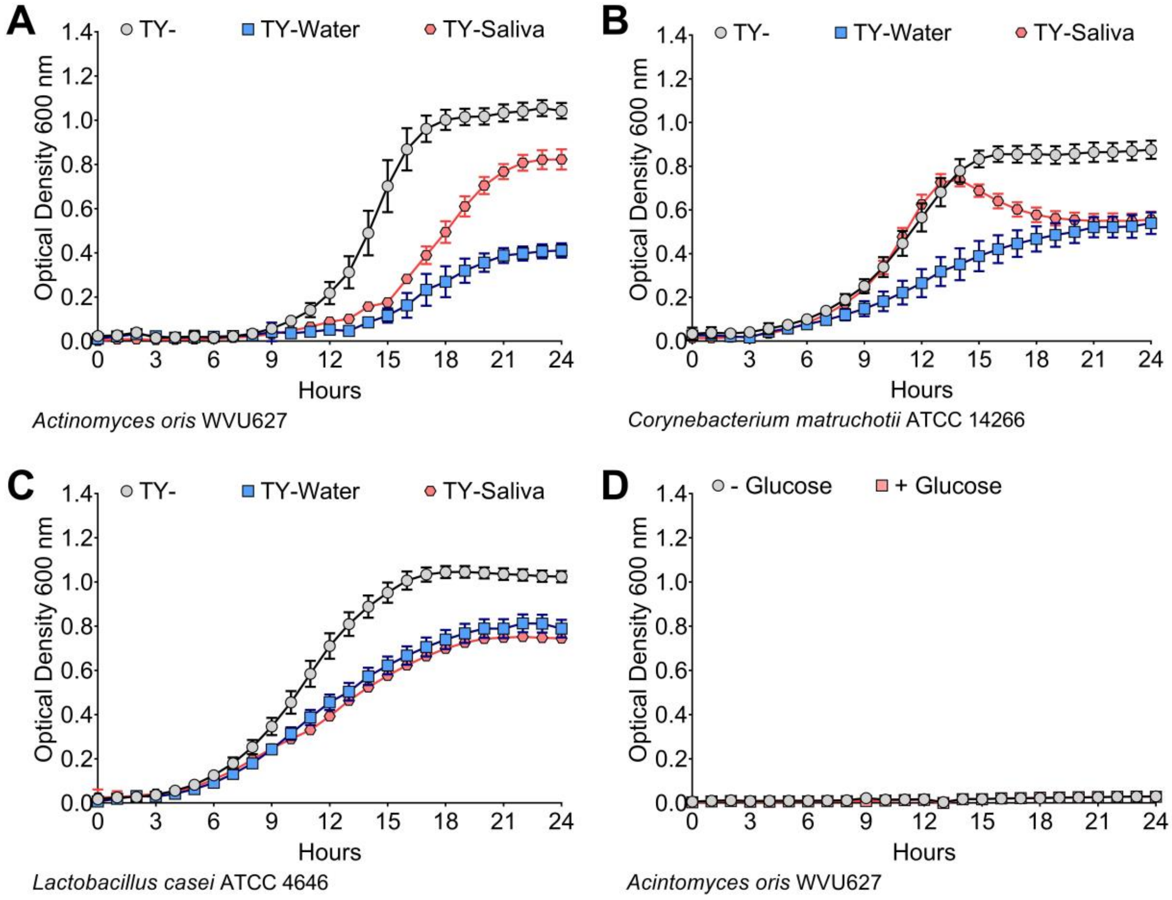


**Supplemental Figure 2. Growth of other oral bacteria in human saliva.** Growth curves of oral bacterial species **(A)** *Actinomyces oris* WVU627, **(B)** *Corynebacterium matruchotii* ATCC 14266 and **(C)** *Lactobacillus casei* ATCC 4646 in either 100% TY (TY-; grey circles), 50% TY / 50% $H_2O$ (TY-Water; blue squares), or 50% TY / 50% human saliva (TY-Saliva; red hexagons). **(D)** *Actinomyces oris* WVU627 in either 100% human saliva with no addition of carbohydrate (-Glucose; grey circles) or supplemented with 20 mM glucose (+ Glucose; red squares). Data points for optical density at 600 nm recorded every hour over a 24 h period are shown. All growth curves were completed with three biological replicates measured in technical triplicates.

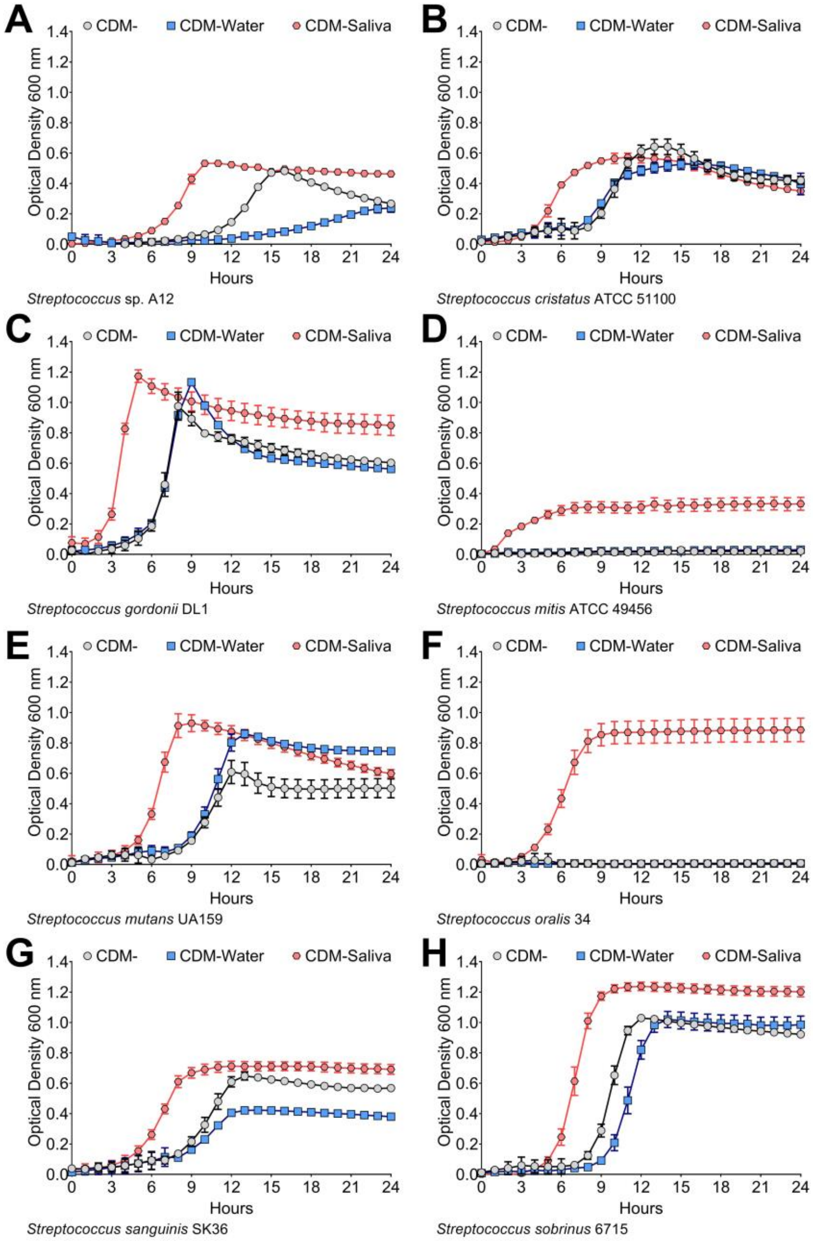


**Supplemental Figure 3. Growth of oral streptococci in human saliva with CDM medium.** Growth curves of oral streptococci species **(A)** *S*. sp. A12, **(B)** *S. cristatus* ATCC 51100, **(C)** *S. gordonii* DL1, **(D)** *S. mitis* ATCC 49456, **(E)** *S. mutans* UA159, **(F)** *S. oralis* 34, **(G)** *S. sanguinis* SK36 and **(H)** *S. sobrinus* 6715 in either 100% CDM (CDM-; grey circles), 50% CDM / 50% $H_2O$ (TY-Water; blue squares), or 50% CDM / 50% human saliva (CDM-Saliva; red hexagons). Data points for optical density at 600 nm recorded every hour over a 24 h period are shown. All growth curves were completed with three biological replicates measured in technical triplicates.

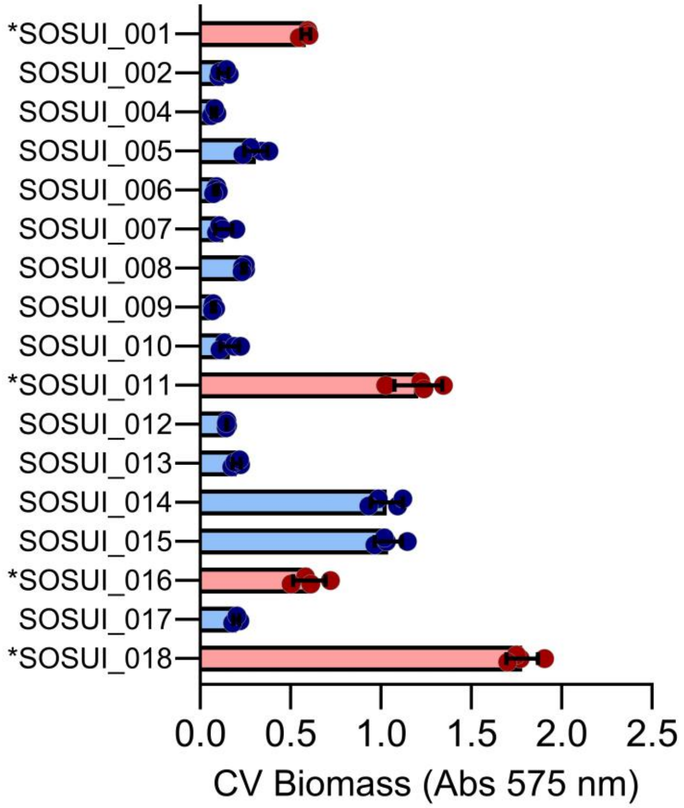


**Supplemental Figure 4. Accumulated biomass of saliva-derived bacterial isolates.** Crystal violet biomass of 17 different saliva-derived bacterial isolates, grown in TY- medium for 24 h. Data was quantified by extracting crystal violet with 30% acetic acid and measuring the absorbance at 575 nm within a Agilent Biotek® Synergy™ H1 multimode plate reader. Red bars and data points, along with asterisk next to SOSUI identifier number, denote isolates that were carried forward for further characterization. n = 4.

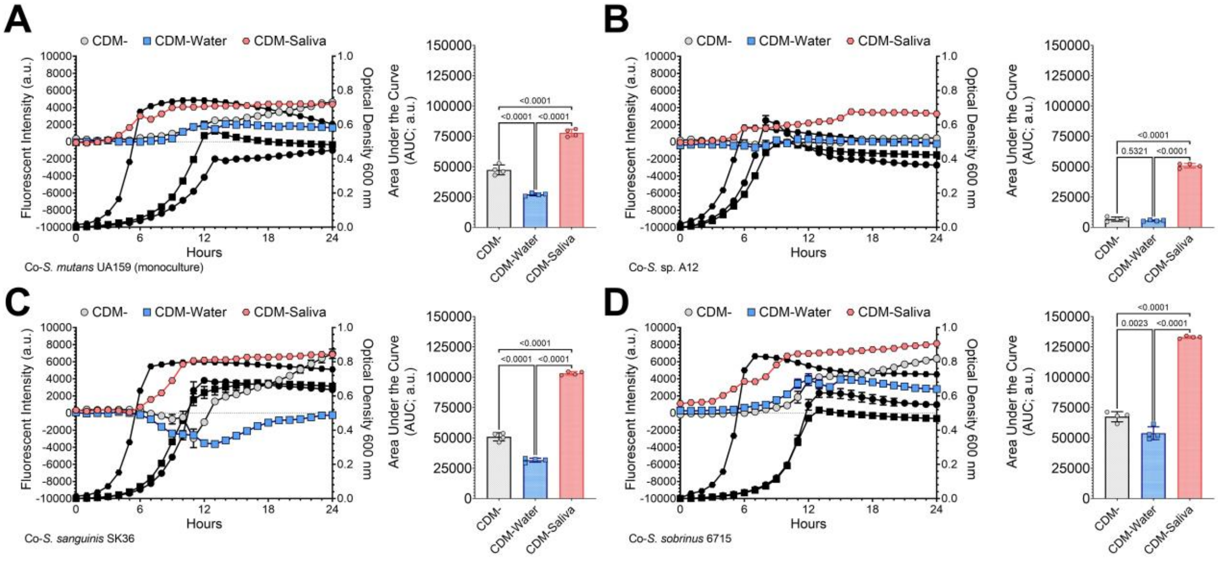


**Supplemental Figure 5. Coculture competition between *S. mutans* and oral streptococci competitors in human saliva with CDM medium.** Fluorescence-based (intensity, arbitrary units, a.u., left y-axis) growth profile of *S. mutans* in coculture competition against **(A)** *S. mutans* UA159 (monoculture growth control), **(B)** *S.* sp. A12, **(C)** *S. sanguinis* SK36 and **(D)** *S. sobrinus* 6715 in either 100% CDM (CDM-; grey circles), 50% CDM / 50% $H_2O$ (TY-Water; blue squares), or 50% CDM / 50% human saliva (CDM-Saliva; red hexagons). Growth of the entire coculture, measured by optical density at 600 nm, is shown in black using the same symbol (right y-axis). The area under the curve (AUC) of the fluorescent intensity of each condition (sum of peaks ≥ 0) was quantified and is shown in the graft on the right (n = 4). Data graphing and one-way analysis of variance with multiple comparisons was completed in GraphPad Prism software. All coculture competitions were completed with four biological replicates measured in technical triplicates.

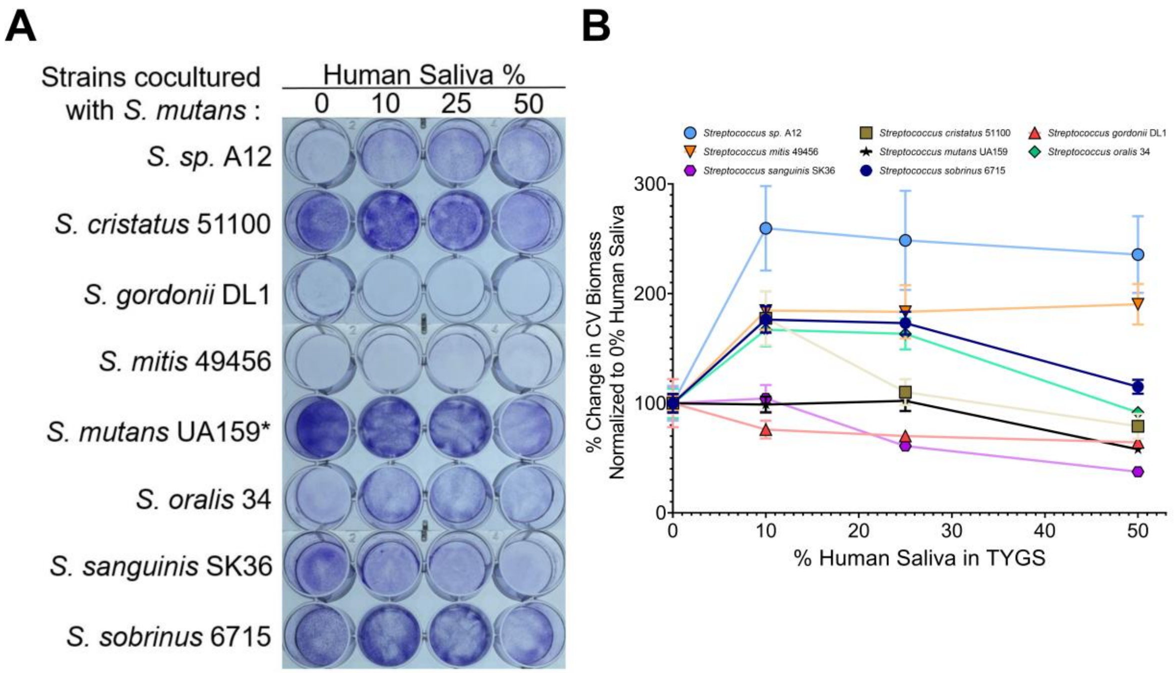


**Supplemental Figure 6. Biofilm biomass accumulation of coculture competition between *S. mutans* and oral streptococci competitors in human saliva. (A)** Representative image of a crystal violet biofilm biomass assay for eight different oral streptococci species that were cocultured with *S. mutans* (listed on y-axis) in tryptone yeast extract medium supplemented with 20 mM glucose and 5 mM sucrose containing different concentrations of human saliva (0, 10, 25 or 50%). Asterisk denotes an *S. mutans* monoculture control. **(B)** Percent (%) change in crystal violet (CV) biomass from TY- containing 0% human saliva. Data was quantified from the image shown on the left by extracting the crystal violet with 30% acetic acid and measuring the absorbance at 575 nm. n = 4; the experiment was completed with three biological replicates measured in technical quadruplicates.

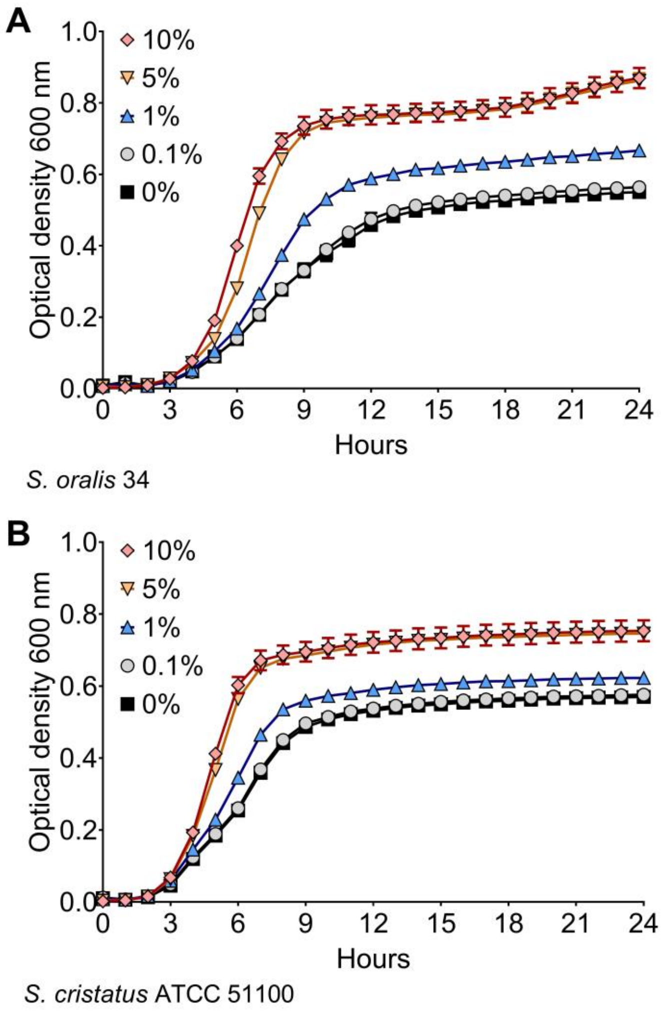


**Supplemental Figure 7. Minimum saliva concentration influencing oral streptococci growth.** Growth curves of oral bacterial species **(A)** *Streptococcus oralis* 34 and **(B)** *Streptococcus cristatus* ATCC 51100 in TY with either 0% (black squares), 0.1% (light grey circles), 1% (blue upright triangles), 5% (orange downward triangles) or 10% saliva (red diamonds). Data points for optical density at 600 nm recorded every hour over a 24 h period are shown. All growth curves were completed with three biological replicates measured in technical triplicates.

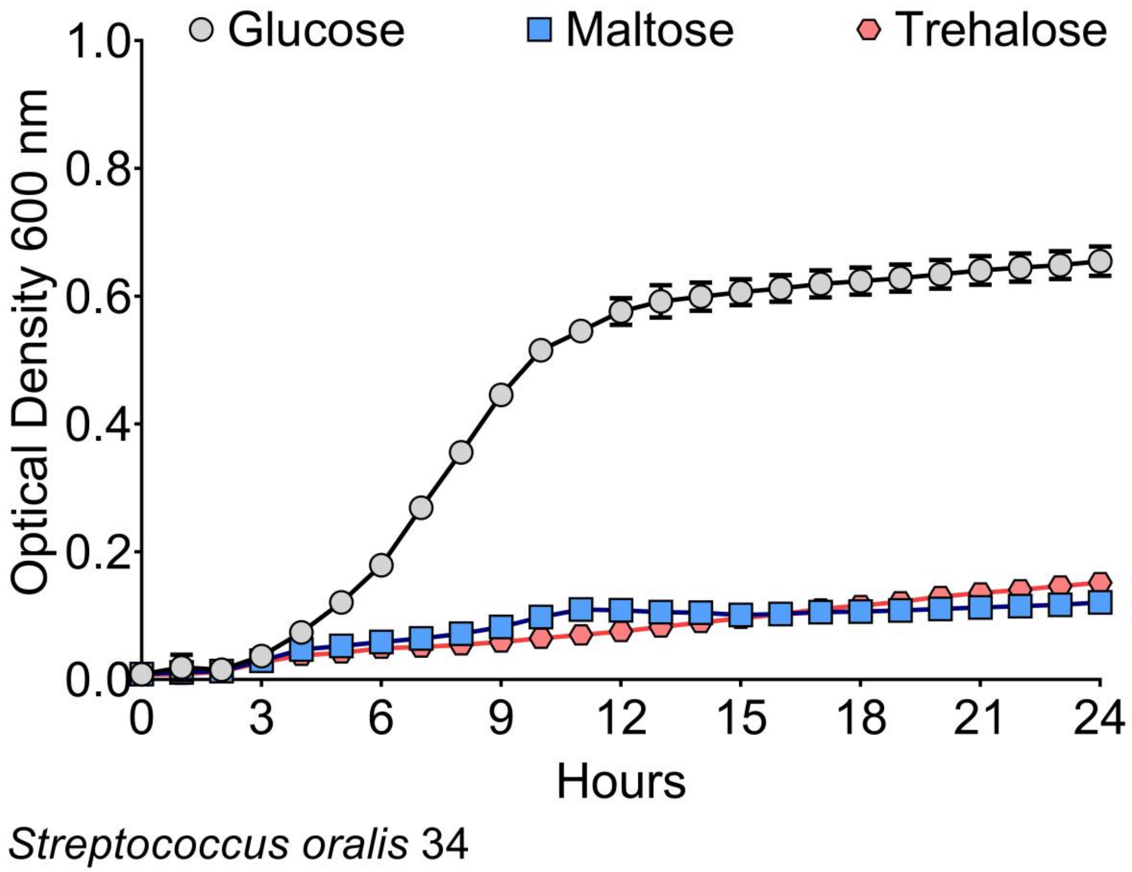


**Supplemental Figure 8. *S. oralis* growth on maltose and trehalose.** Growth curve of *Streptococcus oralis* 34 in TY- with 20 mM glucose (grey circles), 10 mM maltose (blue squares), or 10 mM trehalose (red hexagons). 10 mM was used for maltose and trehalose as both are disaccharides compared to the monosaccharide glucose. Data points for optical density at 600 nm recorded every hour over a 24 h period are shown. All growth curves were completed with three biological replicates measured in technical triplicates.

#### SUPPLEMENTAL TABLES

**Supplemental Table 1.** Differentially expressed genes* (DEGs) between *Streptococcus mutans* UA159 grown in TY- vs TY-Saliva**.** Red, upregulated genes. Blue, downregulated genes. *Differentially expressed genes = genes with ≥4 Log10 *P* value and Log2 fold change (FC) ≥ (-)1.

| Gene ID | Gene | Product | Log2 Fold Change | Fold Change | P value | Log10 P value |
|---|---|---|---|---|---|---|
| **SMU_1960c** | | fructose-specific Enzyme IIB component | 3.125208401 | 8.725322163 | 2.46E-32 | 31.61 |
| **SMU_1957** | | fructose-specific Enzyme IID component | 3.114004866 | 8.657826357 | 1.24E-29 | 28.91 |
| **SMU_1958c** | | fructose-specific Enzyme IIC component | 3.097439116 | 8.55898142 | 6.33E-20 | 19.20 |
| **SMU_1961c** | | fructose-specific Enzyme IIA component | 2.904885035 | 7.489581127 | 1.92E-16 | 15.72 |
| **SMU_1956c** | | conserved hypothetical protein | 2.716391007 | 6.572266617 | 1.04E-24 | 23.98 |
| **SMU_438c** | | (R)-2-hydroxyglutaryl-CoA dehydratase activator-related protein | 2.716257527 | 6.571658571 | 1.98E-41 | 40.70 |
| **SMU_1600** | *ptcB* | PTS system IIB component, required for cellobiose uptake and metabolism | 2.598851835 | 6.058043069 | 5.72E-12 | 11.24 |
| **SMU_1599** | *celR* | transcriptional regulator, required for cellobiose uptake and metabolism | 2.364885953 | 5.151119299 | 9.36E-25 | 24.03 |
| **SMU_1346** | *bacT* | thioesterase II-like protein | 2.152938261 | 4.447326305 | 1.31E-08 | 7.88 |
| **SMU_1538** | *glgC* | glucose-1-phosphate adenylyltransferase | 2.064996569 | 4.184329809 | 1.16E-21 | 20.93 |
| **SMU_1419** | | transcriptional regulator | 2.056051256 | 4.15846548 | 6.61E-26 | 25.18 |
| **SMU_1345c** | | peptide synthetase similar to mycA | 2.05489944 | 4.155146778 | 2.29E-10 | 9.64 |
| **SMU_1537** | *glgD* | glycogen biosynthesis protein | 1.94734188 | 3.856633042 | 2.16E-30 | 29.67 |
| **SMU_1420** | | NADPH-quinone reductase | 1.918076089 | 3.779187477 | 7.65E-19 | 18.12 |
| **SMU_1539** | *glgB* | 1,4-alpha-glucan branching enzyme | 1.914043226 | 3.768638009 | 9.91E-21 | 20.00 |
| **SMU_104** | | glycosyl hydrolase, alpha-glucosidase | 1.890923313 | 3.708725043 | 2.46E-21 | 20.61 |
| **SMU_102** | | PTS system, IID component | 1.88495483 | 3.693413601 | 8.00E-16 | 15.10 |
| **SMU_103** | | PTS system, IIA component | 1.855105274 | 3.617781491 | 1.75E-16 | 15.76 |
| **SMU_1343c** | | polyketide synthase | 1.811891503 | 3.511023137 | 4.55E-10 | 9.34 |
| **SMU_1536** | *glgA* | glycogen synthase | 1.804805862 | 3.493821401 | 1.68E-14 | 13.77 |
| **SMU_2127** | | succinic semialdehyde dehydrogenase (NAD-dependent aldehyde dehydrogenase) | 1.784204324 | 3.444284521 | 3.30E-16 | 15.48 |
| **SMU_1597c** | | conserved hypothetical protein | 1.778894889 | 3.431632092 | 2.21984E-05 | 4.65 |
| **SMU_1341c** | | gramicidin S synthase/mycosubtilin synthetase chain mycB | 1.71400026 | 3.280692236 | 7.24E-08 | 7.14 |
| **SMU_1344c** | | malonyl CoA-acyl carrier protein transacylase | 1.699962219 | 3.248924503 | 3.03E-07 | 6.52 |
| **SMU_1596** | *ptcC* | PTS system, cellobiose-specific IIC component | 1.698527876 | 3.245695991 | 1.11E-14 | 13.96 |
| **SMU_1340** | *bacA2* | bacitracin synthetase 1/ tyrocidin synthetase III | 1.690822557 | 3.228407199 | 3.45E-08 | 7.46 |
| **SMU_105** | | SCR operon transcriptional repressor | 1.683203679 | 3.211402907 | 1.30E-17 | 16.88 |
| **SMU_941c** | | conserved hypothetical protein | 1.627849022 | 3.09051876 | 5.05E-12 | 11.30 |
| **SMU_1347c** | | | 1.621577935 | 3.077114089 | 9.95E-08 | 7.00 |
| **SMU_1006** | | ABC transporter, ATP-binding protein | 1.614283559 | 3.06159523 | 1.24E-14 | 13.91 |
| **SMU_940c** | | hemolysin III-related protein | 1.594939157 | 3.020817772 | 1.20E-16 | 15.92 |
| **SMU_1047c** | | hypothetical protein | 1.593608046 | 3.018031882 | 9.44E-08 | 7.02 |
| **SMU_1342** | *bacA1* | bacitracin synthetase | 1.575621156 | 2.980637971 | 5.25E-07 | 6.28 |
| **SMU_1339** | *bacD* | bacitracin synthetase; surfactin synthetase | 1.559725943 | 2.947978377 | 2.00E-07 | 6.70 |
| **SMU_1365c** | | | 1.5352446 | 2.89837567 | 2.96E-07 | 6.53 |
| **SMU_1535** | *phsG* | glycogen phosphorylase | 1.522063198 | 2.872014828 | 3.73E-14 | 13.43 |
| **SMU_1336** | *pksD* | conserved hypothetical protein | 1.491326654 | 2.8114739 | 2.04946E-05 | 4.69 |
| **SMU_180** | | oxidoreductase, possible fumarate reductase | 1.465122047 | 2.760868258 | 3.27E-12 | 11.49 |
| **SMU_980** | *bglP* | beta-glucoside-specific EII permease | 1.462866312 | 2.756554857 | 3.12E-16 | 15.51 |
| **SMU_1895c** | | hypothetical protein | 1.455773058 | 2.743035053 | 1.08185E-06 | 5.97 |
| **SMU_1004** | *gtfB* | glucosyltransferase-I | 1.450685607 | 2.733379179 | 2.85E-11 | 10.54 |
| **SMU_1337c** | | alpha/beta superfamily hydrolases | 1.435906505 | 2.705521123 | 9.15E-07 | 6.04 |

| | | | | | | |
|---|---|---|---|---|---|---|
| **SMU_402** | *pfl* | pyruvate formate-lyase | 1.396097646 | 2.631887179 | 1.18E-09 | 8.93 |
| **SMU_78** | *fruA* | fructan hydrolase; exo-beta-D-fructosidase | 1.359901719 | 2.566676939 | 7.08E-07 | 6.15 |
| **SMU_1348c** | | | 1.359462823 | 2.565896225 | 5.00E-08 | 7.30 |
| **SMU_1366c** | | | 1.349450294 | 2.548150155 | 7.15E-08 | 7.15 |
| **SMU_1007** | | ABC transporter permease | 1.328916533 | 2.512139419 | 6.88E-12 | 11.16 |
| **SMU_1005** | *gtfC* | glucosyltransferase-SI | 1.315997534 | 2.489744215 | 5.98E-13 | 12.22 |
| **SMU_1335c** | | enoyl-acyl carrier protein(ACP) reductase | 1.296040625 | 2.455540512 | 1.36E-08 | 7.87 |
| **SMU_1077** | *pgm* | phosphoglucomutase | 1.263563113 | 2.400879694 | 1.97E-11 | 10.71 |
| **SMU_1425** | *clpB* | ATP-dependent Clp protease, ATP-binding subunit ClpB | 1.257133111 | 2.390202933 | 1.23E-08 | 7.91 |
| **SMU_1879** | | mannose PTS system component IID | 1.239548802 | 2.361246735 | 1.85E-10 | 9.73 |
| **SMU_981** | *bglB1* | beta-glucosidase, BglB protein | 1.230064405 | 2.345774616 | 9.14644E-06 | 5.04 |
| **SMU_1421** | *pdhC* | dihydrolipoamide acetyltransferase (acetoin dehydrogenase E2 component) | 1.213704933 | 2.3193249 | 5.69E-07 | 6.24 |
| **SMU_101** | | sorbose PTS system, IIC component | 1.198424043 | 2.294888473 | 3.66277E-06 | 5.44 |
| **SMU_79** | *fruB* | fructan hydrolase; exo-beta-D-fructosidase | 1.184050164 | 2.272137533 | 1.22367E-05 | 4.91 |
| **SMU_1338c** | | ABC transport macrolide permease | 1.165052848 | 2.242414297 | 1.23E-07 | 6.91 |
| **SMU_1862** | | hypothetical protein | 1.091136405 | 2.130417825 | 0.000100124 | 4.00 |
| **SMU_1601** | *bgl* | 6-phospho-beta-glucosidase, required for cellobiose uptake and metabolism | 1.038815524 | 2.05454015 | 1.19734E-05 | 4.92 |
| **SMU_197c** | | hypothetical protein | -1.000744983 | -2.001033033 | 2.62472E-05 | 4.58 |
| **SMU_1750c** | | hypothetical protein | -1.001381568 | -2.001916177 | 5.34218E-06 | 5.27 |
| **SMU_673** | | conserved hypothetical protein | -1.009451817 | -2.013146017 | 1.39E-09 | 8.86 |
| **SMU_837** | | oxidoreductase, aldo/keto reductase family | -1.012302782 | -2.017128205 | 1.96E-07 | 6.71 |
| **SMU_80** | *hrcA* | heat-inducible transcription repressor | -1.031585549 | -2.044269708 | 3.23735E-06 | 5.49 |
| **SMU_1752c** | | hypothetical protein | -1.035292613 | -2.049529297 | 4.34E-07 | 6.36 |
| **SMU_860** | *pyrAB* | carbamoyl-phosphate synthase, large subunit | -1.045932413 | -2.064700342 | 1.99877E-06 | 5.70 |
| **SMU_997** | | inorganic ion ABC transporter,ATP-binding protein; possible ferrichrome transport system | -1.048352657 | -2.06816696 | 3.53E-09 | 8.45 |
| **SMU_198c** | | conjugative transposon protein | -1.066080334 | -2.093737146 | 2.00375E-05 | 4.70 |
| **SMU_856** | *pyrR* | bifunctional protein: pyrimidine operon regulatory protein and uracil phosphoribosyltransferase | -1.091559327 | -2.131042443 | 7.05E-07 | 6.15 |
| **SMU_625** | *comEA* | competence protein | -1.09449196 | -2.135378718 | 6.61E-08 | 7.18 |
| **SMU_764** | *ahpC* | alkyl hydroperoxide reductase, subunit C | -1.103296527 | -2.148450483 | 7.43E-11 | 10.13 |
| **SMU_1987** | *comYA* | late competence protein; type II secretion system protein E | -1.110860857 | -2.15974481 | 1.98617E-05 | 4.70 |
| **SMU_644** | | competence protein CoiA | -1.114398035 | -2.165046536 | 1.33378E-06 | 5.87 |
| **SMU_82** | *dnaK* | chaperone protein, DnaK | -1.146340938 | -2.213517743 | 3.89139E-06 | 5.41 |
| **SMU_765** | | alkyl hydroperoxide reductase, subunit F | -1.159612614 | -2.233974339 | 3.89E-13 | 12.41 |
| **SMU_859** | *pyrA* | carbamoyl-phosphate synthase,small subunit | -1.164560829 | -2.241649671 | 9.44E-08 | 7.02 |
| **SMU_1980c** | | conserved hypothetical protein | -1.172035087 | -2.253293257 | 4.57148E-06 | 5.34 |
| **SMU_998** | | ABC transporter, ferrichrome-binding protein | -1.175694746 | -2.259016402 | 5.64E-12 | 11.25 |
| **SMU_671** | *citZ* | citrate synthase | -1.225160935 | -2.337815269 | 2.35E-15 | 14.63 |
| **SMU_81** | *grpE* | co-chaperone protein GrpE | -1.234718699 | -2.353354566 | 3.70462E-06 | 5.43 |
| **SMU_626** | | competence protein; possible integral membrane protein | -1.239662154 | -2.361432265 | 6.50E-09 | 8.19 |
| **SMU_836** | | hypothetical protein | -1.242616169 | -2.36627241 | 5.42E-13 | 12.27 |
| **SMU_858** | *pyrB* | aspartate transcarbamoylase | -1.249030145 | -2.376815873 | 2.32E-10 | 9.63 |
| **SMU_609** | | cell wall protein precursor | -1.264772117 | -2.402892516 | 2.11E-14 | 13.68 |
| **SMU_1984** | *comYC* | competence protein ComYC | -1.265428808 | -2.403986523 | 3.16E-07 | 6.50 |
| **SMU_838** | *gshR* | glutathione reductase | -1.266562711 | -2.405876707 | 6.43E-10 | 9.19 |
| **SMU_2042** | *dexA* | dextranase ( 1,6-alpha-glucanhydrolase ) | -1.286160206 | -2.438780991 | 3.42E-16 | 15.47 |
| **SMU_1967** | *ssb2* | single-stranded DNA-binding protein | -1.288702902 | -2.44308304 | 4.18E-13 | 12.38 |
| **SMU_672** | *idh* | isocitrate dehydrogenase | -1.298853181 | -2.460332296 | 2.21E-15 | 14.66 |
| **SMU_1983** | *comYD* | competence protein ComYD | -1.318385177 | -2.493868124 | 3.032E-05 | 4.52 |
| **SMU_499** | | late competence protein required for DNA uptake | -1.338875903 | -2.52954149 | 6.60E-07 | 6.18 |
| **SMU_670** | *citB* | aconitate hydratase; aconitase A | -1.3838615 | -2.609659353 | 1.10E-16 | 15.96 |
| **SMU_1757c** | | conserved hypothetical protein | -1.384352162 | -2.610547052 | 9.65E-10 | 9.02 |

| | | | | | | |
|---|---|---|---|---|---|---|
| **SMU_1224** | *pyrK* | dihydroorotate dehydrogenase electron transfer subunit | -1.386292679 | -2.614060768 | 2.62E-10 | 9.58 |
| **SMU_1657c** | | nitrogen regulatory protein PII | -1.391746681 | -2.623961734 | 1.84628E-06 | 5.73 |
| **SMU_857** | | xanthine/uracil permease | -1.412326465 | -2.661660316 | 2.35E-09 | 8.63 |
| **SMU_1700c** | | LrgB-like protein; possible murein hydrolase regulator | -1.41331532 | -2.663485302 | 5.77E-10 | 9.24 |
| **SMU_145** | | major facilitator superfamily transporter, efflux protein | -1.435379298 | -2.70453262 | 6.68E-12 | 11.17 |
| **SMU_933** | | amino acid ABC transporter, amino acid substrate-binding protein | -1.442062247 | -2.7170898 | 3.91292E-06 | 5.41 |
| **SMU_1400c** | | conserved hypothetical protein | -1.447811055 | -2.727938377 | 5.80E-08 | 7.24 |
| **SMU_1223** | *pyrDB* | dihydroorotate dehydrogenase | -1.510269236 | -2.848631953 | 6.71E-09 | 8.17 |
| **SMU_932** | | conserved hypothetical protein | -1.5602527 | -2.949054941 | 1.54E-08 | 7.81 |
| **SMU_936** | | amino acid ABC transporter, ATP-binding protein | -1.568137384 | -2.965216376 | 1.08E-10 | 9.97 |
| **SMU_1755c** | | conserved hypothetical protein | -1.591071018 | -3.012729234 | 3.62E-14 | 13.44 |
| **SMU_1762c** | | conserved hypothetical protein | -1.596855965 | -3.024833987 | 6.84E-12 | 11.16 |
| **SMU_1764c** | | conserved hypothetical protein | -1.621327305 | -3.076579568 | 4.74E-17 | 16.32 |
| **SMU_1758c** | | conserved hypothetical protein | -1.643948379 | -3.125199695 | 1.98E-20 | 19.70 |
| **SMU_934** | | amino acid ABC transporter, permease protein | -1.648728347 | -3.135571347 | 2.58E-09 | 8.59 |
| **SMU_1760c** | | conserved hypothetical protein | -1.657225308 | -3.154093238 | 7.84E-23 | 22.11 |
| **SMU_1761c** | | conserved hypothetical protein | -1.676345811 | -3.196173691 | 5.66E-18 | 17.25 |
| **SMU_1753c** | | conserved hypothetical protein | -1.680234525 | -3.204800442 | 2.06E-16 | 15.69 |
| **SMU_1763c** | | conserved hypothetical protein | -1.693336797 | -3.234038372 | 2.08E-18 | 17.68 |
| **SMU_935** | | amino acid ABC transporter, permease protein | -1.711596907 | -3.275231554 | 5.14E-11 | 10.29 |
| **SMU_2027** | | transcriptional regulator/repressor | -1.722083746 | -3.299125697 | 6.41E-13 | 12.19 |
| **SMU_199c** | | hypothetical protein | -1.773424077 | -3.418643719 | 4.45E-08 | 7.35 |
| **SMU_1001** | *smf* | DNA processing protein, Smf family | -1.815238068 | -3.519176976 | 9.65E-15 | 14.02 |
| **SMU_1658** | *nrgA* | ammonium transporter, NrgA protein | -1.823057735 | -3.538303337 | 5.76E-13 | 12.24 |
| **SMU_1175** | | sodium:alanine (or glycine) symporter | -1.874096916 | -3.665720822 | 8.94E-25 | 24.05 |
| **SMU_498** | *comF* | late competence protein F | -1.922658396 | -3.791210064 | 2.27E-17 | 16.64 |
| **SMU_201c** | | conserved hypothetical protein | -1.923670905 | -3.793871738 | 3.59E-13 | 12.44 |
| **SMU_1754c** | | conserved hypothetical protein | -1.935771787 | -3.825827395 | 2.74E-14 | 13.56 |
| **SMU_210c** | | hypothetical protein | -1.938198037 | -3.832266887 | 2.29E-09 | 8.64 |
| **SMU_202c** | | conserved hypothetical protein/Streptococcus-specific protein | -1.950563547 | -3.86525487 | 3.98E-12 | 11.40 |
| **SMU_205c** | | conserved hypothetical protein | -2.195354674 | -4.580022462 | 1.08E-07 | 6.97 |
| **SMU_208c** | | conserved hypothetical protein, FtsK/SpoIIIE family | -2.198727395 | -4.590742128 | 7.02E-21 | 20.15 |
| **SMU_211c** | | hypothetical protein | -2.229807246 | -4.690713044 | 1.84E-11 | 10.73 |
| **SMU_207c** | | transcriptional regulator | -2.247704899 | -4.74926711 | 4.50E-19 | 18.35 |
| **SMU_214c** | | hypothetical protein | -2.27237224 | -4.831168729 | 1.43139E-05 | 4.84 |
| **SMU_213c** | | hypothetical protein | -2.363020549 | -5.14446321 | 2.57E-07 | 6.59 |
| **SMU_209c** | | hypothetical protein | -2.739822831 | -6.679882985 | 1.09E-17 | 16.96 |
| **SMU_212c** | | hypothetical protein | -3.168050845 | -8.988315997 | 1.92138E-05 | 4.72 |
| **SMU_2037** | *treA* | trehalose-6-phosphate hydrolase | -4.353324156 | -20.44001226 | 1.73E-70 | 69.76 |
| **SMU_2038** | *pttB* | phosphotransferase system, trehalose-specific IIBC component (EIIBC-tre) | -4.462238739 | -22.04284811 | 2.02E-71 | 70.69 |

**Supplemental Table 2.** DEGs between *Streptococcus oralis* 34 grown in TY- vs TY-Saliva**.** Red, upregulated genes. Blue, downregulated genes.

| Gene ID | Gene Name | Gene Description | Log2 Fold Change | Fold Change | P value | Log10 P value |
|---|---|---|---|---|---|---|
| KX728_RS08895 | | ABC transporter, permease protein 2 (cluster 1, maltose/g3p/polyamine/iron) | 8.256835896 | 305.8829528 | 1.91E-128 | 127.72 |
| KX728_RS08900 | | ABC transporter, permease protein 1 (cluster 1, maltose/g3p/polyamine/iron) | 7.533501698 | 185.2720826 | 2.19E-142 | 141.66 |
| KX728_RS08905 | | Glycosyl hydrolase-related protein | 7.486551369 | 179.3397361 | 1.94E-156 | 155.71 |
| KX728_RS08890 | | ABC transporter, substrate-binding protein (cluster 1, maltose/g3p/polyamine/iron) | 5.219436038 | 37.25690786 | 2.12E-57 | 56.67 |
| KX728_RS08885 | | Bactriocin immunity protein BlpY | 4.969272165 | 31.32564186 | 3.59E-45 | 44.45 |
| KX728_RS00335 | | Glyoxalase family protein | 4.778266191 | 27.44109586 | 4.40E-57 | 56.36 |
| KX728_RS06730 | | Efflux ABC transporter, permease protein | 4.728386962 | 26.50857037 | 3.45E-14 | 13.46 |
| KX728_RS02155 | | Transcriptional regulator, MarR family | 4.374242951 | 20.73854754 | 4.15E-78 | 77.38 |
| KX728_RS00215 | | Phosphoribosylaminoimidazole-succinocarboxamide synthase (EC 6.3.2.6) | 4.258012065 | 19.13327675 | 7.69E-07 | 6.11 |
| KX728_RS07065 | | MBL-fold metallo-hydrolase superfamily | 3.897029597 | 14.89782274 | 3.31E-39 | 38.48 |
| KX728_RS02160 | | Glycine betaine ABC transport system, ATP-binding protein OpuAA (EC 3.6.3.32) | 3.646732158 | 12.52494322 | 2.82E-51 | 50.55 |
| KX728_RS02165 | | Glycine betaine ABC transport system, permease protein OpuAB | 3.568837583 | 11.86662347 | 5.16E-42 | 41.29 |
| KX728_RS02500 | | Transcriptional regulator, MarR family | 3.120553247 | 8.697213482 | 1.61E-44 | 43.79 |
| KX728_RS05075 | sipA | Signal peptidase I (EC 3.4.21.89) | 2.981542922 | 7.898304134 | 1.25E-32 | 31.90 |
| KX728_RS04325 | | Hydrolase, HAD superfamily | 2.932984092 | 7.636883904 | 6.57E-31 | 30.18 |
| KX728_RS05085 | pepT | Tripeptide aminopeptidase (EC 3.4.11.4) | 2.752447417 | 6.738593116 | 2.79E-39 | 38.55 |
| KX728_RS02495 | | hypothetical protein | 2.636093177 | 6.216459632 | 2.00E-35 | 34.70 |
| KX728_RS06860 | tpiA | Triosephosphate isomerase (EC 5.3.1.1) | 2.617543968 | 6.137044173 | 7.61E-39 | 38.12 |
| KX728_RS08205 | glmS | Glutamine--fructose-6-phosphate aminotransferase [isomerizing] (EC 2.6.1.16) | 2.502312465 | 5.665928771 | 1.67E-15 | 14.78 |
| KX728_RS08940 | | Zinc ABC transporter, substrate-binding lipoprotein AdcA | 2.482812698 | 5.589862122 | 2.99E-28 | 27.52 |
| KX728_RS06750 | | Fructose-bisphosphate aldolase class II (EC 4.1.2.13) | 2.479640177 | 5.577583386 | 3.22E-19 | 18.49 |
| KX728_RS05065 | srtB | NPQTN specific sortase B | 2.434867176 | 5.407145492 | 1.02E-33 | 32.99 |
| KX728_RS00250 | purD | Phosphoribosylamine--glycine ligase (EC 6.3.4.13) | 2.426052373 | 5.374208795 | 1.68E-07 | 6.78 |
| KX728_RS05070 | | EftLSL.A | 2.363064315 | 5.144619275 | 1.23E-30 | 29.91 |
| KX728_RS05080 | pitA | hypothetical protein | 2.344276732 | 5.078057503 | 1.29E-34 | 33.89 |
| KX728_RS00225 | purF | Amidophosphoribosyltransferase (EC 2.4.2.14) | 2.334835134 | 5.044933085 | 1.26E-08 | 7.90 |
| KX728_RS03030 | | Transcriptional regulator, MarR family | 2.323277691 | 5.004679533 | 1.73E-32 | 31.76 |
| KX728_RS00230 | purM | Phosphoribosylformylglycinamidine cyclo-ligase (EC 6.3.3.1) | 2.299369284 | 4.922425203 | 8.88E-08 | 7.05 |
| KX728_RS03085 | | Glycosyltransferase | 2.289775565 | 4.889800361 | 8.06E-08 | 7.09 |
| KX728_RS04430 | | Formate--tetrahydrofolate ligase (EC 6.3.4.3) | 2.263561788 | 4.801754979 | 1.03E-11 | 10.99 |
| KX728_RS06350 | | hypothetical protein | 2.205537237 | 4.612462675 | 1.03E-12 | 11.99 |
| KX728_RS00235 | purN | Phosphoribosylglycinamide formyltransferase (EC 2.1.2.2) | 2.204833236 | 4.610212452 | 2.92E-08 | 7.53 |
| KX728_RS08220 | | Prolyl-tRNA synthetase (EC 6.1.1.15), bacterial type | 2.123474652 | 4.357421432 | 3.22E-24 | 23.49 |
| KX728_RS06775 | | ABC transporter membrane-spanning permease, Pep export, Vex1 | 2.112956147 | 4.325767563 | 7.73E-20 | 19.11 |
| KX728_RS00245 | purH | IMP cyclohydrolase (EC 3.5.4.10) | 2.091420362 | 4.261674369 | 3.00E-09 | 8.52 |
| KX728_RS00240 | | acetyltransferase, GNAT family | 2.080981126 | 4.230948502 | 1.85E-07 | 6.73 |
| KX728_RS08200 | | hypothetical protein | 2.07632019 | 4.217301571 | 5.41E-13 | 12.27 |
| KX728_RS02815 | | Prophage Lp1 protein 6 | 2.074915775 | 4.213198168 | 2.45E-16 | 15.61 |
| KX728_RS07240 | | Fructokinase (EC 2.7.1.4) | 2.01387439 | 4.038653545 | 2.41E-32 | 31.62 |
| KX728_RS00255 | purE | N5-carboxyaminoimidazole ribonucleotide mutase (EC 5.4.99.18) | 2.006705609 | 4.018635171 | 6.66182E-06 | 5.18 |
| KX728_RS07205 | | hypothetical protein | 1.967841998 | 3.911825452 | 1.67E-11 | 10.78 |
| KX728_RS05095 | | Laminin-binding surface protein | 1.955258457 | 3.877853919 | 2.06E-18 | 17.69 |
| KX728_RS00695 | rrf | 5S ribosomal RNA | 1.953662263 | 3.873565842 | 3.06956E-06 | 5.51 |
| KX728_RS01405 | | hypothetical protein | 1.901835397 | 3.736883006 | 3.66E-15 | 14.44 |
| KX728_RS07220 | | hypothetical protein | 1.895836214 | 3.721376128 | 3.41E-08 | 7.47 |
| KX728_RS05100 | | Streptococcal histidine triad protein | 1.87834616 | 3.676533576 | 7.68E-17 | 16.11 |

| | | | | | | |
|---|---|---|---|---|---|---|
| KX728_RS02290 | | Glyoxalase | 1.869578993 | 3.654259259 | 6.00E-12 | 11.22 |
| KX728_RS03350 | | Collagen adhesion protein | 1.867872386 | 3.649939082 | 9.91E-20 | 19.00 |
| KX728_RS00260 | purK | N5-carboxyaminoimidazole ribonucleotide synthase (EC 6.3.4.18) | 1.865770707 | 3.644625823 | 4.12E-07 | 6.39 |
| KX728_RS09040 | | Thiamin ABC transporter ThiZ, ATPase component | 1.79285632 | 3.465002323 | 1.21E-21 | 20.92 |
| KX728_RS06770 | vex2 | ABC transporter, ATP-binding protein Vex2 | 1.792067499 | 3.463108286 | 7.23E-19 | 18.14 |
| KX728_RS00960 | rrf | 5S ribosomal RNA | 1.778282974 | 3.430176884 | 1.06763E-05 | 4.97 |
| KX728_RS01930 | | putative two-component system sensor histidine kinase, putative heat shock protein | 1.73314454 | 3.324516505 | 1.33E-13 | 12.88 |
| KX728_RS02370 | | Late competence protein ComC, processing protease | 1.684996047 | 3.215395153 | 6.65E-10 | 9.18 |
| KX728_RS06125 | | DNA translocase FtsK | 1.680360465 | 3.205080217 | 1.02562E-05 | 4.99 |
| KX728_RS01415 | | lipoprotein, putative | 1.608143597 | 3.048593083 | 1.43635E-06 | 5.84 |
| KX728_RS02025 | rrf | 5S ribosomal RNA | 1.605251073 | 3.042486946 | 2.76877E-05 | 4.56 |
| KX728_RS02285 | | Xanthine phosphoribosyltransferase (EC 2.4.2.22) | 1.601345818 | 3.034262325 | 8.07E-11 | 10.09 |
| KX728_RS02010 | | 16S ribosomal RNA | 1.594962931 | 3.020867551 | 3.47512E-05 | 4.46 |
| KX728_RS00075 | | 16S ribosomal RNA | 1.58997704 | 3.010445585 | 3.19217E-05 | 4.50 |
| KX728_RS00945 | | 16S ribosomal RNA | 1.585899461 | 3.001948988 | 3.16148E-05 | 4.50 |
| KX728_RS07210 | | Type I restriction-modification system, specificity subunit S | 1.581716918 | 2.993258587 | 3.37E-10 | 9.47 |
| KX728_RS02505 | | Antiholin-like protein LrgA | 1.58029368 | 2.990307153 | 6.75E-16 | 15.17 |
| KX728_RS00680 | | 16S ribosomal RNA | 1.568028844 | 2.964993298 | 4.42358E-05 | 4.35 |
| KX728_RS02360 | trpB | Tryptophan synthase beta chain (EC 4.2.1.20) | 1.558899721 | 2.946290573 | 8.25E-07 | 6.08 |
| KX728_RS08325 | rplQ | LSU ribosomal protein L17p | 1.556417173 | 2.941225042 | 5.62E-10 | 9.25 |
| KX728_RS02710 | | hypothetical protein | 1.52889263 | 2.885642607 | 1.94E-08 | 7.71 |
| KX728_RS05960 | tRNA-Arg | | 1.527952553 | 2.883762901 | 4.08E-12 | 11.39 |
| KX728_RS07680 | | Response regulator CsrR | 1.527494865 | 2.882848185 | 1.36E-10 | 9.87 |
| KX728_RS06780 | tRNA-Leu | | 1.526335786 | 2.880533001 | 8.73E-15 | 14.06 |
| KX728_RS02275 | | hypothetical protein | 1.523792843 | 2.875460148 | 6.17357E-06 | 5.21 |
| KX728_RS06590 | thiD | Hydroxymethylpyrimidine kinase (EC 2.7.1.49) | 1.506196501 | 2.840601592 | 2.80E-11 | 10.55 |
| KX728_RS01665 | | Putative large secreted protein SCO0341 | 1.500944299 | 2.830279044 | 5.66E-12 | 11.25 |
| KX728_RS08495 | nrdD | Ribonucleotide reductase of class III (anaerobic), large subunit (EC 1.17.4.2) | 1.498949885 | 2.826369107 | 8.63E-07 | 6.06 |
| KX728_RS01400 | | hypothetical protein | 1.465712147 | 2.761997756 | 1.09E-13 | 12.96 |
| KX728_RS05365 | | Uncharacterized protease YrrO | 1.461680761 | 2.754290557 | 4.01E-17 | 16.40 |
| KX728_RS01935 | | FIG01117953: hypothetical protein | 1.454634129 | 2.74087043 | 7.45E-13 | 12.13 |
| KX728_RS03740 | | Putative cI repressor, metallo-proteinase motif (ACLAME 174) | 1.441165499 | 2.71540144 | 6.33E-12 | 11.20 |
| KX728_RS07235 | | Immunoreactive protein Se23.5 (Fragment) | 1.438124605 | 2.709683979 | 4.01E-20 | 19.40 |
| KX728_RS08350 | infA | Translation initiation factor 1 | 1.437081298 | 2.707725138 | 3.53E-12 | 11.45 |
| KX728_RS02340 | | Anthranilate synthase, amidotransferase component (EC 4.1.3.27) | 1.423669999 | 2.682670748 | 1.77E-07 | 6.75 |
| KX728_RS09070 | | Choline binding protein D | 1.421223108 | 2.678124646 | 3.61E-11 | 10.44 |
| KX728_RS04245 | | hypothetical protein | 1.41212115 | 2.661281554 | 4.55E-07 | 6.34 |
| KX728_RS05945 | cas2 | CRISPR-associated protein Cas2 | 1.409618797 | 2.656669564 | 1.4888E-05 | 4.83 |
| KX728_RS00590 | | hypothetical protein | 1.398316609 | 2.635938314 | 3.74E-12 | 11.43 |
| KX728_RS08990 | hslO | 33 kDa chaperonin HslO | 1.391079545 | 2.622748634 | 3.86493E-05 | 4.41 |
| KX728_RS00540 | | Lj965 prophage protein | 1.389678932 | 2.620203624 | 7.06E-10 | 9.15 |
| KX728_RS05370 | | Transcriptional regulator in cluster with beta-lactamase, GntR family | 1.385217442 | 2.61211324 | 1.38E-16 | 15.86 |
| KX728_RS00450 | | FIG01119883: hypothetical protein | 1.378234706 | 2.599500993 | 2.48409E-05 | 4.60 |
| KX728_RS05360 | | hypothetical protein | 1.372585759 | 2.589342413 | 2.64E-13 | 12.58 |
| KX728_RS00545 | | hypothetical protein | 1.364010335 | 2.573996933 | 5.24E-12 | 11.28 |
| KX728_RS06580 | | Copper-exporting ATPase (EC 3.6.3.4) | 1.335820227 | 2.524189507 | 1.73E-12 | 11.76 |
| KX728_RS02280 | | Xanthine permease | 1.315920561 | 2.489611382 | 4.90347E-06 | 5.31 |
| KX728_RS06555 | | 6-phospho-beta-glucosidase (EC 3.2.1.86) | 1.308455402 | 2.476762274 | 8.09E-15 | 14.09 |
| KX728_RS07675 | | Choline-binding protein | 1.29527603 | 2.454239478 | 1.07E-14 | 13.97 |
| KX728_RS02350 | trpC | Indole-3-glycerol phosphate synthase (EC 4.1.1.48) | 1.294348971 | 2.452662919 | 2.68312E-06 | 5.57 |
| KX728_RS03580 | | hypothetical protein | 1.285295601 | 2.437319871 | 1.47E-07 | 6.83 |
| KX728_RS07580 | | Acyl-CoA dehydrogenase, short-chain specific (EC 1.3.8.1) | 1.285141259 | 2.437059135 | 3.38E-14 | 13.47 |

| | | | | | | |
|---|---|---|---|---|---|---|
| KX728_RS05305 | | TPR-repeat-containing protein | 1.283624424 | 2.434498183 | 1.70E-09 | 8.77 |
| KX728_RS02005 | tRNA-Glu | | 1.282320911 | 2.432299543 | 5.55E-07 | 6.26 |
| KX728_RS00265 | | hypothetical protein | 1.280676407 | 2.429528585 | 6.29E-10 | 9.20 |
| KX728_RS07215 | | hypothetical protein | 1.265216669 | 2.403633059 | 6.24E-10 | 9.20 |
| KX728_RS07245 | | PTS system, sucrose-specific IIB component (EC 2.7.1.211) | 1.257156395 | 2.39024151 | 2.34E-11 | 10.63 |
| KX728_RS00315 | | PTS system, galactosamine-specific IIA component | 1.253445882 | 2.384101871 | 1.36E-12 | 11.87 |
| KX728_RS05460 | | Arginine pathway regulatory protein ArgR, repressor of arg regulon | 1.248279482 | 2.375579491 | 4.66E-09 | 8.33 |
| KX728_RS02695 | | hypothetical protein | 1.246256026 | 2.372249949 | 4.53E-07 | 6.34 |
| KX728_RS07905 | | Undecaprenyl-diphosphatase (EC 3.6.1.27) | 1.243028297 | 2.366948468 | 2.63E-15 | 14.58 |
| KX728_RS00535 | | hypothetical protein | 1.238962989 | 2.360288135 | 5.35E-12 | 11.27 |
| KX728_RS00310 | | PTS system, galactosamine-specific IID component | 1.237515612 | 2.357921375 | 1.84E-12 | 11.73 |
| KX728_RS06585 | | Negative transcriptional regulator-copper transport operon | 1.232803679 | 2.350232817 | 7.10E-07 | 6.15 |
| KX728_RS02365 | trpA | Tryptophan synthase alpha chain (EC 4.2.1.20) | 1.228618785 | 2.343425263 | 2.05461E-05 | 4.69 |
| KX728_RS00275 | | Serine/threonine kinase | 1.222334022 | 2.333238884 | 8.10E-10 | 9.09 |
| KX728_RS05695 | | MutT/Nudix family protein | 1.22212941 | 2.332907993 | 2.22181E-05 | 4.65 |
| KX728_RS02380 | | Putative oxidoreductase | 1.203613207 | 2.303157716 | 2.10E-14 | 13.68 |
| KX728_RS02400 | | hypothetical protein | 1.191265792 | 2.283530078 | 1.78E-09 | 8.75 |
| KX728_RS01475 | | hypothetical protein | 1.174462922 | 2.257088397 | 2.67E-09 | 8.57 |
| KX728_RS01410 | | putative lipoprotein | 1.169564485 | 2.249437816 | 3.78524E-05 | 4.42 |
| KX728_RS07120 | ugpC | Maltodextrin ABC transporter, ATP-binding protein MsmX | 1.168126493 | 2.247196828 | 7.43013E-06 | 5.13 |
| KX728_RS05060 | srtB | NPQTN specific sortase B | 1.165593414 | 2.243254669 | 1.30E-13 | 12.89 |
| KX728_RS03585 | | Alanyl-tRNA synthetase (EC 6.1.1.7) | 1.163368542 | 2.239797869 | 2.10E-07 | 6.68 |
| KX728_RS04125 | | hypothetical protein | 1.156705203 | 2.229476825 | 1.46316E-05 | 4.83 |
| KX728_RS00340 | | hypothetical protein | 1.154992705 | 2.226831976 | 5.28E-10 | 9.28 |
| KX728_RS04675 | | Uracil-DNA glycosylase, family 1 (EC 3.2.2.27) | 1.153176269 | 2.224030032 | 1.58E-10 | 9.80 |
| KX728_RS05410 | ccdA2 | Cytochrome c-type biogenesis protein CcdA homolog | 1.148109628 | 2.216233098 | 6.60E-13 | 12.18 |
| KX728_RS01575 | | hypothetical protein | 1.131264374 | 2.190506315 | 1.52822E-05 | 4.82 |
| KX728_RS04535 | | Phosphoglucomutase (EC 5.4.2.2) | 1.129335436 | 2.187579483 | 1.69E-14 | 13.77 |
| KX728_RS08815 | sdaAA | L-serine dehydratase, alpha subunit (EC 4.3.1.17) | 1.128305234 | 2.18601793 | 6.83E-14 | 13.17 |
| KX728_RS00285 | | beta-glycosyl hydrolase | 1.12045467 | 2.174154809 | 9.41E-07 | 6.03 |
| KX728_RS06355 | | transcription regulator | 1.119665828 | 2.172966341 | 2.91689E-05 | 4.54 |
| KX728_RS01600 | | hypothetical protein | 1.11913932 | 2.172173467 | 4.72465E-06 | 5.33 |
| KX728_RS03755 | | Phage protein | 1.116849449 | 2.168728491 | 2.17706E-06 | 5.66 |
| KX728_RS03210 | | Transcriptional regulator, Xre family | 1.112041423 | 2.161512864 | 1.76477E-05 | 4.75 |
| KX728_RS03735 | | hypothetical protein | 1.109366287 | 2.157508566 | 5.95E-10 | 9.23 |
| KX728_RS00500 | ilvD | Dihydroxy-acid dehydratase (EC 4.2.1.9) | 1.106551853 | 2.153303764 | 3.00E-08 | 7.52 |
| KX728_RS02705 | | hypothetical protein | 1.105517707 | 2.151760795 | 2.1599E-06 | 5.67 |
| KX728_RS04665 | | 2-haloalkanoic acid dehalogenase (EC 3.8.1.2) | 1.086874514 | 2.124133607 | 3.04E-09 | 8.52 |
| KX728_RS07320 | | UPF0348 protein family | 1.07136845 | 2.101425703 | 1.58E-10 | 9.80 |
| KX728_RS04250 | | GMP reductase (EC 1.7.1.7) | 1.066127408 | 2.093805466 | 1.00203E-06 | 6.00 |
| KX728_RS00885 | yajC | Protein translocase subunit YajC | 1.053991296 | 2.076266015 | 6.42E-11 | 10.19 |
| KX728_RS00305 | | PTS system, galactosamine-specific IIC component | 1.053465817 | 2.075509905 | 5.19E-09 | 8.28 |
| KX728_RS02660 | | Sialic acid utilization regulator, RpiR family | 1.047891138 | 2.067505457 | 4.32E-10 | 9.36 |
| KX728_RS00595 | | Lead, cadmium, zinc and mercury transporting ATPase (EC 3.6.3.3) (EC 3.6.3.5) | 1.037685609 | 2.052931669 | 4.41E-08 | 7.36 |
| KX728_RS00330 | | Aldose 1-epimerase (EC 5.1.3.3) | 1.037495078 | 2.052660565 | 5.49E-13 | 12.26 |
| KX728_RS02725 | | hypothetical protein | 1.036978658 | 2.051925937 | 6.09E-08 | 7.22 |
| KX728_RS07095 | tpx | Thiol peroxidase, Tpx-type (EC 1.11.1.15) | 1.033287883 | 2.046683304 | 4.40E-09 | 8.36 |
| KX728_RS01460 | | Putative secretion accessory protein EsaB/YukD | 1.015432808 | 2.021509254 | 2.91806E-06 | 5.53 |
| KX728_RS05955 | cas9 | CRISPR-associated endonuclease Cas9 | 1.010464951 | 2.014560246 | 4.35422E-05 | 4.36 |
| KX728_RS05465 | | hypothetical protein | 1.005747445 | 2.007983543 | 2.71E-10 | 9.57 |
| KX728_RS06955 | | Tributyrin esterase | -1.002657568 | -2.003687567 | 4.28658E-06 | 5.37 |

| | | | | | | |
|---|---|---|---|---|---|---|
| KX728_RS01320 | | Substrate-specific component RibU of riboflavin ECF transporter | -1.018869428 | -2.026330398 | 3.19E-10 | 9.50 |
| KX728_RS08795 | | Argininosuccinate synthase (EC 6.3.4.5) | -1.042920745 | -2.060394715 | 9.63687E-06 | 5.02 |
| KX728_RS07520 | fabF | 3-oxoacyl-[acyl-carrier-protein] synthase, KASII (EC 2.3.1.179) | -1.047446791 | -2.066868769 | 2.16E-11 | 10.67 |
| KX728_RS03710 | | Methylthioadenosine deaminase (EC 3.5.4.31) | -1.047782672 | -2.067350023 | 4.63E-11 | 10.33 |
| KX728_RS03885 | | Substrate-specific component BioY of biotin ECF transporter | -1.051240676 | -2.072311209 | 1.01568E-06 | 5.99 |
| KX728_RS06015 | | Carbamoyl-phosphate synthase small chain (EC 6.3.5.5) | -1.052169837 | -2.073646302 | 9.12441E-05 | 4.04 |
| KX728_RS07295 | fmt | Methionyl-tRNA formyltransferase (EC 2.1.2.9) | -1.053783496 | -2.075966979 | 4.30E-12 | 11.37 |
| KX728_RS04110 | | O-acetylhomoserine sulfhydrylase (EC 2.5.1.49) | -1.055333605 | -2.078198708 | 2.02E-07 | 6.69 |
| KX728_RS04715 | | Lipoate-protein ligase A | -1.096329202 | -2.13809981 | 2.97E-11 | 10.53 |
| KX728_RS04350 | | Glutamine ABC transporter, substrate-binding protein GlnH | -1.118167286 | -2.170710432 | 3.69E-10 | 9.43 |
| KX728_RS01780 | glnA | Glutamine synthetase type I (EC 6.3.1.2) | -1.125186725 | -2.181297767 | 9.89E-09 | 8.00 |
| KX728_RS06080 | | Beta-lactamase class C-like and penicillin binding proteins (PBPs) superfamily | -1.127203707 | -2.184349497 | 3.84E-14 | 13.42 |
| KX728_RS06075 | | Permease of the drug/metabolite transporter (DMT) superfamily | -1.128360916 | -2.186102302 | 4.69849E-06 | 5.33 |
| KX728_RS01550 | | glycosyl hydrolase family 29 (alpha-L-fucosidase) | -1.155113011 | -2.227017679 | 2.70E-13 | 12.57 |
| KX728_RS00655 | | Glutamine amidotransferase, class I | -1.17119556 | -2.251982412 | 5.55E-09 | 8.26 |
| KX728_RS00580 | glgP | Maltodextrin phosphorylase (EC 2.4.1.1) | -1.182036775 | -2.268968807 | 3.89E-12 | 11.41 |
| KX728_RS05680 | | Lysyl aminopeptidase (EC 3.4.11.15) | -1.191003785 | -2.283115404 | 4.43E-08 | 7.35 |
| KX728_RS06990 | | ABC transporter, permease protein PebF (cluster 3, basic aa/glutamine/opines) | -1.197992591 | -2.294202266 | 4.82089E-06 | 5.32 |
| KX728_RS07300 | | Helicase PriA essential for oriC/DnaA-independent DNA replication | -1.209939935 | -2.313280054 | 3.02E-15 | 14.52 |
| KX728_RS05675 | ciaR | Two component system response regulator CiaR | -1.222609988 | -2.333685241 | 2.01E-07 | 6.70 |
| KX728_RS08130 | | ABC transporter, permease protein (cluster 3, basic aa/glutamine/opines) | -1.230732817 | -2.346861685 | 5.99575E-05 | 4.22 |
| KX728_RS03595 | | Putative ribosomal RNA large subunit methyltransferase YwbD | -1.233657768 | -2.35162459 | 1.79E-12 | 11.75 |
| KX728_RS02295 | xth | Exodeoxyribonuclease III (EC 3.1.11.2) | -1.247532223 | -2.374349352 | 5.80E-21 | 20.24 |
| KX728_RS04405 | | Substrate-specific component PanT of predicted pantothenate ECF transporter | -1.251747127 | -2.381296272 | 4.60E-18 | 17.34 |
| KX728_RS08075 | recA | RecA protein | -1.255790122 | -2.387978955 | 3.24E-07 | 6.49 |
| KX728_RS02240 | | Hydrolase, HAD superfamily | -1.280330544 | -2.428946214 | 7.80E-08 | 7.11 |
| KX728_RS07305 | rpoZ | DNA-directed RNA polymerase omega subunit (EC 2.7.7.6) | -1.282392443 | -2.432420145 | 2.46E-15 | 14.61 |
| KX728_RS08140 | | ABC transporter, substrate-binding protein (cluster 3, basic aa/glutamine/opines) | -1.296857237 | -2.456930822 | 3.84855E-05 | 4.41 |
| KX728_RS02855 | | probable esterase of alpha/beta hydrolase superfamily, YBBA B. subtilis ortholog | -1.343306006 | -2.537320926 | 1.02496E-05 | 4.99 |
| KX728_RS00575 | malQ | 4-alpha-glucanotransferase (amylomaltase) (EC 2.4.1.25) | -1.348854476 | -2.547098012 | 2.97E-14 | 13.53 |
| KX728_RS08135 | | ABC transporter, permease protein (cluster 3, basic aa/glutamine/opines) | -1.39308266 | -2.626392728 | 1.09822E-05 | 4.96 |
| KX728_RS05780 | | Late competence protein ComEA, DNA receptor | -1.526538785 | -2.880938343 | 7.07E-07 | 6.15 |
| KX728_RS02845 | | Vitamin B12 ABC transporter, ATP-binding protein BtuD | -1.541059246 | -2.91008087 | 5.16E-07 | 6.29 |
| KX728_RS02860 | | Acyl-ACP:1-acyl-sn-glycerol-3-phosphate acyltransferase (EC 2.3.1.n4) | -1.62817947 | -3.09122672 | 5.67E-11 | 10.25 |
| KX728_RS01540 | | hypothetical protein | -1.66834866 | -3.178505667 | 4.73E-14 | 13.32 |
| KX728_RS07100 | | Mn-dependent transcriptional regulator MntR | -1.692286353 | -3.231684486 | 2.24E-20 | 19.65 |
| KX728_RS03850 | | Hydrolase, alpha/beta fold family | -1.716531597 | -3.286453556 | 4.13E-13 | 12.38 |
| KX728_RS06240 | pdxS | Pyridoxal 5'-phosphate synthase (glutamine hydrolyzing) | -1.752155503 | -3.368614887 | 1.14E-10 | 9.94 |
| KX728_RS02475 | queE | 7-carboxy-7-deazaguanine synthase (EC 4.3.99.3) | -1.810144673 | -3.506774526 | 1.24E-12 | 11.91 |
| KX728_RS01530 | | CI-like repressor, phage associated | -1.818559742 | -3.527288897 | 5.16E-16 | 15.29 |
| KX728_RS02850 | | Vitamin B12 ABC transporter, substrate-binding protein BtuF | -1.838093452 | -3.575372241 | 5.99E-10 | 9.22 |
| KX728_RS06245 | | NADH peroxidase (EC 1.11.1.1) | -1.843522445 | -3.588852039 | 2.73E-12 | 11.56 |
| KX728_RS04645 | | hypothetical protein | -1.843590788 | -3.589022053 | 1.62E-08 | 7.79 |
| KX728_RS01735 | | Iron compound ABC uptake transporter permease protein PiuB | -1.912929633 | -3.76573018 | 3.09E-08 | 7.51 |
| KX728_RS01545 | | hypothetical protein | -1.977967067 | -3.939375845 | 9.61E-25 | 24.02 |
| KX728_RS01730 | | Iron compound ABC uptake transporter permease protein PiuC | -1.987874004 | -3.966520497 | 1.32E-09 | 8.88 |
| KX728_RS01720 | | Iron compound ABC uptake transporter substrate-binding protein PiuA | -2.082138505 | -4.234344076 | 4.50E-12 | 11.35 |
| KX728_RS04630 | | Exopolysaccharide biosynthesis protein | -2.09190019 | -4.263092001 | 2.64E-08 | 7.58 |
| KX728_RS01535 | | Putative UV-damage repair protein UvrX | -2.146079642 | -4.426233749 | 5.48E-29 | 28.26 |
| KX728_RS02610 | | Sialidase (EC 3.2.1.18) | -2.17396383 | -4.512615412 | 1.24E-07 | 6.91 |
| KX728_RS04635 | | hypothetical protein | -2.319922173 | -4.993052835 | 2.30E-08 | 7.64 |
| KX728_RS08880 | | Mundticin KS immunity protein | -2.387082171 | -5.230983326 | 1.54E-28 | 27.81 |

| | | | | | | |
|---|---|---|---|---|---|---|
| KX728_RS02490 | trxA | Thioredoxin | -2.459023386 | -5.498443909 | 9.69E-27 | 26.01 |
| KX728_RS02485 | queC | 7-cyano-7-deazaguanine synthase (EC 6.3.4.20) | -2.487020961 | -5.606191247 | 3.38E-34 | 33.47 |
| KX728_RS01725 | | Iron compound ABC uptake transporter ATP-binding protein PiuD | -2.489647425 | -5.616406762 | 9.81E-11 | 10.01 |
| KX728_RS02480 | queD | 6-carboxy-5,6,7,8-tetrahydropterin synthase (EC 4.1.2.50) | -2.500909178 | -5.660420289 | 1.13E-26 | 25.95 |
| KX728_RS02230 | galR | Galactose operon repressor, GalR-LacI family of transcriptional regulators | -3.095317533 | -8.546404096 | 2.06E-60 | 59.69 |

**Supplemental Table 3.** DEGs in *Streptococcus mutans* UA159 grown in coculture with *Streptococcus oralis* 34 between TY- vs TY-Saliva**.** Red, upregulated genes. Blue, downregulated genes.

| Geneid | Gene | Product | Log2 Fold Change | Fold Change | P value | Log10 P value |
|---|---|---|---|---|---|---|
| SMU_1284c | | conserved hypothetical protein | 2.864765735 | 7.284175769 | 1.17251E-05 | 4.93 |
| SMU_1185 | *mtlA1* | mannitol PTS EII | 2.687742709 | 6.443045171 | 1.31918E-05 | 4.88 |
| SMU_910 | *gtfD* | glucosyltransferase-S | 2.321953509 | 5.000088079 | 2.29E-07 | 6.64 |
| SMU_2147c | | conserved hypothetical protein | 2.217019708 | 4.649319943 | 4.10E-07 | 6.39 |
| SMU_184 | *sloC* | ABC transporter element, iron predicted binding protein | 2.161151134 | 4.472715933 | 4.56724E-06 | 5.34 |
| SMU_1419 | | transcriptional regulator | 2.118394979 | 4.342106101 | 7.34861E-06 | 5.13 |
| SMU_1821c | | glutamyl-tRNA (Gln) amidotransferase subunit C | 2.113995439 | 4.328884892 | 8.13911E-06 | 5.09 |
| SMU_2146c | | conserved hypothetical protein | 2.057888028 | 4.163763215 | 1.5251E-05 | 4.82 |
| SMU_182 | *sloA* | ABC transporter, ATP-binding protein, iron and/or manganese | 1.960192048 | 3.891137733 | 3.40417E-05 | 4.47 |
| SMU_1942c | | amino acid ABC transporter substrate-binding protein | 1.908858238 | 3.755117987 | 4.05E-07 | 6.39 |
| SMU_1124 | *pdp* | pyrimidine-nucleoside phosphorylase | 1.838245937 | 3.575750158 | 7.69339E-05 | 4.11 |
| SMU_183 | *sloB* | manganese ABC transporter permease element | 1.79349964 | 3.466547767 | 8.47E-07 | 6.07 |
| SMU_672 | *idh* | isocitrate dehydrogenase | 1.790525605 | 3.459409033 | 2.70E-07 | 6.57 |
| SMU_673 | | conserved hypothetical protein | 1.751072008 | 3.366085934 | 4.5925E-06 | 5.34 |
| SMU_1822 | *gatA* | aspartyl-tRNA synthetase | 1.74146718 | 3.343750448 | 1.45E-07 | 6.84 |
| SMU_1390 | | conserved hypothetical protein | 1.740080333 | 3.340537681 | 6.52908E-06 | 5.19 |
| SMU_496 | *cysK* | cysteine synthetase A | 1.730419604 | 3.318243145 | 1.04182E-05 | 4.98 |
| SMU_866 | | conserved hypothetical protein | 1.609873822 | 3.052251457 | 7.15922E-05 | 4.15 |
| SMU_186 | *sloR* | metal-dependent transcriptional regulator (probable DtxR homolog) | 1.601348324 | 3.034267596 | 3.27397E-06 | 5.48 |
| SMU_2019 | *rl29* | 50s ribosomal protein L29 | 1.588652005 | 3.007681927 | 1.97765E-05 | 4.70 |
| SMU_831 | | conserved hypothetical protein | 1.519315384 | 2.866549881 | 2.61E-07 | 6.58 |
| SMU_80 | *hrcA* | heat-inducible transcription repressor | 1.512070503 | 2.852190814 | 2.6366E-06 | 5.58 |
| SMU_957 | | 50S ribosomal protein L10 | 1.492572133 | 2.813902095 | 1.52956E-06 | 5.82 |
| SMU_1565 | *malQ* | 4-alpha-glucanotransferase | 1.490340757 | 2.809553274 | 7.76E-07 | 6.11 |
| SMU_2033c | | conserved hypothetical protein | 1.489422581 | 2.807765756 | 7.21E-07 | 6.14 |
| SMU_671 | *citZ* | citrate synthase | 1.47120111 | 2.772526231 | 9.82167E-06 | 5.01 |
| SMU_1514 | *rnc* | ribonuclease III | 1.414706449 | 2.666054826 | 4.49571E-05 | 4.35 |
| SMU_1877 | *ptnA* | mannose PTS system component IIAB | 1.393707593 | 2.62753065 | 1.0636E-06 | 5.97 |
| SMU_81 | *grpE* | co-chaperone protein GrpE | 1.392729337 | 2.62574959 | 2.22627E-05 | 4.65 |
| SMU_670 | *citB* | aconitate hydratase; aconitase A | 1.389473725 | 2.619830955 | 4.56973E-06 | 5.34 |
| SMU_459 | | amino acid ABC transporter, substrate-binding protein | 1.389224124 | 2.619377737 | 5.68259E-05 | 4.25 |
| SMU_832 | | hypothetical protein | 1.389180444 | 2.619298432 | 7.3093E-06 | 5.14 |
| SMU_941c | | conserved hypothetical protein | 1.387069038 | 2.615467853 | 8.86223E-05 | 4.05 |
| SMU_1568 | *malX* | maltose / maltodextrin-binding protein | 1.34247369 | 2.535857524 | 6.02695E-05 | 4.22 |
| SMU_848 | | conserved hypothetical protein | 1.332083426 | 2.517659931 | 1.31263E-05 | 4.88 |
| SMU_649 | | conserved hypothetical protein | 1.323733186 | 2.503129942 | 1.60787E-05 | 4.79 |
| SMU_830 | *rgpF* | polysaccharide biosynthesis protein | 1.307383943 | 2.474923518 | 1.58302E-06 | 5.80 |
| SMU_1288 | *rl19* | 50S ribosomal protein L19 | 1.306813066 | 2.47394438 | 5.98806E-06 | 5.22 |
| SMU_1611c | | multi-drug resistance efflux pump, major facilitator superfamily | 1.288638285 | 2.44297362 | 4.10658E-05 | 4.39 |
| SMU_1939c | | amino acid ABC transport ATP-binding protein | 1.27611169 | 2.421853655 | 5.2669E-05 | 4.28 |
| SMU_2166 | *rplW* | Ribosomal protein L23 | 1.260287659 | 2.395434988 | 6.36E-07 | 6.20 |
| SMU_1564 | *glgP* | glycogen phosphorylase | 1.253671352 | 2.384474497 | 4.09483E-06 | 5.39 |
| SMU_1006 | | ABC transporter, ATP-binding protein | 1.244172492 | 2.36882643 | 4.68768E-05 | 4.33 |
| SMU_1206c | | conserved hypothetical protein | 1.199750359 | 2.296999208 | 5.00521E-05 | 4.30 |
| SMU_774 | | hydrolase, haloacid dehalogenase-like | 1.192529915 | 2.285531836 | 1.20796E-05 | 4.92 |
| SMU_532 | *trpE* | anthranilate synthase, component I | 1.181248558 | 2.267729493 | 5.69817E-05 | 4.24 |

| | | | | | | |
|---|---|---|---|---|---|---|
| SMU_500 | | ribosome-associated protein | 1.174133117 | 2.256572479 | 4.86656E-05 | 4.31 |
| SMU_2038 | *pttB* | phosphotransferase system, trehalose-specific IIBC component (EIIBC-tre) | 1.165040227 | 2.24239468 | 3.2798E-05 | 4.48 |
| SMU_697 | | translation initiation factor IF-3 | 1.162657511 | 2.238694259 | 8.01234E-06 | 5.10 |
| SMU_460 | | amino acid ABC transporter, permease | 1.155721302 | 2.227956866 | 0.000100403 | 4.00 |
| SMU_1177c | | amino acid ABC transporter, amino acid-binding protein | 1.138828479 | 2.202021384 | 1.86363E-05 | 4.73 |
| SMU_2037 | *treA* | trehalose-6-phosphate hydrolase | 1.132342899 | 2.192144499 | 4.60745E-06 | 5.34 |
| SMU_2047 | *ptsG* | PTS system, enzyme II, A component | 1.13162311 | 2.191051067 | 4.57303E-06 | 5.34 |
| SMU_1879 | | mannose PTS system component IID | 1.124384971 | 2.180085884 | 1.87161E-06 | 5.73 |
| SMU_1178c | | amino acid ABC transporter, ATP-binding protein | 1.110674425 | 2.159465736 | 2.69842E-05 | 4.57 |
| SMU_1878 | *ptnC* | mannose PTS system component IIC | 1.09053183 | 2.12952524 | 6.03348E-06 | 5.22 |
| SMU_829 | *rgpE* | glycosyltransferase | 1.082436703 | 2.117609691 | 4.61152E-06 | 5.34 |
| SMU_2011 | *rl6* | 50S ribosomal protein L6 (BL10) | 1.068378089 | 2.097074461 | 2.44814E-06 | 5.61 |
| SMU_611 | | ATP-dependent RNA helicase/DEAD family | 1.068151812 | 2.096745576 | 9.30E-07 | 6.03 |
| SMU_1940c | | peptidase, possible succinyl-diaminopimelic descuccinlyase | 1.060221174 | 2.085251178 | 5.34848E-05 | 4.27 |
| SMU_1545c | | conserved hypothetical protein | 1.057014686 | 2.080621713 | 7.75756E-05 | 4.11 |
| SMU_1179c | | amino acid ABC transporter, permease | 1.050786727 | 2.071659251 | 1.94071E-05 | 4.71 |
| SMU_834 | | glycosyltransferase | 1.041950871 | 2.059010048 | 1.26181E-05 | 4.90 |
| SMU_835 | | conserved hypothetical protein (possible membrane protein) | 1.04095088 | 2.057583359 | 1.01757E-05 | 4.99 |
| SMU_1820c | | glutamyl-tRNA(Gln) amidotransferase subunit A | 1.029583841 | 2.041435295 | 2.22683E-06 | 5.65 |
| SMU_1745c | | transcriptional regulator, MarR family | 1.027782423 | 2.038887853 | 3.3796E-05 | 4.47 |
| SMU_16 | | amino acid permease | 1.013141913 | 2.018301791 | 8.56546E-05 | 4.07 |
| SMU_2016 | *rl24* | 50S ribosomal protein L24 | -1.015809181 | -2.022036699 | 1.90674E-05 | 4.72 |
| SMU_463 | *trxB* | thioredoxin reductase (NADPH) | -1.016601059 | -2.023146874 | 5.07695E-05 | 4.29 |
| SMU_2112 | *gbpA* | glucan-binding protein | -1.029235991 | -2.040943142 | 1.30022E-06 | 5.89 |
| SMU_2079c | | conserved hypothetical protein | -1.032872797 | -2.046094526 | 7.55527E-06 | 5.12 |
| SMU_1833 | *recG* | ATP-dependent DNA helicase | -1.046465727 | -2.065463731 | 8.46767E-06 | 5.07 |
| SMU_2049c | | conserved hypothetical protein | -1.052901872 | -2.074698753 | 3.33456E-05 | 4.48 |
| SMU_2042 | *dexA* | dextranase ( 1,6-alpha-glucanhydrolase ) | -1.05670073 | -2.080168982 | 4.2679E-06 | 5.37 |
| SMU_610 | *spaP* | cell surface antigen | -1.059815047 | -2.084664251 | 2.59719E-05 | 4.59 |
| SMU_1040c | | oxidoreductase, short-chain dehydrogenase/reductase | -1.090446745 | -2.129399653 | 1.71252E-05 | 4.77 |
| SMU_991 | | conserved hypothetical protein (similar to ribonucleotide reductase alpha subunit) | -1.094497043 | -2.13538624 | 1.7052E-05 | 4.77 |
| SMU_527 | | conserved hypothetical protein | -1.11732101 | -2.169437479 | 6.47E-07 | 6.19 |
| SMU_1297 | | conserved hypothetical protein, DHH family | -1.127412277 | -2.184665311 | 1.82803E-05 | 4.74 |
| SMU_1451 | *aldB* | alpha-acetolactate decarboxylase | -1.128909175 | -2.186933232 | 2.50707E-06 | 5.60 |
| SMU_401c | | acetyltransferase | -1.180203902 | -2.266088024 | 3.87243E-06 | 5.41 |
| SMU_275 | | L-ribulose 5-phosphate 4-epimerase | -1.187284845 | -2.277237631 | 7.18223E-05 | 4.14 |
| SMU_400 | | beta-lactamase family protein | -1.215470963 | -2.322165768 | 3.31E-07 | 6.48 |
| SMU_1679c | | conserved hypothetical protein | -1.227141582 | -2.341027013 | 1.33207E-05 | 4.88 |
| SMU_32 | *purF* | amidophosphoribosyltransferase | -1.22793267 | -2.342311044 | 7.15652E-05 | 4.15 |
| SMU_1958c | | fructose-specific Enzyme IIC component | -1.233164175 | -2.350820158 | 4.67705E-06 | 5.33 |
| SMU_1382 | *leuC* | alpha-isopropylmalate isomerase large subunit | -1.237425117 | -2.357773475 | 1.27088E-06 | 5.90 |
| SMU_51 | *purK* | phosphoribosylaminoimidazole carboxylase, ATPase subunit | -1.241897924 | -2.365094655 | 2.17794E-06 | 5.66 |
| SMU_270 | *sgaT* | ribulose monophosphate PTS pathway enzyme IIC | -1.242895767 | -2.366731044 | 1.19347E-05 | 4.92 |
| SMU_1706 | | conserved hypothetical protein | -1.255439715 | -2.387399024 | 4.57324E-05 | 4.34 |
| SMU_1627 | *rl11* | 50S ribosomal protein L11 | -1.267974674 | -2.408232487 | 5.1827E-05 | 4.29 |
| SMU_1038c | | response regulator | -1.268379127 | -2.408907718 | 5.2196E-06 | 5.28 |
| SMU_1452 | *alsS* | alpha-acetolactate synthase | -1.272133177 | -2.415184121 | 5.26695E-06 | 5.28 |
| SMU_119 | *adh* | alcohol dehydrogenase class III | -1.279926654 | -2.428266313 | 2.02047E-06 | 5.69 |
| SMU_360 | *gapC* | glyceraldehyde-3-phosphate dehydrogenase; plasmin receptor | -1.286747966 | -2.439774762 | 1.06514E-05 | 4.97 |
| SMU_34 | *purM* | phosphoribosylformylglycinamide cyclo-ligase (AIRS) | -1.29060831 | -2.446311821 | 2.60862E-05 | 4.58 |
| SMU_1383 | *leuB* | 3-isopropylmalate dehydrogenase | -1.2963552 | -2.456075994 | 1.55236E-06 | 5.81 |

| | | | | | | |
|---|---|---|---|---|---|---|
| SMU_118c | | esterase | -1.29795673 | -2.458803988 | 1.97388E-05 | 4.70 |
| SMU_1384 | *leuA* | 2-isopropylmalate synthase | -1.307213682 | -2.474631456 | 1.83479E-06 | 5.74 |
| SMU_54 | | amino acid racemase | -1.309747293 | -2.478981136 | 8.97818E-06 | 5.05 |
| SMU_595 | *pyrD* | dihydroorotate dehydrogenase (dihydroorotate oxidase) | -1.309932978 | -2.479300219 | 3.65453E-06 | 5.44 |
| SMU_1602 | | NAD(P)H-flavin oxidoreductase | -1.314597144 | -2.487328652 | 1.41896E-05 | 4.85 |
| SMU_1626 | *rl1* | 50S ribosomal protein L1 | -1.337102571 | -2.526434139 | 3.79822E-05 | 4.42 |
| SMU_1574c | | conserved hypothetical protein | -1.349207056 | -2.547720574 | 2.91273E-06 | 5.54 |
| SMU_2025 | *rl3* | 50S ribosomal protein L3 | -1.400632949 | -2.640173883 | 2.93434E-05 | 4.53 |
| SMU_924 | *tpx* | thiol peroxidase | -1.457748424 | -2.746793441 | 4.17E-07 | 6.38 |
| SMU_48 | *purD* | phosphoribosylamine-glycine ligase | -1.465597859 | -2.761778963 | 8.39E-07 | 6.08 |
| SMU_37 | *purH* | phosphoribosylaminoimidazolecarboxamide formyltransferase / IMP cyclohydrolase | -1.473950843 | -2.777815621 | 5.28E-07 | 6.28 |
| SMU_1494 | *lacC* | tagatose-6-phosphate kinase | -1.537455785 | -2.902821348 | 4.98274E-05 | 4.30 |
| SMU_1603 | *lguL* | lactoylglutathione lyase | -1.601389508 | -3.034354215 | 4.15733E-06 | 5.38 |
| SMU_2023c | | 30S ribosomal protein S19, C-terminal fragment | -1.635224231 | -3.106358268 | 4.21977E-06 | 5.37 |
| SMU_30 | *purL* | phosphoribosylformylglycinamidine synthase | -1.648450531 | -3.134967597 | 1.97E-07 | 6.71 |
| SMU_618 | | hypothetical protein | -1.669354142 | -3.180721688 | 5.70E-08 | 7.24 |
| SMU_1489 | *lacX* | aldose 1-epimerase | -1.676524695 | -3.196570017 | 1.43533E-06 | 5.84 |
| SMU_528c | | conserved hypothetical protein | -1.702185142 | -3.253934345 | 1.41693E-05 | 4.85 |
| SMU_629 | *sod* | superoxide dismutase | -1.712633614 | -3.277585952 | 2.89E-07 | 6.54 |
| SMU_59 | *purB* | adenylosuccinate lyase | -1.732280524 | -3.32252608 | 1.37E-07 | 6.86 |
| SMU_2157 | *guaB* | inosine-5'- monophosphate dehydrogenase | -1.755695695 | -3.376891193 | 6.30E-09 | 8.20 |
| SMU_1073 | *fthS* | formate--tetrahydrofolate ligase | -1.758916077 | -3.384437505 | 2.47E-07 | 6.61 |
| SMU_764 | *ahpC* | alkyl hydroperoxide reductase, subunit C | -1.762514521 | -3.392889679 | 4.25E-07 | 6.37 |
| SMU_714 | | translation elongation factor Tu | -1.788151322 | -3.453720471 | 2.75341E-06 | 5.56 |
| SMU_1493 | *lacD* | tagatose-1,6-bisphosphate aldolase | -1.793865226 | -3.467426318 | 2.12756E-06 | 5.67 |
| SMU_1860 | *rs6* | 30S ribosomal protein S6 | -1.799706119 | -3.48149299 | 1.41E-07 | 6.85 |
| SMU_2167 | *rplB* | Ribosomal protein L2 | -1.807711751 | -3.50086578 | 3.76443E-06 | 5.42 |
| SMU_1858 | *rs18* | 30S ribosomal protein S18 | -1.813039262 | -3.513817498 | 1.01189E-05 | 4.99 |
| SMU_2018 | *rs17* | 30S ribosomal protein S17 | -1.864842389 | -3.642281402 | 1.49493E-06 | 5.83 |
| SMU_1491 | *lacE* | PTS system, lactose-specific component IIBC | -1.864939664 | -3.642526995 | 2.01595E-06 | 5.70 |
| SMU_1490 | *lacG* | phospho-beta-D-galactosidase | -1.880126378 | -3.681073046 | 3.96E-07 | 6.40 |
| SMU_99 | *fbaA* | fructose-bisphosphate aldolase | -1.926543422 | -3.801433152 | 1.59623E-06 | 5.80 |
| SMU_925 | | bacteriocin immunity protein | -1.93523247 | -3.824397469 | 5.44E-08 | 7.26 |
| SMU_2002 | *rs11* | 30S ribosomal protein S11 | -1.951094272 | -3.866677046 | 1.13632E-06 | 5.94 |
| SMU_1247 | *eno* | enolase | -1.956921697 | -3.882327159 | 2.05225E-06 | 5.69 |
| SMU_765 | | alkyl hydroperoxide reductase, subunit F | -1.959132314 | -3.888280541 | 1.38E-07 | 6.86 |
| SMU_2021 | *rs3* | 30S ribosomal protein S3 | -1.965554899 | -3.905628963 | 2.27903E-06 | 5.64 |
| SMU_651c | | ABC transporter, periplasmic substrate-binding protein | -1.980339848 | -3.945860213 | 9.29738E-05 | 4.03 |
| SMU_1492 | *lacF* | PTS system, cellobiose-specific IIA component | -1.996062568 | -3.989098005 | 5.54667E-05 | 4.26 |
| SMU_698 | | 50S ribosomal protein L35 | -2.009564796 | -4.02660735 | 1.2338E-06 | 5.91 |
| SMU_1296 | | glutathione S-transferase | -2.041912403 | -4.117910296 | 3.46E-07 | 6.46 |
| SMU_2104a | | 50S ribosomal protein L32 | -2.052211659 | -4.147412831 | 3.92336E-06 | 5.41 |
| SMU_2015 | *rl5* | 50S ribosomal protein L5 | -2.074661873 | -4.212456746 | 8.59E-07 | 6.07 |
| SMU_2020 | *rl16* | 50S ribosomal protein L16 | -2.105466714 | -4.303369484 | 2.63253E-06 | 5.58 |
| SMU_2017 | *rl14* | 50S ribosomal protein L14 | -2.225337637 | -4.676203247 | 9.17E-07 | 6.04 |
| SMU_2010 | *rl18* | 50S ribosomal protein L18 | -2.233194223 | -4.701738243 | 2.43118E-06 | 5.61 |
| SMU_1037c | | histidine kinase | -2.566059376 | -5.921896912 | 1.59E-09 | 8.80 |
| SMU_1682c | | conserved hypothetical protein (possible intracellular protease) | -2.770532219 | -6.823595938 | 6.02E-07 | 6.22 |
| SMU_175 | | | -3.112026508 | -8.645962073 | 6.45071E-05 | 4.19 |
| SMU_996 | | ABC transporter, permease protein;possible ferrichrome transport system | -3.128731765 | -8.746657277 | 6.79E-10 | 9.17 |
| SMU_995 | | ferrichrome ABC transporter (permease) | -3.192363805 | -9.14107479 | 4.65E-10 | 9.33 |
| SMU_998 | | ABC transporter, ferrichrome-binding protein | -3.211129551 | -9.260753292 | 2.58E-09 | 8.59 |

| | | | | | |
|---|---|---|---|---|---|
| SMU_997 | inorganic ion ABC transporter,ATP-binding protein; possible ferrichrome transport system | -3.248657148 | -9.504805786 | 2.34E-09 | 8.63 |
| SMU_1764c | conserved hypothetical protein | -3.36334353 | -10.29123009 | 2.44676E-06 | 5.61 |
| SMU_176 | hypothetical protein | -3.368657465 | -10.32920608 | 4.86245E-05 | 4.31 |
| SMU_1470c | conserved hypothetical protein | -3.497170161 | -11.29153847 | 5.21E-10 | 9.28 |
| SMU_2027 | transcriptional regulator/repressor | -3.525700096 | -11.51705616 | 7.81E-09 | 8.11 |
| SMU_1761c | conserved hypothetical protein | -3.723177386 | -13.20651024 | 6.42E-08 | 7.19 |
| SMU_1763c | conserved hypothetical protein | -3.840962276 | -14.32995601 | 6.22E-08 | 7.21 |
| SMU_1750c | hypothetical protein | -3.856652662 | -14.48665555 | 4.91E-09 | 8.31 |
| SMU_1762c | conserved hypothetical protein | -3.860899882 | -14.52936637 | 4.36E-08 | 7.36 |
| SMU_217c | conserved hypothetical protein; Streptococcus-specific protein | -3.982812376 | -15.81051411 | 2.42436E-06 | 5.62 |
| SMU_1752c | hypothetical protein | -4.142507808 | -17.66115522 | 3.46E-08 | 7.46 |
| SMU_932 | conserved hypothetical protein | -4.177333571 | -18.09267186 | 2.76E-08 | 7.56 |
| SMU_1760c | conserved hypothetical protein | -4.191425154 | -18.27025867 | 6.82E-09 | 8.17 |
| SMU_1757c | conserved hypothetical protein | -4.215661513 | -18.57977999 | 2.57E-08 | 7.59 |
| SMU_961 | macrophage infectivity potentiator-related protein | -4.409685709 | -21.25434226 | 3.58E-09 | 8.45 |
| SMU_1755c | conserved hypothetical protein | -4.413690095 | -21.31341838 | 1.36E-09 | 8.87 |
| SMU_1758c | conserved hypothetical protein | -4.442797452 | -21.74779836 | 8.35E-09 | 8.08 |
| SMU_936 | amino acid ABC transporter, ATP-binding protein | -4.478416261 | -22.29141451 | 1.44E-07 | 6.84 |
| SMU_935 | amino acid ABC transporter, permease protein | -4.506700334 | -22.73275032 | 4.98E-09 | 8.30 |
| SMU_933 | amino acid ABC transporter, amino acid substrate-binding protein | -4.577178462 | -23.87085708 | 1.12E-07 | 6.95 |
| SMU_191c | phage-related integrase | -4.602476667 | -24.29313319 | 2.09E-07 | 6.68 |
| SMU_962 | acyl-CoA dehydrogenase | -4.619620473 | -24.58353491 | 1.37E-09 | 8.86 |
| SMU_195c | hypothetical protein | -4.642631101 | -24.97877984 | 4.14E-07 | 6.38 |
| SMU_1754c | conserved hypothetical protein | -4.647998181 | -25.07187837 | 5.13E-10 | 9.29 |
| SMU_934 | amino acid ABC transporter, permease protein | -4.650404314 | -25.11372825 | 1.68E-08 | 7.77 |
| SMU_1753c | conserved hypothetical protein | -4.683232169 | -25.6917306 | 6.47E-09 | 8.19 |
| SMU_196c | immunogenic secreted protein (transfer protein) | -4.964494532 | -31.22207535 | 3.52E-10 | 9.45 |
| SMU_194c | conserved hypothetical protein, phage-related | -5.015345414 | -32.34218881 | 2.21488E-06 | 5.65 |
| SMU_215c | hypothetical protein | -5.25254536 | -38.12182708 | 2.95E-07 | 6.53 |
| SMU_216c | hypothetical protein | -5.691541616 | -51.68026692 | 6.16E-07 | 6.21 |
| SMU_200c | hypothetical protein | -6.131075572 | -70.08702938 | 3.03E-07 | 6.52 |
| SMU_198c | conjugative transposon protein | -6.300662506 | -78.82943376 | 1.01E-09 | 8.99 |
| SMU_213c | hypothetical protein | -6.351137642 | -81.63622928 | 2.28839E-06 | 5.64 |
| SMU_212c | hypothetical protein | -6.354243385 | -81.81215994 | 2.53255E-06 | 5.60 |
| SMU_199c | hypothetical protein | -6.435304082 | -86.54053186 | 1.12E-09 | 8.95 |
| SMU_206c | hypothetical protein | -6.4607037 | -88.07762733 | 4.41E-07 | 6.36 |
| SMU_197c | hypothetical protein | -6.474903618 | -88.94882347 | 2.26E-11 | 10.65 |
| SMU_204c | hypothetical protein | -6.705046802 | -104.3326431 | 5.18E-09 | 8.29 |
| SMU_214c | hypothetical protein | -6.921377529 | -121.2110595 | 1.74E-07 | 6.76 |
| SMU_202c | conserved hypothetical protein/Streptococcus-specific protein | -6.940149763 | -122.7985541 | 1.31E-09 | 8.88 |
| SMU_205c | conserved hypothetical protein | -7.104055285 | -137.5731659 | 1.71E-08 | 7.77 |
| SMU_210c | hypothetical protein | -7.273954601 | -154.7670562 | 1.12E-07 | 6.95 |
| SMU_211c | hypothetical protein | -7.62219843 | -197.0200207 | 1.21E-09 | 8.92 |
| SMU_208c | conserved hypothetical protein, FtsK/SpoIIIE family | -7.680653949 | -205.1668662 | 2.80E-11 | 10.55 |
| SMU_207c | transcriptional regulator | -7.751483451 | -215.490947 | 3.65E-10 | 9.44 |
| SMU_209c | hypothetical protein | -7.789318636 | -221.2170289 | 2.13E-09 | 8.67 |
| SMU_201c | conserved hypothetical protein | -8.018553668 | -259.3135293 | 1.38E-08 | 7.86 |

**Supplemental Table 4.** *Streptococcus mutans* UA159 DEGs during coculture with *Streptococcus oralis* 34 in either TY-, TY-Saliva, or both. Light grey,TY- only; Red, TY-Saliva; Dark blue, both.

| TY | | | TY-Saliva | | |
|---|---|---|---|---|---|
| **Gene ID** | **Gene** | **Product** | **Geneid** | **Gene** | **Product** |
| | | | SMU_05 | | conserved hypothetical protein |
| | | | SMU_102 | | PTS system, IID component |
| | | | SMU_103 | | PTS system, IIA component |
| | | | SMU_104 | | glycosyl hydrolase, alpha-glucosidase |
| | | | SMU_105 | | SCR operon transcriptional repressor |
| | | | SMU_106c | | |
| SMU_1036 | | conserved hypothetical protein | | | |
| SMU_1037c | | histidine kinase | | | |
| SMU_1063 | *opuAa* | amino acid ABC transporter, ATP-binding protein | | | |
| | | | SMU_1073 | *fthS* | formate--tetrahydrofolate ligase |
| | | | SMU_1077 | *pgm* | phosphoglucomutase |
| | | | SMU_1090 | | conserved hypothetical protein |
| | | | SMU_1091 | *wapE* | hypothetical protein (possible cell wall protein) |
| SMU_1093 | | ABC transporter permease protein | | | |
| | | | SMU_110 | *mutR* | transcriptional regulator |
| SMU_1131c | | hypothetical protein | | | |
| | | | SMU_1175 | | sodium:alanine (or glycine) symporter |
| SMU_1177c | | amino acid ABC transporter, amino acid-binding protein | | | |
| SMU_1178c | | amino acid ABC transporter, ATP-binding protein | | | |
| SMU_1189c | | conserved hypothetical protein | | | |
| SMU_1197 | | conserved hypothetical protein | | | |
| | | | SMU_120 | | 50S ribosomal protein L28 |
| SMU_1223 | *pyrDB* | dihydroorotate dehydrogenase | | | |
| SMU_1246c | | transcriptional regulator (TetR/AcrR family) | | | |
| SMU_1254 | | phosphatase | | | |
| | | | SMU_1260c | | conserved hypothetical protein |
| | | | SMU_1262c | | hypothetical protein |
| SMU_132 | | amino acid amidohydrolase (hippurate amidohydrolase) | | | |
| SMU_1326 | *rf2* | peptide chain release factor | | | |
| | | | SMU_1396 | *gbpC* | glucan-binding protein C |
| | | | SMU_1400c | | conserved hypothetical protein |
| | | | SMU_1403c | | conserved hypothetical protein |
| | | | SMU_1425 | *clpB* | ATP-dependent Clp protease, ATP-binding subunit ClpB |
| SMU_1470c | | conserved hypothetical protein | | | |
| | | | SMU_148 | *adhE* | alcohol-acetaldehyde dehydrogenase |
| | | | SMU_1488c | | conserved hypothetical protein |
| | | | SMU_1489 | *lacX* | aldose 1-epimerase |
| | | | SMU_149 | | transposase fragment (IS605/IS200-like) |
| | | | SMU_1490 | *lacG* | phospho-beta-D-galactosidase |
| | | | SMU_1491 | *lacE* | PTS system, lactose-specific component IIBC |
| | | | SMU_1492 | *lacF* | PTS system, cellobiose-specific IIA component |
| | | | SMU_1493 | *lacD* | tagatose-1,6-bisphosphate aldolase |
| | | | SMU_1494 | *lacC* | tagatose-6-phosphate kinase |
| | | | SMU_1495 | *lacB* | galactose-6-phosphate isomerase |
| | | | SMU_1496 | *lacA* | galactose-6-phosphate isomerase |
| SMU_150 | | non-lantibiotic mutacin IV A | | | |

| | | | | | |
|---|---|---|---|---|---|
| SMU_151 | | non-lantibiotic mutacin IV B | | | |
| SMU_152 | | hypothetical protein | | | |
| | | | SMU_1535 | *phsG* | glycogen phosphorylase |
| | | | SMU_1536 | *glgA* | glycogen synthase |
| | | | SMU_1537 | *glgD* | glycogen biosynthesis protein |
| | | | SMU_1538 | *glgC* | glucose-1-phosphate adenylyltransferase |
| | | | SMU_1539 | *glgB* | 1,4-alpha-glucan branching enzyme |
| SMU_1546 | | conserved hypothetical protein | | | |
| | | | SMU_1596 | *ptcC* | PTS system, cellobiose-specific IIC component |
| | | | SMU_1599 | *celR* | transcriptional regulator, required for cellobiose uptake and metabolism |
| | | | SMU_1600 | *ptcB* | PTS system IIB component, required for cellobiose uptake and metabolism |
| | | | SMU_1601 | *bgl* | 6-phospho-beta-glucosidase, required for cellobiose uptake and metabolism |
| SMU_1602 | | NAD(P)H-flavin oxidoreductase | | | |
| | | | SMU_1604c | | conserved hypothetical protein |
| SMU_1626 | *rl1* | 50S ribosomal protein L1 | | | |
| | | | SMU_1641c | | conserved hypothetical protein |
| SMU_1657c | | nitrogen regulatory protein PII | SMU_1657c | | nitrogen regulatory protein PII |
| SMU_1658 | *nrgA* | ammonium transporter, NrgA protein | SMU_1658 | *nrgA* | ammonium transporter, NrgA protein |
| SMU_1682c | | conserved hypothetical protein | | | |
| SMU_169 | | 50S ribosomal protein L13 | | | |
| SMU_1734 | *accA* | acetyl-CoA carboxylase alpha subunit | | | |
| SMU_1745c | | transcriptional regulator, MarR family | | | |
| SMU_1750c | | hypothetical protein | | | |
| SMU_1752c | | hypothetical protein | SMU_1752c | | hypothetical protein |
| SMU_1753c | | conserved hypothetical protein | | | |
| SMU_1754c | | conserved hypothetical protein | SMU_1754c | | conserved hypothetical protein |
| SMU_1755c | | conserved hypothetical protein | | | |
| SMU_1757c | | conserved hypothetical protein | SMU_1757c | | conserved hypothetical protein |
| SMU_1758c | | conserved hypothetical protein | | | |
| SMU_1760c | | conserved hypothetical protein | SMU_1760c | | conserved hypothetical protein |
| SMU_1761c | | conserved hypothetical protein | SMU_1761c | | conserved hypothetical protein |
| SMU_1762c | | conserved hypothetical protein | SMU_1762c | | conserved hypothetical protein |
| SMU_1763c | | conserved hypothetical protein | SMU_1763c | | conserved hypothetical protein |
| SMU_1764c | | conserved hypothetical protein | SMU_1764c | | conserved hypothetical protein |
| | | | SMU_1774c | | conserved hypothetical protein |
| | | | SMU_1775c | | hypothetical protein |
| | | | SMU_179 | | conserved hypothetical protein (possible oxidoreductase) |
| | | | SMU_180 | | oxidoreductase, possible fumarate reductase |
| SMU_1802c | | conserved hypothetical protein | | | |
| | | | SMU_182 | *sloA* | ABC transporter, ATP-binding protein, iron and/or manganese |
| | | | SMU_183 | *sloB* | manganese ABC transporter permease element |
| | | | SMU_184 | *sloC* | ABC transporter element, iron predicted binding protein |
| | | | SMU_185 | | hypothetical protein |
| SMU_1853 | | conserved hypothetical protein | | | |
| | | | SMU_1858 | *rs18* | 30S ribosomal protein S18 |
| | | | SMU_186 | *sloR* | metal-dependent transcriptional regulator (probable DtxR homolog) |
| | | | SMU_1860 | *rs6* | 30S ribosomal protein S6 |
| | | | SMU_1862 | | hypothetical protein |
| | | | SMU_1877 | *ptnA* | mannose PTS system component IIAB |
| | | | SMU_1878 | *ptnC* | mannose PTS system component IIC |
| | | | SMU_1879 | | mannose PTS system component IID |
| SMU_1889c | | hypothetical protein (possible relation to bacteriocin BlpU) | | | |
| | | | SMU_189 | | hypothetical protein |

| | | | | | |
|---|---|---|---|---|---|
| | | | SMU_1895c | | hypothetical protein |
| | | | SMU_1896c | | hypothetical protein |
| SMU_191c | | phage-related integrase | | | |
| SMU_193c | | conserved hypothetical protein | | | |
| | | | SMU_1941 | *atmB* | ABC transporter solute-binding protein |
| | | | SMU_1942c | | amino acid ABC transporter substrate-binding protein |
| SMU_194c | | conserved hypothetical protein, phage-related | | | |
| SMU_1945 | | conserved hypothetical protein | | | |
| | | | SMU_1955 | *groES* | co-chaperonin 10kDa |
| SMU_1956c | | conserved hypothetical protein | SMU_1956c | | conserved hypothetical protein |
| SMU_1957 | | fructose-specific Enzyme IID component | SMU_1957 | | fructose-specific Enzyme IID component |
| SMU_1958c | | fructose-specific Enzyme IIC component | SMU_1958c | | fructose-specific Enzyme IIC component |
| SMU_195c | | hypothetical protein | | | |
| SMU_1960c | | fructose-specific Enzyme IIB component | SMU_1960c | | fructose-specific Enzyme IIB component |
| SMU_1961c | | fructose-specific Enzyme IIA component | SMU_1961c | | fructose-specific Enzyme IIA component |
| SMU_196c | | immunogenic secreted protein (transfer protein) | | | |
| SMU_197c | | hypothetical protein | | | |
| SMU_198c | | conjugative transposon protein | | | |
| SMU_199c | | hypothetical protein | SMU_1988c | | probable DNA binding protein |
| SMU_200c | | hypothetical protein | | | |
| SMU_2002 | *rs11* | 30S ribosomal protein S11 | SMU_2002 | *rs11* | 30S ribosomal protein S11 |
| SMU_2010 | *rl18* | 50S ribosomal protein L18 | SMU_2010 | *rl18* | 50S ribosomal protein L18 |
| | | | SMU_2015 | *rl5* | 50S ribosomal protein L5 |
| SMU_2017 | *rl14* | 50S ribosomal protein L14 | SMU_2017 | *rl14* | 50S ribosomal protein L14 |
| SMU_2018 | *rs17* | 30S ribosomal protein S17 | SMU_2018 | *rs17* | 30S ribosomal protein S17 |
| SMU_201c | | conserved hypothetical protein | | | |
| | | | SMU_2020 | *rl16* | 50S ribosomal protein L16 |
| | | | SMU_2023c | | 30S ribosomal protein S19, C-terminal fragment |
| SMU_2027 | | transcriptional regulator/repressor | SMU_2027 | | transcriptional regulator/repressor |
| SMU_2028 | *sacB* | fructosyltransferase | | | |
| SMU_202c | | conserved hypothetical protein/Streptococcus-specific protein | SMU_202c | | conserved hypothetical protein/Streptococcus-specific protein |
| | | | SMU_2037 | *treA* | trehalose-6-phosphate hydrolase |
| | | | SMU_2038 | *pttB* | phosphotransferase system, trehalose-specific IIBC component (EIIBC-tre) |
| SMU_204c | | hypothetical protein | | | |
| SMU_205c | | conserved hypothetical protein | | | |
| SMU_206c | | hypothetical protein | | | |
| SMU_207c | | transcriptional regulator | | | |
| SMU_208c | | conserved hypothetical protein, FtsK/SpoIIIE family | | | |
| SMU_209c | | hypothetical protein | | | |
| SMU_210c | | hypothetical protein | | | |
| | | | SMU_2104a | | 50S ribosomal protein L32 |
| SMU_2117 | *opuCb* | glycine betaine / carnitine / choline ABC transporter permease | | | |
| SMU_211c | | hypothetical protein | | | |
| SMU_212c | | hypothetical protein | | | |
| | | | SMU_2127 | | succinic semialdehyde dehydrogenase |
| SMU_2146c | | conserved hypothetical protein | | | |
| SMU_2147c | | conserved hypothetical protein | SMU_2147c | | conserved hypothetical protein |
| SMU_215c | | hypothetical protein | | | |
| | | | SMU_2154c | | peptidase, M16 family |
| | | | SMU_217c | | conserved hypothetical protein; Streptococcus-specific protein |
| SMU_231 | *ilvB* | acetolactate synthase, large subunit (AHAS) | | | |
| | | | SMU_262 | *otcA* | ornithine carbamoyltransferase |
| | | | SMU_263 | | amino acid permease /putrescine antiporter |

| | | | | | |
|---|---|---|---|---|---|
| | | | SMU_267c | | glutamate--cysteine ligase (gamma ECS) |
| | | | SMU_27 | *acpP* | acyl carrier protein |
| SMU_272 | *ptxA* | PTS system, enzyme IIA component | | | |
| SMU_299c | | bacteriocin peptide precursor | SMU_299c | | bacteriocin peptide precursor |
| | | | SMU_30 | *purL* | phosphoribosylformylglycinamidine synthase |
| SMU_358 | | 30S ribosomal protein S7 | | | |
| SMU_361 | *pgk* | phosphoglycerate kinase | | | |
| SMU_367 | | Streptococcus-specific protein; similar to glucan-binding protein | | | |
| | | | SMU_378 | | hypothetical protein |
| SMU_40 | | conserved hypothetical protein | | | |
| | | | SMU_402 | *pfl* | pyruvate formate-lyase |
| SMU_412c | | Hit-like protein | | | |
| | | | SMU_43 | | conserved hypothetical protein |
| | | | SMU_438c | | (R)-2-hydroxyglutaryl-CoA dehydratase activator-related protein |
| | | | SMU_44 | | conserved hypothetical protein |
| | | | SMU_45 | | hypothetical protein |
| SMU_454 | *ftsL* | cell division protein | | | |
| | | | SMU_459 | | amino acid ABC transporter, substrate-binding protein |
| | | | SMU_460 | | amino acid ABC transporter, permease |
| | | | SMU_461 | | amino acid ABC transporter, ATP-binding protein |
| | | | SMU_48 | *purD* | phosphoribosylamine-glycine ligase |
| | | | SMU_491 | | transcriptional regulator, DeoR family |
| | | | SMU_494 | | transaldolase family protein |
| | | | SMU_495 | *gldA* | glycerol dehydrogenase |
| SMU_496 | *cysK* | cysteine synthetase A | SMU_496 | *cysK* | cysteine synthetase A |
| | | | SMU_50 | *purE* | phosphoribosylaminoimidazole carboxylase catalytic subunit |
| SMU_525 | | ABC transporter ATP-binding / permease protein | | | |
| | | | SMU_56 | | Streptococcus-specific protein |
| | | | SMU_576 | *lytR* | response regulator |
| | | | SMU_577 | *lytS* | sensor histidine kinase |
| SMU_586 | | conserved hypothetical protein, DevG family | | | |
| SMU_600c | | conserved hypothetical protein | | | |
| SMU_627 | | conserved hypothetical protein | | | |
| | | | SMU_628 | | DNA polymerase III, delta subunit |
| SMU_635 | | conserved hypothetical protein | SMU_635 | | conserved hypothetical protein |
| SMU_651c | | ABC transporter, periplasmic substrate-binding protein | | | |
| SMU_652c | | ABC transporter, ATP-binding protein (possible nitrate transport system) | | | |
| | | | SMU_675 | | phosphoenolpyruvate-protein phosphotransferase (enzyme I) |
| SMU_689 | | autolysin/endolysin | | | |
| SMU_698 | | 50S ribosomal protein L35 | SMU_698 | | 50S ribosomal protein L35 |
| | | | SMU_714 | | translation elongation factor Tu |
| | | | SMU_735 | | hypothetical protein |
| | | | SMU_768c | | conserved hypothetical protein |
| | | | SMU_769 | | conserved hypothetical protein |
| SMU_770c | | manganese transporter | SMU_770c | | manganese transporter/possible HitA ferric iron-binding periplasmic protein |
| | | | SMU_78 | *fruA* | fructan hydrolase; exo-beta-D-fructosidase |
| | | | SMU_79 | *fruB* | fructan hydrolase; exo-beta-D-fructosidase |
| | | | SMU_81 | *grpE* | co-chaperone protein GrpE |
| | | | SMU_82 | *dnaK* | chaperone protein, DnaK |
| SMU_84 | *truA* | tRNA pseudouridine synthase A | | | |
| SMU_867 | *rimM* | 16S rRNA processing protein | | | |
| | | | SMU_872 | | fructose-specific PTS system enzyme IIBC component |
| SMU_873 | *metE* | 5-methyltetrahydropteroyltriglutamate--homocysteine S-methyltransferase | | | |

| | | | | | |
|---|---|---|---|---|---|
| SMU_909 | malate permease | | | | |
| | | | SMU_930c | | transcriptional regulator |
| SMU_932 | conserved hypothetical protein | | SMU_932 | | conserved hypothetical protein |
| SMU_933 | amino acid ABC transporter, amino acid substrate-binding protein | | SMU_933 | | amino acid ABC transporter, amino acid substrate-binding protein |
| SMU_934 | amino acid ABC transporter, permease protein | | SMU_934 | | amino acid ABC transporter, permease protein |
| SMU_935 | amino acid ABC transporter, permease protein | | SMU_935 | | amino acid ABC transporter, permease protein |
| SMU_936 | amino acid ABC transporter, ATP-binding protein | | SMU_936 | | amino acid ABC transporter, ATP-binding protein |
| | | | SMU_940c | | hemolysin III-related protein |
| SMU_957 | 50S ribosomal protein L10 | | | | |
| SMU_959c | hypothetical protein | | | | |
| SMU_961 | macrophage infectivity potentiator-related protein | | | | |
| SMU_962 | acyl-CoA dehydrogenase | | | | |
| | | | SMU_980 | *bglP* | beta-glucoside-specific EII permease |
| | | | SMU_999 | | hypothetical protein |

**Supplemental Table 5.** DEGs in *Streptococcus oralis* 34 grown in coculture with *Streptococcus mutans* UA159 between TY- vs TY-Saliva**.** Red, upregulated genes. Blue, downregulated genes.

| Gene ID | Gene Name | Gene Description | Log2 Fold Change | Fold Change | P value | Log10 P value |
|---|---|---|---|---|---|---|
| KX728_RS08895 | | ABC transporter substrate-binding protein | 8.594658788 | 386.5895388 | 5.36E-135 | 134.27 |
| KX728_RS08905 | | sugar ABC transporter permease | 7.978349892 | 252.186964 | 2.36E-168 | 167.63 |
| KX728_RS08900 | | carbohydrate ABC transporter permease | 7.425799661 | 171.9445599 | 2.15E-141 | 140.67 |
| KX728_RS08885 | | bacteriocin immunity protein | 6.849969947 | 115.3576562 | 5.42E-64 | 63.27 |
| KX728_RS08890 | | CPBP family intramembrane metalloprotease | 5.638925535 | 49.82940819 | 3.35E-65 | 64.47 |
| KX728_RS06730 | | S8 family serine peptidase | 5.592911853 | 48.26521342 | 7.73E-18 | 17.11 |
| KX728_RS02155 | | MarR family transcriptional regulator | 3.734372888 | 13.30939316 | 7.97E-61 | 60.10 |
| KX728_RS02160 | | ABC transporter ATP-binding protein | 3.677840918 | 12.79795078 | 6.40E-52 | 51.19 |
| KX728_RS02165 | | ABC transporter permease/substrate-binding protein | 3.64125563 | 12.47748816 | 2.28E-43 | 42.64 |
| KX728_RS00335 | | VOC family protein | 3.517143232 | 11.44894877 | 4.17E-35 | 34.38 |
| KX728_RS04325 | | VIT family protein | 3.318274812 | 9.974709404 | 8.10E-38 | 37.09 |
| KX728_RS06775 | vex2 | ABC transporter ATP-binding subunit Vex2 | 3.21408834 | 9.279765444 | 1.19E-39 | 38.92 |
| KX728_RS07065 | | ferric reductase-like transmembrane domain-containing protein | 3.133894995 | 8.778016576 | 1.13E-27 | 26.95 |
| KX728_RS08940 | | alpha-L-fucosidase | 2.921151292 | 7.57450333 | 5.10E-37 | 36.29 |
| KX728_RS06770 | vex3 | ABC transporter permease subunit Vex3 | 2.780001048 | 6.86852848 | 9.15E-40 | 39.04 |
| KX728_RS06860 | | DNA starvation/stationary phase protection protein | 2.770436653 | 6.823143948 | 8.72E-43 | 42.06 |
| KX728_RS02500 | | DUF4649 family protein | 2.748995886 | 6.722490847 | 5.52E-36 | 35.26 |
| KX728_RS02495 | trxA | thioredoxin | 2.740237763 | 6.681804459 | 8.12E-38 | 37.09 |
| KX728_RS06780 | | ABC transporter permease | 2.662453987 | 6.331090364 | 5.64E-38 | 37.25 |
| KX728_RS03350 | | LPXTG-anchored adhesin/beta-galactosidase BgaA | 2.479929477 | 5.578701957 | 4.08E-32 | 31.39 |
| KX728_RS08220 | | glycoside hydrolase family 1 protein | 2.454596315 | 5.481597181 | 4.69E-31 | 30.33 |
| KX728_RS00240 | | GNAT family N-acetyltransferase | 2.192728694 | 4.57169353 | 8.16E-08 | 7.09 |
| KX728_RS00225 | purF | amidophosphoribosyltransferase | 2.150928601 | 4.441135538 | 1.49E-07 | 6.83 |
| KX728_RS05075 | | phosphate-transport permease PitB | 2.060081583 | 4.170098851 | 1.54E-17 | 16.81 |
| KX728_RS00235 | purN | phosphoribosylglycinamide formyltransferase | 2.029800409 | 4.083483529 | 4.98E-07 | 6.30 |
| KX728_RS05085 | pitA | DUF5979 domain-containing protein | 2.018806003 | 4.052482633 | 4.52E-23 | 22.35 |
| KX728_RS00255 | purE | 5-(carboxyamino)imidazole ribonucleotide mutase | 1.985205672 | 3.959191012 | 9.10935E-06 | 5.04 |
| KX728_RS03030 | | NAD(P)H-dependent oxidoreductase | 1.965554425 | 3.90562768 | 2.93E-24 | 23.53 |
| KX728_RS08205 | | hypothetical protein | 1.943849719 | 3.847309034 | 2.02E-10 | 9.70 |
| KX728_RS00230 | purM | phosphoribosylformylglycinamidine cyclo-ligase | 1.873513636 | 3.664239076 | 1.15911E-05 | 4.94 |
| KX728_RS02505 | | MarR family transcriptional regulator | 1.783269439 | 3.442053303 | 1.36E-19 | 18.87 |
| KX728_RS01665 | | glycoside hydrolase N-terminal domain-containing protein | 1.752294577 | 3.368939631 | 1.38E-15 | 14.86 |
| KX728_RS00260 | purK | 5-(carboxyamino)imidazole ribonucleotide synthase | 1.752197519 | 3.368712992 | 1.91126E-06 | 5.72 |
| KX728_RS00245 | purH | bifunctional phosphoribosylaminoimidazolecarboxamide formyltransferase | 1.736370218 | 3.331958005 | 7.27E-07 | 6.14 |
| KX728_RS05080 | sipA | PI-2 pilus system signal peptidase SipA | 1.720812896 | 3.296220824 | 3.49E-20 | 19.46 |
| KX728_RS05065 | srtB | class B sortase | 1.697254888 | 3.242833357 | 4.76E-18 | 17.32 |
| KX728_RS05070 | srtB | class B sortase | 1.681768865 | 3.208210636 | 5.19E-17 | 16.28 |
| KX728_RS05095 | | pneumococcal-type histidine triad protein | 1.588943953 | 3.008290631 | 6.21E-13 | 12.21 |
| KX728_RS05105 | | pneumococcal-type histidine triad protein | 1.588470527 | 3.007303611 | 1.24E-09 | 8.91 |
| KX728_RS06555 | manA | mannose-6-phosphate isomerase%2C class I | 1.565109061 | 2.958998696 | 3.01E-20 | 19.52 |
| KX728_RS00285 | | LPXTG-anchored beta-N-acetylhexosaminidase StrH | 1.560396091 | 2.949348065 | 1.63E-11 | 10.79 |
| KX728_RS02370 | trpA | tryptophan synthase subunit alpha | 1.543939419 | 2.915896311 | 1.46E-08 | 7.83 |
| KX728_RS04430 | | hypothetical protein | 1.496376219 | 2.821331558 | 3.72635E-06 | 5.43 |
| KX728_RS02290 | | xanthine phosphoribosyltransferase | 1.489493378 | 2.807903545 | 3.01E-08 | 7.52 |
| KX728_RS05100 | | zinc ABC transporter substrate-binding protein | 1.458073221 | 2.747411903 | 5.59E-11 | 10.25 |
| KX728_RS07210 | | hypothetical protein | 1.435388383 | 2.704549651 | 6.29E-09 | 8.20 |

| | | | | | | |
|---|---|---|---|---|---|---|
| KX728_RS06750 | | hypothetical protein | 1.429062367 | 2.692716541 | 7.64E-08 | 7.12 |
| KX728_RS02815 | | G5 domain-containing protein | 1.384204006 | 2.610278979 | 2.33E-08 | 7.63 |
| KX728_RS06350 | | DUF6161 domain-containing protein | 1.383948184 | 2.609816159 | 3.3961E-06 | 5.47 |
| KX728_RS00500 | ilvD | dihydroxy-acid dehydratase | 1.373954783 | 2.591800698 | 8.05E-12 | 11.09 |
| KX728_RS05365 | | CHY zinc finger protein | 1.369157272 | 2.583196287 | 2.93E-15 | 14.53 |
| KX728_RS09230 | | TetR/AcrR family transcriptional regulator | 1.319536575 | 2.495859245 | 1.54345E-05 | 4.81 |
| KX728_RS07205 | | hypothetical protein | 1.319109513 | 2.495120537 | 3.66774E-06 | 5.44 |
| KX728_RS08910 | | beta-N-acetylhexosaminidase | 1.313168566 | 2.484866881 | 1.28E-08 | 7.89 |
| KX728_RS09040 | | LPXTG cell wall anchor domain-containing protein | 1.308979163 | 2.47766161 | 1.51E-12 | 11.82 |
| KX728_RS07245 | | ROK family protein | 1.294425187 | 2.452792494 | 6.33E-12 | 11.20 |
| KX728_RS07235 | | type I restriction endonuclease subunit R | 1.293345305 | 2.450957223 | 1.25E-16 | 15.90 |
| KX728_RS02350 | trpD | anthranilate phosphoribosyltransferase | 1.259318469 | 2.393826296 | 4.20769E-06 | 5.38 |
| KX728_RS08200 | | hypothetical protein | 1.255137641 | 2.386899199 | 6.00457E-06 | 5.22 |
| KX728_RS04535 | | thioredoxin | 1.245197417 | 2.370509898 | 3.26E-17 | 16.49 |
| KX728_RS02285 | | purine permease | 1.240521913 | 2.362839956 | 3.74E-07 | 6.43 |
| KX728_RS08920 | | alpha-mannosidase | 1.233427494 | 2.351249267 | 1.79E-07 | 6.75 |
| KX728_RS07120 | | bifunctional 2'%2C3'-cyclic-nucleotide 2'-phosphodiesterase/3'-nucleotidase | 1.217849224 | 2.32599698 | 3.11964E-06 | 5.51 |
| KX728_RS05370 | | U32 family peptidase | 1.206423916 | 2.307649181 | 4.85E-13 | 12.31 |
| KX728_RS02340 | trpE | anthranilate synthase component I | 1.199571909 | 2.296715104 | 9.46704E-06 | 5.02 |
| KX728_RS05155 | | transcription antiterminator | 1.162716837 | 2.23878632 | 6.04529E-06 | 5.22 |
| KX728_RS01405 | | hypothetical protein | 1.150402986 | 2.219758899 | 1.11293E-06 | 5.95 |
| KX728_RS07240 | | TVP38/TMEM64 family protein | 1.14273449 | 2.207991304 | 5.84E-12 | 11.23 |
| KX728_RS02280 | | endonuclease | 1.126964283 | 2.183987021 | 6.86412E-05 | 4.16 |
| KX728_RS08915 | | ROK family protein | 1.125295726 | 2.181462579 | 2.87319E-05 | 4.54 |
| KX728_RS07075 | | hypothetical protein | 1.109478649 | 2.157676607 | 2.46E-12 | 11.61 |
| KX728_RS00310 | | PTS system mannose/fructose/sorbose family transporter subunit IID | 1.077137795 | 2.109846141 | 7.42E-10 | 9.13 |
| KX728_RS05020 | | 4-oxalocrotonate tautomerase | 1.073160208 | 2.104037194 | 2.90E-07 | 6.54 |
| KX728_RS05195 | | PTS sugar transporter subunit IIA | 1.058248734 | 2.08240219 | 4.84E-08 | 7.32 |
| KX728_RS04125 | | DEAD/DEAH box helicase | 1.051245111 | 2.072317579 | 7.72256E-05 | 4.11 |
| KX728_RS00315 | | PTS sugar transporter subunit IIA | 1.048521943 | 2.068409654 | 2.55E-09 | 8.59 |
| KX728_RS02400 | | Rrf2 family transcriptional regulator | 1.047590392 | 2.067074509 | 1.15E-07 | 6.94 |
| KX728_RS08325 | | ACT domain-containing protein | 1.045165004 | 2.063602363 | 2.3933E-05 | 4.62 |
| KX728_RS03740 | | AAA family ATPase | 1.038749521 | 2.054446157 | 5.57E-07 | 6.25 |
| KX728_RS05060 | hemH | ferrochelatase | 1.027872537 | 2.03901521 | 5.55E-11 | 10.26 |
| KX728_RS07730 | | beta-N-acetylglucosaminidase domain-containing protein | 1.024537102 | 2.034306559 | 9.43514E-06 | 5.03 |
| KX728_RS08805 | | ABC transporter substrate-binding protein | -1.015249382 | -2.021252253 | 1.07773E-05 | 4.97 |
| KX728_RS04055 | | cysteine desulfurase | -1.0231731 | -2.032384125 | 3.49E-11 | 10.46 |
| KX728_RS06080 | | EamA family transporter | -1.050139232 | -2.070729681 | 1.53E-12 | 11.81 |
| KX728_RS02470 | | aquaporin | -1.052375609 | -2.073942088 | 1.03E-08 | 7.99 |
| KX728_RS08705 | | M20/M25/M40 family metallo-hydrolase | -1.065773519 | -2.093291924 | 1.81E-14 | 13.74 |
| KX728_RS08800 | | argininosuccinate synthase | -1.076360093 | -2.10870911 | 2.41E-07 | 6.62 |
| KX728_RS03410 | | CshA/CshB family fibrillar adhesin-related protein | -1.099059404 | -2.142149849 | 3.78E-13 | 12.42 |
| KX728_RS09005 | | DUF1275 family protein | -1.154040126 | -2.225362135 | 1.08E-09 | 8.97 |
| KX728_RS03595 | | alpha-amylase | -1.160287131 | -2.235019055 | 3.18E-11 | 10.50 |
| KX728_RS04355 | | ABC transporter substrate-binding protein/permease | -1.209381243 | -2.312384398 | 1.08E-12 | 11.97 |
| KX728_RS08210 | glmS | glutamine--fructose-6-phosphate transaminase (isomerizing) | -1.254474317 | -2.385802 | 3.21E-10 | 9.49 |
| KX728_RS06240 | pdxT | pyridoxal 5'-phosphate synthase glutaminase subunit PdxT | -1.271839986 | -2.414693347 | 2.36481E-06 | 5.63 |
| KX728_RS09130 | cysK | cysteine synthase A | -1.305594729 | -2.471856049 | 1.13E-07 | 6.95 |
| KX728_RS03360 | | Cna B-type domain-containing protein | -1.311886196 | -2.482659136 | 2.97E-10 | 9.53 |
| KX728_RS04350 | | amino acid ABC transporter ATP-binding protein | -1.34378575 | -2.538164809 | 6.77E-14 | 13.17 |
| KX728_RS06245 | pdxS | pyridoxal 5'-phosphate synthase lyase subunit PdxS | -1.354340111 | -2.556801413 | 2.06E-07 | 6.69 |

| | | | | | | |
|---|---|---|---|---|---|---|
| KX728_RS06980 | | amino acid ABC transporter ATP-binding protein | -1.385399359 | -2.612442637 | 1.97E-08 | 7.71 |
| KX728_RS01540 | | hypothetical protein | -1.407811474 | -2.65334353 | 1.41E-10 | 9.85 |
| KX728_RS07830 | | peptide ABC transporter substrate-binding protein | -1.462539261 | -2.755930031 | 3.75E-12 | 11.43 |
| KX728_RS04630 | | bifunctional glycosyltransferase family 2/GtrA family protein | -1.468265608 | -2.76689061 | 6.86921E-05 | 4.16 |
| KX728_RS00575 | malQ | 4-alpha-glucanotransferase | -1.46830233 | -2.76696104 | 1.64E-16 | 15.78 |
| KX728_RS01545 | | DUF5960 family protein | -1.473373239 | -2.776703705 | 8.20E-15 | 14.09 |
| KX728_RS04645 | | peptidase | -1.475744817 | -2.78127195 | 5.70922E-06 | 5.24 |
| KX728_RS06985 | | transporter substrate-binding domain-containing protein | -1.547519309 | -2.923140774 | 3.76E-10 | 9.43 |
| KX728_RS06995 | | amino acid ABC transporter permease | -1.589534921 | -3.009523163 | 1.95E-08 | 7.71 |
| KX728_RS07995 | | hypothetical protein | -1.67538197 | -3.194039092 | 9.21204E-05 | 4.04 |
| KX728_RS06990 | | amino acid ABC transporter permease | -1.678781217 | -3.201573692 | 2.50E-10 | 9.60 |
| KX728_RS00655 | | gamma-glutamyl-gamma-aminobutyrate hydrolase family protein | -1.681462541 | -3.207529515 | 1.48E-16 | 15.83 |
| KX728_RS04635 | | phosphodiester glycosidase family protein | -1.791304326 | -3.461276816 | 1.1182E-05 | 4.95 |
| KX728_RS07585 | | acyl-CoA dehydrogenase family protein | -1.794512366 | -3.468982027 | 1.37E-11 | 10.86 |
| KX728_RS01530 | | XRE family transcriptional regulator | -1.831952874 | -3.560186636 | 3.43E-16 | 15.47 |
| KX728_RS01730 | | iron chelate uptake ABC transporter family permease subunit | -1.835180204 | -3.56815974 | 3.51E-08 | 7.46 |
| KX728_RS07100 | tpx | thiol peroxidase | -1.875027437 | -3.668085933 | 2.23E-24 | 23.65 |
| KX728_RS01535 | | Y-family DNA polymerase | -1.899969694 | -3.732053567 | 1.98E-23 | 22.70 |
| KX728_RS07590 | | carboxymuconolactone decarboxylase family protein | -1.925056157 | -3.797516299 | 1.56E-11 | 10.81 |
| KX728_RS02855 | | ABC transporter substrate-binding protein | -2.004799632 | -4.013329566 | 1.08E-10 | 9.97 |
| KX728_RS02475 | queF | preQ(1) synthase | -2.099964536 | -4.286988468 | 3.33E-16 | 15.48 |
| KX728_RS02860 | | alpha/beta hydrolase-fold protein | -2.141639219 | -4.412631335 | 2.45E-17 | 16.61 |
| KX728_RS02850 | | ABC transporter ATP-binding protein | -2.160880975 | -4.471878452 | 6.16E-13 | 12.21 |
| KX728_RS02845 | | iron chelate uptake ABC transporter family permease subunit | -2.275281952 | -4.840922348 | 4.65E-13 | 12.33 |
| KX728_RS03850 | | alpha/beta hydrolase | -2.322085996 | -5.000547272 | 7.96E-22 | 21.10 |
| KX728_RS01725 | | ATP-binding cassette domain-containing protein | -2.420706113 | -5.354330203 | 2.77E-10 | 9.56 |
| KX728_RS02230 | | cation diffusion facilitator family transporter | -2.472033778 | -5.548253775 | 6.88E-42 | 41.16 |
| KX728_RS08145 | | transporter substrate-binding domain-containing protein | -2.486984516 | -5.606049629 | 2.16E-14 | 13.67 |
| KX728_RS08135 | | amino acid ABC transporter permease | -2.512371045 | -5.70557009 | 2.16E-14 | 13.67 |
| KX728_RS08130 | | amino acid ABC transporter ATP-binding protein | -2.527172942 | -5.764409956 | 2.70E-15 | 14.57 |
| KX728_RS02490 | queC | 7-cyano-7-deazaguanine synthase QueC | -2.541642165 | -5.822513844 | 2.59E-28 | 27.59 |
| KX728_RS02485 | queD | 6-carboxytetrahydropterin synthase QueD | -2.544797935 | -5.835264028 | 1.27E-35 | 34.90 |
| KX728_RS02480 | queE | 7-carboxy-7-deazaguanine synthase QueE | -2.571590225 | -5.944643215 | 5.78E-28 | 27.24 |
| KX728_RS08140 | | amino acid ABC transporter permease | -2.589941128 | -6.020741296 | 3.32E-15 | 14.48 |
| KX728_RS08880 | | rhodanese-related sulfurtransferase | -2.598490839 | -6.056527396 | 5.53E-33 | 32.26 |
| KX728_RS01720 | | siderophore ABC transporter substrate-binding protein | -2.60398491 | -6.079635836 | 2.95E-17 | 16.53 |
| KX728_RS08150 | | uroporphyrinogen decarboxylase family protein | -2.652419631 | -6.287208604 | 2.69E-16 | 15.57 |
| KX728_RS08155 | | hypothetical protein | -2.822062082 | -7.071724551 | 2.36E-20 | 19.63 |
| KX728_RS08160 | | uroporphyrinogen decarboxylase | -2.858915191 | -7.254696149 | 6.29E-17 | 16.20 |

**Supplemental Table 6.** *Streptococcus oralis* 34 DEGs during coculture with *Streptococcus mutans* UA159 in either TY-, TY-Saliva, or both. Light grey,TY- only; Red, TY-Saliva; Dark blue, both.

| TY | | | | TY-Saliva | | |
|---|---|---|---|---|---|---|
| **Gene ID** | **Gene** | **Product** | | **Geneid** | **Gene** | **Product** |
| | | | (red) | KX728_RS00335 | | VOC family protein |
| | | | (red) | KX728_RS00735 | tRNA-Tyr | |
| | | | (red) | KX728_RS00900 | adhE | bifunctional acetaldehyde-CoA/alcohol dehydrogenase |
| KX728_RS02040 | tRNA-Lys | | (light grey) | | | |
| KX728_RS02275 | | hypothetical protein | (light grey) | | | |
| KX728_RS02565 | | hypothetical protein | (light grey) | | | |
| KX728_RS05075 | sipA | Signal peptidase I (EC 3.4.21.89) | (light grey) | | | |
| KX728_RS05780 | | Late competence protein ComEA, DNA receptor | (light grey) | | | |
| KX728_RS06240 | pdxS | Pyridoxal 5'-phosphate synthase (glutamine hydrolyzing) | (light grey) | | | |
| KX728_RS06245 | | NADH peroxidase (EC 1.11.1.1) | (light grey) | | | |
| KX728_RS06645 | pyrF | Orotidine 5'-phosphate decarboxylase (EC 4.1.1.23) | (light grey) | | | |
| KX728_RS06650 | | Cell division protein FtsQ | (light grey) | | | |
| KX728_RS06750 | | Fructose-bisphosphate aldolase class II (EC 4.1.2.13) | (light grey) | | | |
| KX728_RS06770 | vex2 | ABC transporter, ATP-binding protein Vex2 | (light grey) | | | |
| KX728_RS06780 | tRNA-Leu | | (light grey) | | | |

**Supplemental Table 7.** Bacterial strains used in this study.

| Species | Strain | Genotype or description | Antibiotic resistance | Reference or source |
|---|---|---|---|---|
| *Actinomyces oris* | WVU627 | | | Lab Stock |
| *Corynebacterium matruchotii* | ATCC 14266 | | | Lab Stock |
| *Lactobacilllus casei* | ATCC 4646 | | | Lab Stock |
| *Streptococcus cristatus* | ATCC 51100 | | | Lab Stock |
| *Streptococcus gordonii* | DL1 | | | Lab Stock |
| *Streptococcus mits* | ATCC 49456 | | | Lab Stock |
| *Streptococcus mutans* | UA159 | | | Lab stock |
| | pMZ- / UA159 | UA159 with pMZ plasmid integrated into genome. Serves as GFP(-) background control strain during fluorescent intensity-based competitive growth assays and as a "marked" *S. mutans* strain during competitive index assays | Kanamycin | Sheilds et al. 2019, Appl Environ Microbiol. |
| | pMZ-P*veg*::*gfp* / UA159 | UA159 with pMZ plasmid integrated into genome harboring constitutively-active *gfp*. Serves as GFP(+) strain during fluorescent intensity-based competitive growth assays and used in coculture microscopy experiments | Kanamycin | Sheilds et al. 2019, Appl Environ Microbiol. |
| *Streptococcus oralis* | 34 | | | Lab Stock |
| | ΔKX728_03760 | Mutation of gene KX728_03760 which is not actively expressed in *S. oralis* 34 by RNA-Seq. This strain is used within competitive index assays as a "marked" *S. oralis* strain | Spectinomycin | Lab Stock |
| *Streptococcus sanguinis* | SK36 | | | Lab Stock |
| *Streptococcus sobrinus* | 6715 | | | Lab Stock |

**Supplemental Table 8.** Catalog numbers of growth medium additives.

| Category | Item | Vendor | Catalog Number |
|---|---|---|---|
| Carbohydrates | D-(+)-Cellobiose | Sigma-Aldrich | 22150 |
| | D-(-)-Fructose | Sigma-Aldrich | F0127 |
| | D-(+)-Galactose | Sigma-Aldrich | G0750 |
| | N-Acetyl-D-Glucosamine | Sigma-Aldrich | A8625 |
| | D-(+)-Glucose | Sigma-Aldrich | G8270 |
| | Lactose Monohydrate | Sigma-Aldrich | 1.07657 |
| | D-(+)-Mannose | Sigma-Aldrich | 63580 |
| Metals | Iron(II) Sulfate Heptahydrate | Sigma-Aldrich | F8633 |
| | Manganese(II) Sulfate Heptahydrate | Sigma-Aldrich | 229784 |
| | Zinc Sulfate Heptahydrate | Sigma-Aldrich | 20251 |

**Supplemental Table 9.** Concentration* of inoculated bacterial strains during competitive competitions.
* Concentrations were determined based on the growth rate of individual species, such that the initial growth rate and entrance into exponential growth phase is matched to that of *Streptococcus mutans* pMZ-P*veg*::*gfp* / UA159 strain.

| Strain Name | Concentration of Competitor (in comparision to *S. mutans*) | *S. mutans* Concentration |
|---|---|---|
| *S.* sp. A12 | 0.5x | 1x |
| *S. cristatus* ATCC 51100 | 10x | 1x |
| *S. gordonii* DL1 | 10% solution (1:10 dilution); 1x | 1x |
| *S. mitis* ATCC 49456 | 1x | 1x |
| *S. oralis* 34 | 1x | 1x |
| *S. sanguinis* SK36 | 1x | 1x |
| *S. sobrinus* 6715 | 2x | 1x |

**Supplemental Table 10.** Species identity of human saliva-derived bacterial isolates used in this study.

| Strain Identifier | Species Identified by 16S Sequencing |
|---|---|
| SOSUI001 | *Streptococcus salivarius* |
| SOSUI002 | *Streptococcus salivarius* |
| SOSUI003 | *Rothia mucilaginosa* |
| SOSUI004 | *Rothia mucilaginosa* |
| SOSUI005 | *Streptococcus infantis* |
| SOSUI006 | *Streptococcus parasanguinis* |
| SOSUI007 | *Rothia mucilaginosa* |
| SOSUI008 | *Streptococcus salivarius* |
| SOSUI009 | *Streptococcus parasanguinis* |
| SOSUI010 | *Streptococcus salivarius* |
| SOSUI011 | *Neisseria mucosa* |
| SOSUI012 | *Streptococcus mitis* |
| SOSUI013 | *Staphylococcus epidermidis* |
| SOSUI014 | N/A |
| SOSUI015 | *Streptococcus pseudopneumonia* |
| SOSUI016 | *Granulicatella elegans* |
| SOSUI017 | *Streptococcus salivarius* |
| SOSUI018 | *Rothia dentocariosa* |
| SOSUI019 | *Rothia dentocariosa* |
| SOSUI020 | *Actinomyces oris* |
| SOSUI021 | *Streptococcus mitis* |
| SOSUI022 | *Actinomyces oris* |
| SOSUI023 | N/A |
| SOSUI024 | *Neisseria perflava* |
| SOSUI025 | *Streptococcus sanguinis* |
| SOSUI026 | *Actinomyces oris* |
| SOSUI027 | *Streptococcus mitis* |
| SOSUI028 | *Rothia dentocariosa* |
| SOSUI029 | *Streptococcus mitis* |
| SOSUI030 | *Streptococcus mitis* |
| SOSUI031 | *Streptococcus mitis* |
| SOSUI032 | *Streptococcus mitis* |
| SOSUI033 | *Streptococcus infantis* |
| SOSUI034 | *Streptococcus mitis* |
| SOSUI035 | *Streptococcus sanguinis* |
| SOSUI036 | *Streptococcus sanguinis* |
| SOSUI037 | *Streptococcus mitis* |

N/A = identity not available at the time of publication

**Supplemental Table 11.** Genome files used in this study.

| Species | Isolate | Accession | Reference? | Source |
|---|---|---|---|---|
| *Streptococcus mutans* | UA159 | NC_004350.2 | Yes | NCBI GenBank |
| *Streptococcus oralis* | 34 | CP079724.1 | No | NCBI GenBank |
